# AML/T cell interactomics uncover correlates of patient outcomes and the key role of ICAM1 in T cell killing of AML

**DOI:** 10.1101/2023.09.21.558911

**Authors:** Ece Canan Sayitoglu, Bogdan A. Luca, Allison P. Boss, Benjamin Craig Thomas, Robert Arthur Freeborn, Molly J. Uyeda, Pauline P. Chen, Yusuke Nakauchi, Colin Waichler, Norman Lacayo, Rosa Bacchetta, Ravindra Majeti, Andrew J. Gentles, Alma-Martina Cepika, Maria Grazia Roncarolo

## Abstract

T cells are important for the control of acute myeloid leukemia (AML), a common and often deadly malignancy. We observed that some AML patient samples are resistant to killing by human engineered cytotoxic CD4^+^ T cells. Single-cell RNA-seq of primary AML samples and CD4^+^ T cells before and after their interaction uncovered transcriptional programs that correlate with AML sensitivity or resistance to CD4^+^ T cell killing. Resistance-associated AML programs were enriched in AML patients with poor survival, and killing-resistant AML cells did not engage T cells *in vitro*. Killing-sensitive AML potently activated T cells before being killed, and upregulated *ICAM1*, a key component of the immune synapse with T cells. Without ICAM1, killing-sensitive AML became resistant to killing to primary *ex vivo*-isolated CD8^+^ T cells *in vitro*, and engineered CD4^+^ T cells *in vitro* and *in vivo*. Thus, ICAM1 on AML acts as an immune trigger, allowing T cell killing, and could affect AML patient survival *in vivo*.

**SIGNIFICANCE:** AML is a common leukemia with sub-optimal outcomes. We show that AML transcriptional programs correlate with susceptibility to T cell killing. Killing resistance-associated AML programs are enriched in patients with poor survival. Killing-sensitive, but not resistant AML activate T cells and upregulate *ICAM1* that binds to LFA-1 on T cells, allowing immune synapse formation which is critical for AML elimination.

**GRAPHICAL ABSTRACT:** 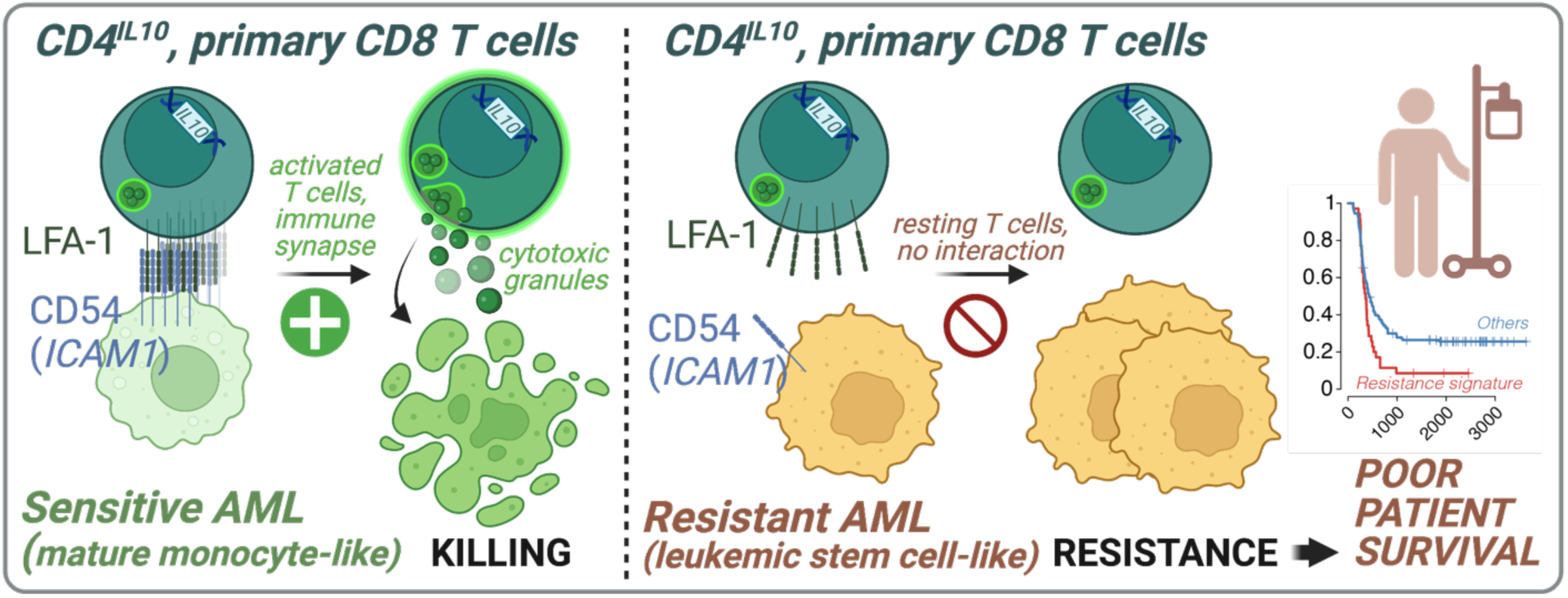

## INTRODUCTION

Acute myeloid leukemia (AML) affects ∼20,000 new patients annually in the US. Many patients experience relapse after standard-of-care therapy^1^, which can lead to poor outcomes. Indeed, adult and pediatric AML patients have only ∼30% and ∼70% overall survival (OS), respectively^2,3^. T cells play an important role in anti-tumor immunity and long-term survival of AML patients. Elevated frequency of T cells in AML patient bone marrow correlates with increased survival^4^. Peripheral T cells in AML patients can exhibit dysregulated gene expression profiles, showing signs of activation but defects in immune synapse formation^5^, which is required for tumor cell killing. T cell exhaustion and reduced CD4^+^ T helper (Th) cell activity have also been observed in patients with AML^6^.

CD8^+^ T cells have potent cytotoxic activity and are considered critical for tumor control^7^, but AML can also cause CD8^+^ T cell dysfunction^8,9^. While CD8^+^ T cells recognize AML antigens via HLA class I, relapse in up to 40% of AML patients is driven by the loss of HLA class II expression on AML cells^10,11^. Expression of HLA class II is needed for the presentation of AML antigens to CD4^+^ T cells. Antigen-activated CD4^+^ T cells secrete IL-2, IFN-γ and TNF-α that can activate CD8^+^ T cells, helping them to kill tumor cells^12^. Further, IFN-γ secreted by CD4^+^ T cells can restore HLA class II expression on AML^10,11^. Finally, CD4^+^ T cells can also acquire cytotoxic features themselves, and kill the tumor cells directly^12^. Thus, CD4^+^ T cells have an important role in the immune response to AML cells. Yet, the interaction between AML cells and CD4^+^ T cells, and its effect on AML patient survival, remains poorly understood.

Since primary human cytotoxic CD4^+^ T cells are rare, comprising on average 2.2% of the CD4^+^ T cell compartment in healthy donors^13^, we opted to study AML-CD4^+^ T cell interaction using engineered human cytotoxic CD4^+^ T cells called CD4^IL10^. CD4^IL10^ cells are generated by lentiviral overexpression of the human *IL10* gene in healthy donor CD4^+^ T cells^14–17^. Constitutive expression of IL-10 in CD4^+^ T cells has dual benefits: it can re-invigorate exhausted T cells^18^, and induces cytotoxic phenotype and functions^19^. Indeed, CD4^IL10^ cells kill AML cell lines and primary AML patient samples *in vitro* via perforin and granzyme B^14–16,20^, and inhibit leukemia progression *in vivo*^15^. Notably, CD4^IL10^ cells kill AML cells in an antigen-independent manner, perhaps due to tonically active T cell receptor (TCR) after *in vitro* expansion^21^, bypassing the dependence on the very rare AML antigen-specific T cells^22^ to study their interaction with AML cells.

While most primary adult^15^ and pediatric AML cells are killed by CD4^IL10^ cells, some are resistant to killing^16^. Previously, we showed that CD200 glycoprotein on resistant AML impairs the release cytotoxic granules from CD4^IL10^ cells^16^, and others have described the same effect on CD8^+^ T cells^23^ and NK cells^24^. However, the inhibitory effect of CD200 on T cells was only partial, and not sufficient to completely explain the AML resistance to T cell killing. To better understand the determinants of AML sensitivity or resistance to T cell killing, we analyzed killing-sensitive and -resistant primary AML samples before and after co-culture with CD4^IL10^ cells using single-cell RNA sequencing (scRNA-seq). We identified AML transcriptional programs associated with sensitivity or resistance to CD4^IL10^ killing, showing that resistance-related programs are found in less differentiated, stem cell-like AML cells, and in patients with poor survival rates. We also revealed that killing-sensitive, but not killing-resistant AML, successfully activate CD4^IL10^ cells. We demonstrated that AML expression of *ICAM1*, which is required for the immune synapse formation with T cells, is necessary for AML killing by both CD4^IL10^ cells and primary, *ex vivo*-isolated CD8^+^ T cells. These results emphasize the role of T cell response to AML in AML patient survival, and identify ICAM1 as one of the immune triggers necessary for AML killing by T cells.

## METHODS

### Primary human cells, cell lines, and culture

De-identified pediatric AML bone marrow aspirates were obtained from the Children’s Oncology Group (COG; CA, USA; protocol #AAML-18B2-Q, **Table S1**). Each AML sample contained ≥ 80% blasts, as reported by COG (**Table S1**). Primary adult AML samples SU540 and SU555 were obtained with the help of Dr. Ravindra Majeti and Dr. Yusuke Nakauchi (Stanford University Medical Center Institutional Review Board protocol # 6453). Human blood leukocytes were obtained from de-identified healthy donors (Stanford Blood Center, CA, USA), from which the CD4^IL10^ cells were generated, cultured, and functionally tested as described^16,17^. Human CD8^+^ T cells were isolated from healthy donors via magnetic bead labeling (CD8^+^ T cell isolation kit, Miltenyi Biotec, Germany). U937 (ATCC, VA, USA), K562 (ATCC, VA, USA) and primary AML cells were cultured as before^16^.

### Killing and degranulation assays

CD4^IL10^ cells were co-cultured with target cells at a 1:1 effector: target (E : T) ratio for 3 days as described^16^, unless otherwise specified. When indicated, LFA-1 inhibitors (BIRT377, BMS688571 and A286982 from Tocris Bioscience, UK; BMS587101 from MedChemExpress, NJ, USA) were added into the killing assay at 0.1, 1 and 10 μM, and elimination efficiency was calculated 24 hr after co-culture. For killing assay with *ex vivo*-isolated primary CD8^+^ T cells, target U937 cells (killing-sensitive) and control K562 cells (killing-resistant) were pre-treated with soluble anti-CD3 monoclonal antibody (mAb; OKT3, Miltenyi Biotec, at 200 ng/ml) for 15 min at 37°C prior to co-culture with primary human CD8^+^ T cells, to allow the anti-CD3 mAb to bind to Fc-receptors on U937 cells (control K652 cells do not express Fc receptors). Next, freshly isolated human CD8^+^ T cells are added to pre-treated target cells at a 1:1 E:T ratio, and co-cultured for 3 days in the presence of 50 IU/ml IL-2 and 50 ng/ml final concentration of anti-CD3 mAb. After culture, remaining cells in each condition were quantified via CountBeads using flow cytometry as in the CD4^IL10^ cell killing assay^16^. Killing was calculated as elimination efficiency (%): [1 – (AML cell number cultured with T cells) / (AML cell number cultured alone)] x 100^16^. For CD8^+^ T cell degranulation assay, target cells were pre-treated with anti-CD3 mAb as above, and co-cultured with human CD8^+^ T cells at 1:1 E:T ratio in the presence of anti-CD107a-APC mAb (BD Biosciences, CA, USA). After 1 hour, brefeldin A and monensin (BioLegend, CA, USA; used as per manufacturer’s instructions) were added to each well, cells were cultured for 4 more hours, stained with a viability dye and anti-CD3 and anti-CD8 mAb (**Table S2**), and fixed with 4% formaldehyde until acquisition by flow cytometry.

### Flow cytometry

Data was acquired on BD LSR Fortessa and BD FACS Aria II (BD, NJ, USA), and analyzed using FlowJo 10.8 (BD, NJ, USA). Antibodies are listed in **Table S2**.

### CRISPR/Cas9-mediated genome editing

CRISPR/Cas9 knock-outs of *IFNGR1, TNFRSF1B, ITGB2* were performed as previously^25^, with minor optimizations in sgRNA concentration and electroporation (**Supplemental Methods**). *ICAM1* knockout was made by first cloning an *ICAM1*-targeting sgRNA into a modified lentiCRISPRv2 plasmid with an RFP reporter instead of puromycin (RFP subclone kindly provided by Prof. Ravindra Majeti; original clone was a gift from Feng Zhang, Addgene plasmid #52961). Third generation lentivirus was produced as described^16^. RFP^+^ICAM1^-^ cells were FACS-sorted one week post transduction. Knockout was confirmed by flow cytometry or Sanger sequencing ≥ 5 days after editing.

### scRNA-seq

We thawed four cryopreserved primary AML samples with pre-determined susceptibility to CD4^IL10^ killing, rested them for 2 hours at 37°C, then FACS-sorted an aliquot of live cells for scRNA-seq samples on day 0. CD4^IL10^ cells at the end of their expansion cycle (resting state) were also sorted for live cells prior to scRNA-seq on day 0. Harvested T cells and remaining AML cells and were co-cultured for 24 hours at a 1:1 (E:T) ratio, then stained with anti-NGFR and anti-CD33 mAb to sort CD4^IL10^ cells and AML cells, respectively (CD4^IL10^ cells are engineered to overexpress truncated *NGFR* as a marker gene, besides *IL10*). After sorting, cells were adjusted to a 1:1 ratio to allow equal representation of AML and CD4^IL10^ cells in scRNA-seq data. This was repeated twice, so that each AML sample was co-cultured with T cells from two different donors. scRNA-seq was completed at Stanford Functional Genomics Facility (SFGF) using the Chromium Single Cell Gene Expression kit (10X Genomics, CA, USA) and Illumina Novaseq (Illumina Inc, CA, USA) sequencer according to manufacturer’s protocols.

### Single-cell analysis

Raw counts output by CellRanger^26^ were loaded into Seurat v4.0.5^27^ according to the accompanying vignette; details and modifications from default settings are listed in **Supplemental Methods**.

### Cell type annotation

Cells were assigned to cell types using marker genes *ANPEP* (CD13), *CD33, CD3D* and *CD3E*. The Wilcoxon test was used to assess differential gene expression ^28^ between clusters, and *p*-values were adjusted for multiple hypothesis testing using the Benjamini-Hochberg method. Clusters with a *q*-value < 0.05 for *ANPEP* or *CD33* were considered AML, and those with *q*-value < 0.05 for *CD3D* or *CD3E* were considered CD4^IL10^ cells. Clusters over-expressing both AML and CD4^IL10^ markers, or none of the two, were filtered out.

### Identification of AML transcriptional programs

We first clustered AML cells from each patient/timepoint pair, then averaged the expression of each gene across cells from each patient/timepoint pair within each cluster and merged the resulting average transcriptomes into a single matrix, *T*^∗^. We filtered out clusters (columns) with < 25 cells and genes (rows) with no expression, resulting in matrix *T*. We identified AML cell programs by applying non-negative matrix factorization (NMF)^29^ on matrix *T* using 25 NMF restarts as in Luca et al^30^, details of which are in **Supplemental Methods**.

### AML-induced CD4^IL10^ cell transcriptional changes

DGE analysis was performed between CD4^IL10^ cells before and after co-culture with each AML sample. The Wilcoxon test z-scores of CD4^IL10^ cells co-cultured with sensitive and resistant AML cells were separately combined into meta-z scores^31^, with the natural logarithm of the number of cells as weights and converted to two-sided *p*-values. Genes with a *q*-value < 0.05 (Benjamini-Hochberg procedure) in only one of the two conditions were analyzed further.

### Ligand-receptor analysis

Potential ligand-receptor interactions between AML and CD4^IL10^ were identified using the CellChat database^32^, focusing on overexpressed genes in AML programs and in CD4^IL10^ cells after co-culture with sensitive or resistant AML, and interactions for which all ligand genes were overexpressed in an AML program and all receptor genes in the CD4^IL10^ cells, or vice versa.

### U937 in vivo experiments

U937 WT or *ICAM1-KO* cells were transduced with the lentiviral vector expressing GFP-Luciferase (plasmid was a kind gift from Dr. Le Cong) and sorted via FACS based on GFP and ICAM-1 expression. All male NSG mice were purchased from Jackson Labs (Bar Harbor, ME) at 5 - 6 weeks of age. PBS, 1 - 1.4 million U937-WT-Luc^+^ or U937-*ICAM1-KO*-Luc^+^ cells were injected to mice (n = 10 mice per cohort) at 8 weeks of age intravenously. Imaging was performed using IVIS imaging system (PerkinElmer, Shelton, CT) after intraperitoneal luciferin (Promega, Madison, WI) injections according to the manufacturer’s protocol. Five days post U937 or PBS injection, half of the mice in each cohort (n = 5 per group) were injected with 2 million CD4^IL10^ cells intravenously. Imaging was done on days 8 and 12 post U937 injection, and mice were sacrificed on day 14.

### Statistical analysis

Performed in R 4.0.2 for scRNA-seq, and in GraphPad Prism 9.5.1 for other *in vitro* and *in vivo* experiments. Statistical tests used (α at 0.05) are indicated in legends of corresponding Figures.

### Data sharing statement

scRNA-seq data is available at Stanford repository accessible here: https://drive.google.com/drive/folders/1OA3eHE-v5RIteOhd3glTTU6ECuRaunhv.

## Supplemental Methods

### Single cell RNA data preprocessing

After loading the 10X Genomics CellRanger output into Seurat 4.0.5, genes expressed in less than 3 cells and ribosomal genes were filtered out, and 28,570 genes remained. Cells expressing less than 500 reads and/or 500 features, and more than 50% mitochondrial reads were also filtered out. The mitochondrial cut-off was higher than suggested in Seurat^27^ vignette describing *ex-vivo* isolated peripheral blood mononuclear cell (PBMC) analysis, because cultured AML cells and CD4^IL10^ cells are expected to be more metabolically active than resting PBMC. Notably, both AML and CD4^IL10^ cells at each time point were FACS-sorted for live cells, and 10X capture and processing for scRNA-seq started within 1h post-sort.

### Bulk RNA data preprocessing

NCI TARGET bulk-RNA-seq dataset and the dataset used in **Figure 3A** were processed using STAR^33^, as described in Cieniewicz et al^16^. For each dataset, the transcript-per-million (TPM) values were used. TPM profiles of cell-lines from Cancer Cell Line Encyclopedia (CCLE) were downloaded from the Broad Institute’s website (https://data.broadinstitute.org/ccle/CCLE_RNAseq_rsem_genes_tpm_20180929.txt). Gene IDs were mapped to gene symbols using EnsDb.Hsapiens.v86^34^ package. The expression matrix was filtered for protein-coding genes and hematopoietic cell-lines. The resulting expression values were re-converted to TPM.

### Seurat analysis

Raw counts were scaled to 10,000 reads per cell, and log-normalized. The top 2000 genes were selected using the *vst* method implemented in function *FindVariableFeatures*, run with the default parameters. Clusters of cells were identified by *FindNeighbors*, using the top 30 PCA components and otherwise default parameters, and *FindClusters,* with the resolution parameter set to 0.8 and otherwise default parameters. UMAP plots were generated using the top 30 PCA components with the *RunUMAP* function.

### Differential gene expression ^28^ analysis

Unless otherwise specified, DGE analysis was performed using the function *FindMarkers,* run with parameters: mean.fxn = rowMeans, min.cells.group = 0, min.cells.feature = 0, min.pct = 0, logfc.threshold = 0, only.pos = FALSE, max.cells.per.ident = 500, return.thresh = 1.

### Gene-set enrichment analysis

We leveraged full transcriptomes of T cells: CD4 naïve, T cells: CD4 memory resting, and T cells: CD4 memory activated from Newman et al^35^. For each cell population, we ordered each gene in the transcriptome by calculating the average log_2_ fold change of each population relative to the others. From the gene lists identified as described in section *AML-induced CD4^IL10^ cell transcriptional changes*, top 50 with the highest average log_2_ fold change across CD4^IL10^ cells cultured with sensitive and resistant samples were evaluated in mean log2 fold change-ordered transcriptomes using pre-ranked Gene Set Enrichment Analysis (GSEA) (*fgsea* R package)^36^, with 1,000 permutations.

### Selection of AML transcriptional programs

We selected the number of programs at which the Cophenetic coefficient fell below 0.99 and ignored programs with less than 200 marker genes in the downstream analysis. To assign single cells and external expression data to the programs, the recovery procedure described in Luca et al^30^ was used. The single cells and bulk samples for which the abundance of the filtered programs was highest were considered unassigned. We used a recovery procedure^30^ to assign external expression data to programs, considering the single cells and bulk samples with the highest abundance of the filtered program as unassigned.

### Optimization of published CRISPR/Cas9 knock-out strategy^25^

As previously described^25^, we used the kits (Synthego, CA, USA) that contained 3 individual single-guide ^37^ RNAs per target to knock out individual *IFNGR1, TNFRSF1B,* and *ITGB2*. Ribonucleoprotein (RNP) complexes were generated by mixing sgRNAs (30 pmol each) with 62 pmol of HiFi Cas9 (Integrated DNA Technologies, IA, USA). We optimized the published protocol by adding of 4 μM electroporation enhancer (Integrated DNA Technologies, IA, USA) and using the SF Cell Line nucleofection solution (Lonza, Switzerland) for the sensitive AML cell line U937.

## RESULTS

### Single-cell interaction analysis of primary AML and cytotoxic CD4^IL10^ cells

To identify CD4^IL10^ killing-sensitive and -resistant primary AML samples, we co-cultured primary AML cells with CD4^IL10^ cells (killing assay; see **Methods**) and determined that AML cells from patients PARCEV and PARCHW were sensitive, while AML cells from patients PATISD and PAPZCL were resistant to CD4^IL10^ mediated killing (**Figure 1A**, **Figure S1; Table S1**). As previously shown, CD4^IL10^ cells expressed the expected phenotype^16,17^, including CD2, CD18, CD226, and other surface proteins (**Figure S2**).

**Figure 1.**
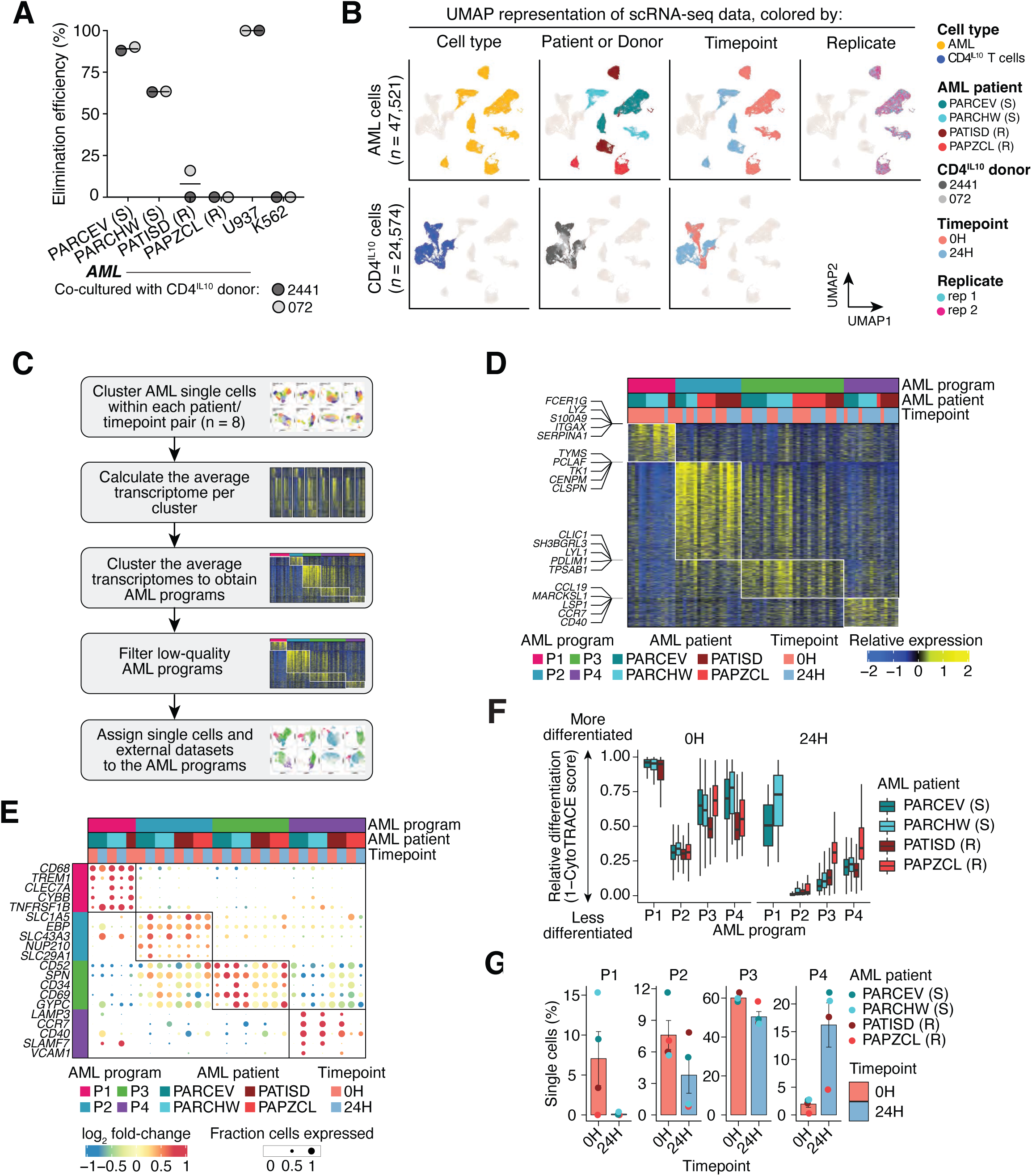
Longitudinal single-cell analysis of primary AML cell/CD4^IL10^ cell interactions identifies AML transcriptional programs. **A.** Each aliquot of primary AML cells was co-cultured with CD4^IL10^ cells for 3 days in 1:1 ratio, then enumerated using flow cytometry, and killing expressed as % elimination efficiency (**Methods**). X-axis indicates AML sample ID and their sensitivity (S) or resistance (R) to CD4^IL10^ cell killing. CD4^IL10^ killing-sensitive, monocytic myeloid leukemia cell line U937 was used as positive control, while CD4^IL10^ killing-resistant, erythroleukemic K562 cell line was used as negative control in each experiment. **B.** UMAP representation of scRNA-seq data (**Methods**), showing AML (**upper panel**) and CD4^IL10^ single cells (**lower panel**). **C.** Workflow depicting the derivation of AML programs (**Methods**). **D.** Heatmap of the four AML transcriptional programs identified. Rows represent genes, columns average expression of genes in sample-specific clusters, and colors indicate the relative expression of genes. **E.** Top 5 genes encoding surface molecules, ranked by log_2_-fold change, which are differentially expressed in each AML program (**Methods**). Bubble size = the proportion of cells expressing the gene; bubble color = log_2_-fold change. **F.** Relative differentiation of single cells assigned to each AML program, across timepoints, using CytoTRACE. **G.** Percentage of AML single cells assigned to each program at 0 and 24h-time points.

To understand how sensitive and resistant AML cells change transcriptionally after their interaction with CD4^IL10^ cells, and how the contact with either type of AML affects CD4^IL10^ cells, we performed scRNA-seq before (0h) and 24 hours after (24h) AML-CD4^IL10^ cell co-culture (**Figure S3A**). To obtain high-quality samples, we sorted live AML cells and CD4^IL10^ cells (**Methods**) immediately prior to single-cell capture for scRNA-seq. Unlike in the 3-day killing assay (**Figure S1**), 24-hour co-culture did not result in complete elimination of sensitive AML cells by CD4^IL10^ cells (**Figure S3B**).

Following pre-processing and quality control, 72,095 single cells were considered for downstream analysis. Of those, 47,521 cells were annotated to AML and 24,574 to CD4^IL10^ cells (**Methods; Table S3).** AML cells showed heterogeneity between patients, but not between replicates (i.e., two aliquots of the same AML patient sample thawed and analyzed at different time points; **Figure 1B**), suggesting that the variability introduced by sample processing was minimal. CD4^IL10^ cells derived from healthy donors showed less heterogeneity than AML (**Figure 1B**). Within the co-cultures, the transcriptome of both AML and CD4^IL10^ cells separated by timepoint, suggesting that *in vitro* incubation by itself induced transcriptional changes.

### AML samples contain cells expressing four different transcriptional programs

To resolve the heterogeneity of AML samples while mitigating patient- and *in vitro*-induced differences, we employed a two-tiered approach (**Methods**; **Figure 1B**). We first clustered AML single cells within each patient at each timepoint (**Methods; Figure S4A**), then calculated the average transcriptome per cluster. The averages were then clustered, revealing five AML cell transcriptional programs shared across all patients and timepoints (**Figure S4B,C**). One of the programs was low quality (< 200 overexpressed markers features and > 20% of mitochondrial reads; **Figure S4D**) and was removed from further analysis.

The remaining four programs revealed major AML transcriptional heterogeneity (**Figure 1C,D, S4E; Table S3**). Program P1 was enriched mostly in cells from killing-sensitive AML samples at 0h. Programs P2 and P3 were enriched in all samples and conditions, while program P4 was found in cells mostly after 24 hours of co-culture (**Figure 1D**). Top 5 genes encoding surface proteins found in each program are depicted in **Figure 1E** (complete list in **Table S3**). Cells enriched in program P1 overexpressed genes associated with mature monocytes or other myeloid cells, such as *S100A8*/*S100A9, CD68*, *CD14*, integrin- and adhesion protein-encoding genes *ITGAX* and *ITGB2*, *ICAM1* (encoding CD54), and cytokine or cytokine receptor-encoding genes such as *IL10RA*, *TNFRSF1B*, and *IFNGR1*. We did not find notable immune-related genes in program P2. Cells enriched in program P3 overexpressed *CD34*, which is expressed by hematopoietic stem cells (HSC) and leukemic stem cells (LSC)^38^. AML cells enriched in program P4 overexpressed genes encoding HLA class II proteins, as well as *CD200,* found on AML and LSC^39^, for which we have previously elucidated the role in AML resistance to CD4^IL10^ killing^16^ (**Figure 1E**).

Because the AML programs P1-P4 contained genes associated with different maturation levels of hematopoietic cells, we further assessed the comparative differentiation stage of AML cells in programs P1-P4 using CytoTRACE^40^. AML cells in program P1 had highest estimated differentiation, whereas AML cells in programs P2-P4 had a lower estimated differentiation (**Figure 1F**). Cells that survived 24h culture, irrespective of the program, displayed a less differentiated transcriptional signature (*p* < 10^-16^, Wilcoxon test; **Figure 1F**). These differentiation levels were in accordance with the French-American-British (FAB) AML classification^41^, which is based on morphology and differentiation stage. FAB classification was reported for two out of four AML patient samples used in the analysis: sensitive PARCEV sample was FAB M4 (myelomonocytic AML), while resistant PAPZCL sample was M0 (undifferentiated AML; **Table S1**).

Next, we examined the representation of AML programs across time. Program P1 was present in sensitive AML samples initially, but disappeared after 24-hour co-culture (**Figure 1G**), suggesting an association of program P1 with sensitivity to killing. Representation of AML cells expressing program P2 was overall low, and decreased in some, but not all samples after co-culture. On the other hand, majority of AML cells expressed program P3, and this remained largely unchanged after co-culture. AML cells expressing program P4 increased after co-culture (**Figure 1G**). These data suggest an association of programs P3 and P4 with resistance to CD4^IL10^ cell-mediated killing.

### AML transcriptional programs correlate with AML susceptibility to T cell killing and patient survival

Our scRNA-seq analysis suggested that AML transcriptional programs are linked to AML differentiation stage and sensitivity to CD4^IL10^ killing. Next, we asked whether they associate with: 1) sensitivity to killing in a larger, independent cohort of AML samples; 2) AML FAB subtypes^41^, risk group, or World Health Organization (WHO) classification^28^ and 3) AML patient survival. First, we correlated the abundance of each AML transcriptional program in bulk RNA-seq samples from our previously published AML cohort (n = 14)^16^, which had known sensitivity to CD4^IL10^ killing (measured by elimination efficiency; see **Methods**). In line with our observations in single-cell data, the abundance of the AML program P1 positively correlated with killing (*r* = 0.48, *p* = 0.086), while the abundance of programs P3 and P4 showed a significant negative correlation with killing (r = −0.6, *p* = 0.022, and r = −0.6, *p* = 0.023, respectively; **Figure 2A**).

**Figure 2.**
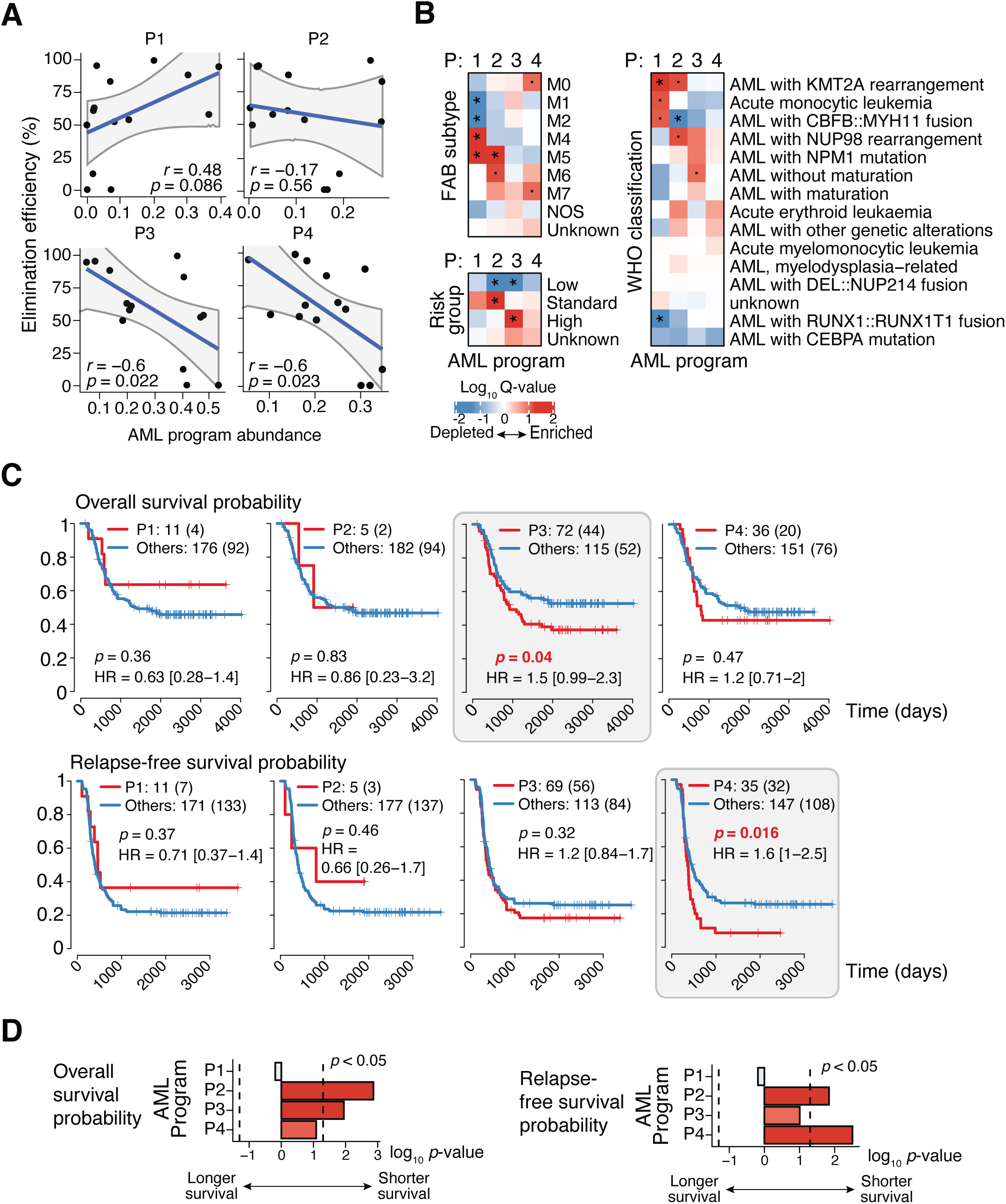
AML transcriptional programs that correlate with resistance to T cell killing also correlate with poor patient survival. **A.** The association of AML program abundance with sensitivity to CD4^IL10^ killing across AML samples with known elimination efficiencies (n = 14) analyzed by RNA-seq^7^ (**Supplemental Methods**). **B**. Heatmap of the log_10_ *q*-values of the enrichment of AML programs in AML samples from the TARGET cohort (**Methods, Supplemental Methods**) with known French-American British (FAB) subtypes (**top left**), risk groups (**bottom left**), and WHO subtypes (**right**). Enrichment was calculated by performing Wilcoxon tests between the program abundance of samples from each subtype and the rest of the categories. * – *q* < 0.05, · – *q* < 0.1. **C.** Kaplan-Meier plots of the association of samples from the TARGET cohort grouped by the dominant AML program with overall survival (**top row**) and relapse-free survival (**bottom row**). *p* – log-rank *p*-value, HR – hazard ratio. Numbers in square brackets indicate the 95% confidence interval of the hazard ratio, and the numbers in round brackets indicate the number of events. **D**. The –log_10_ *p*-values of the association of AML program abundance with overall survival (**left**) and relapse free survival (**right**), calculated using univariate cox models.

As tumor biology studies often utilize model cell lines, we scored the abundance of AML programs P1-P4 in cell lines derived from patients with leukemias and lymphomas, using the Cancer Cell Line Encyclopedia (CCLE) RNA-seq dataset^42^ (**Figure S5A-C, Table S4**). The cell lines with high abundance of AML program P1 were derived from mature monocytic tumors, including THP-1 and U937, which we use as killing-sensitive controls in CD4^IL10^ killing assays^14–16^ (**Figure S5B, S5C**). Other cell lines, such as the killing-resistant K562 cells with undifferentiated/erythroleukemic phenotype that we use as a negative control for CD4^IL10^ killing, exhibited low P1 but high P3 and P4 scores (**Figure S5B, S5C**). Thus, the relative AML program abundance in RNA-seq data of tumor cell lines can be an indicator of their sensitivity to CD4^IL10^ cell killing.

We also evaluated the association of AML transcriptional programs with the FAB classification^41^, the WHO classification, and risk groups in a large cohort of pediatric AML samples from the National Cancer Institute (NCI) TARGET RNA-seq dataset (**Figure 2B and S5D)**. The AML program P1, associated with sensitivity to killing, was significantly depleted from FAB subtypes M1 and M2 (acute myeloblastic leukemia with minimal maturation or with maturation, respectively), and was significantly enriched (*q* < 0.05) in AML subtypes M4 (acute myelomonocytic leukemia) and M5 (acute monocytic leukemia) (**Figure 2B**). P1 was also significantly enriched in WHO subtype AML with *KMT2A* rearrangement (*q* < 0.05), acute monocytic leukemia, and AML with *CBFB::MYH11* fusion (**Figure 2B**). We did not observe an association between P1 program abundance and risk groups, but P1 was depleted from samples with *RUNX1::RUNX1T1* fusion (**Figure 2B**), which is associated with worse prognosis in pediatric AML^43^.

The AML program P2, with undefined association to killing (**Figure 2A**), was significantly enriched in FAB subtype M5 (*q* < 0.05), enriched in FAB subtype M6 (acute erythroid leukemia) and WHO subtype AML with *KMT2A*, as well as in AML with *NUP98* rearrangement, which has poor prognosis^44^. Program P2 was significantly depleted from WHO subtype AML with *CBFB::MYH11* fusion (**Figure 2B**), which is mainly associated with good survival^45^, significantly depleted from low-risk AML, and significantly enriched in AML with standard risk (*q* < 0.05). Program P3, associated with resistance to killing, was found enriched in AML without maturation and high-risk AML, while being significantly depleted from low-risk AML. Program P4, also associated with resistance to killing, was enriched in AML subtype M0 (undifferentiated acute myeloblastic leukemia) and M7 (acute megakaryoblastic leukemia; **Figure 2B**). Altogether, AML cells expressing the T cell killing sensitivity-associated program P1 had mature myeloid-like features and mutations linked to better survival, while AML cells expressing program P2 and killing resistance-associated programs P3 and P4 had less differentiated features, and were linked to higher risk AML.

Finally, we examined the association of AML transcriptional programs with patient survival, using the TARGET RNA-seq dataset analyzed in our previous study^16^ (n=187). Patient samples with enrichment of program P1, associated with sensitivity to T cell killing, showed a trend towards better survival outcomes, but it was not statistically significant (**Figure 2C**). In contrast, AML patients with enrichment of program P3 and P4, associated with resistance to killing, had significantly reduced overall survival (P3; log-rank *p =* 0.04, HR = 1.5, 95% CI = [0.99 – 2.3]; **Figure 2C** upper panel) and relapse-free survival (P4; log-rank *p =* 0.016, HR = 1.6, 95% CI = [1 – 2.5]; **Figure 2C** lower panel). These enrichments were also consistent when considering the abundance of programs as continuous variables in univariate Cox models, where program P2 was also found in patients with worse outcomes (**Figure 2D**).

Altogether, these AML programs, identified by studying the AML/T cell interaction, reinforce the importance of T cell response in AML elimination, and suggest that the expression of T cell killing-resistance associated programs P3 and P4 in patient AML cells could indicate a risk of poor outcome.

### Only killing-sensitive AML cells activate CD4^IL10^ cells

After mitigating the effect of *in vitro*-induced transcriptional changes (**Methods**), we also analyzed the transcriptional changes occurring in CD4^IL10^ cells after interaction with sensitive or resistant AML cells (**Figure 3A**). The transcriptome of CD4^IL10^ cells co-cultured with sensitive AML was markedly more perturbed than those co-cultured with resistant AML (1,646 vs. 343 differentially expressed genes). This suggests that CD4^IL10^ cells make more interactions (or higher affinity interactions) with sensitive AML compared to the resistant ones. CD4^IL10^ cells co-cultured with sensitive AML upregulated many genes associated with T cell activation, such as genes encoding for activation-induced proteins *IL2RA* (CD25) and *TNFRSF4* (OX40), cytokines *CSF1* (M-CSF), *CSF2* (GM-CSF), *IFNG*, and *LTA*, metabolism related genes such as *FABP5* and *PKM*, and co-inhibitory molecules such as *HAVCR2* (TIM3) and *CTLA4* (**Figure 3A**; full list in **Table S5**). CD4^IL10^ cells co-cultured with resistant AML upregulated few T cell activation-related genes. Accordingly, gene set enrichment analysis revealed that CD4^IL10^ cells co-cultured with sensitive AML cells have a transcriptional signature of activated memory CD4^+^ T cells, while CD4^IL10^ cells co-cultured with resistant AML cells have a signature of resting memory CD4^+^ T cells (**Figure 3B, S6A**). Collectively, these data suggest that sensitive AML cells can engage and activate T cells, which facilitates their killing, whereas the resistant AML cells do not activate T cells and survive the interaction.

**Figure 3.**
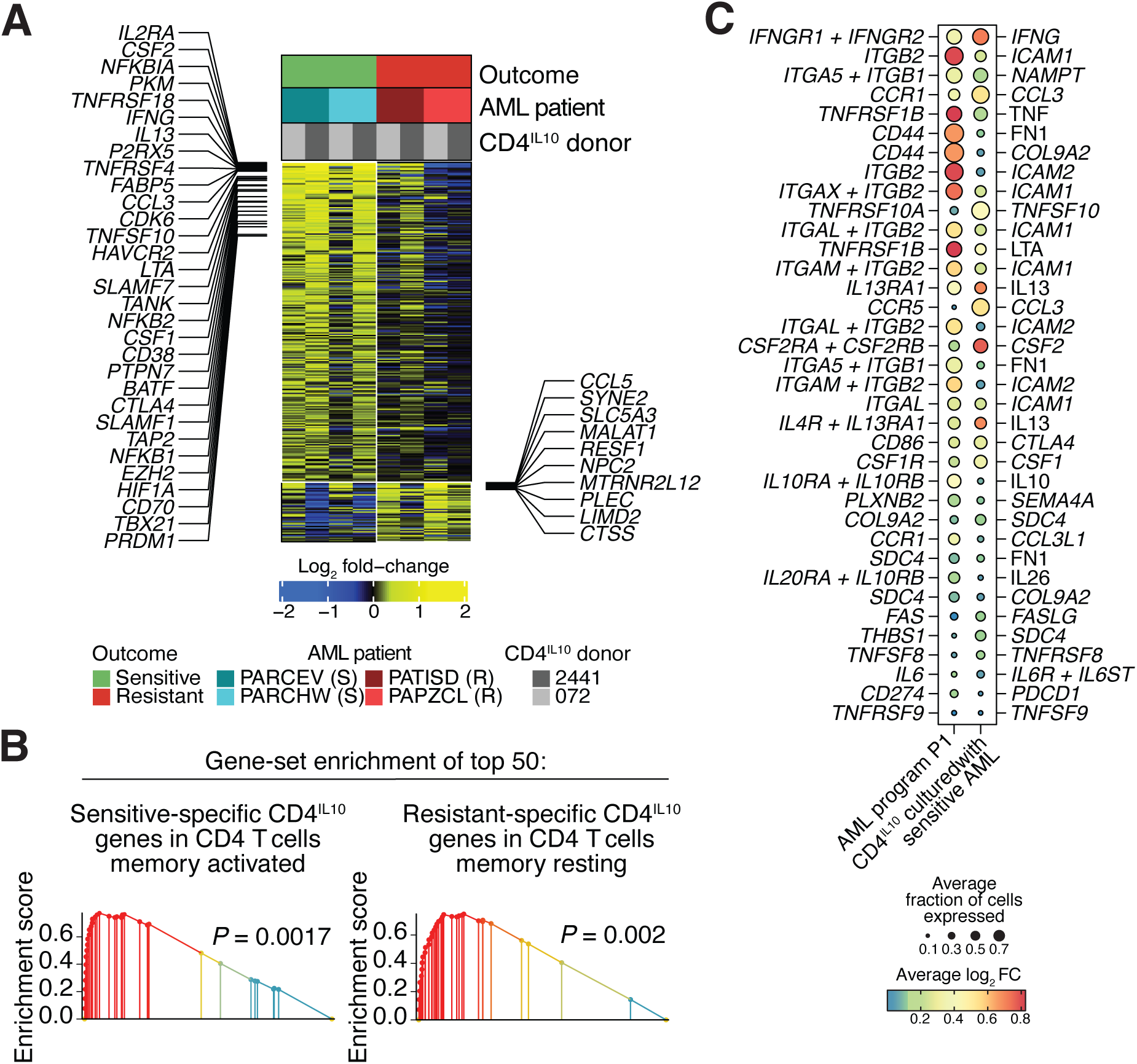
Killing-sensitive AML cells activate CD4^IL10^ cells. **A.** Heatmap of the genes significantly changing in CD4^IL10^ cells after co-culture with sensitive or resistant blasts (**Methods**). Rows – genes, columns – CD4^IL10^ samples, colors – gene expression. **B.** Gene-set enrichment of the top 50 genes uniquely over-expressed in CD4^IL10^ after co-culture with sensitive blasts (**left**) and with resistant blasts (**right**) (**Supplemental Methods)**. **C.** Bubble plots representing the potential cell-cell interactions between AML cells expressing sensitivity-related program P1 and CD4^IL10^ cells co-cultured with sensitive AML (**Methods**). Bubble sizes are proportional to the average fraction of single cells in each population expressing the genes encoding the first and second interaction partners. Colors indicate the average log_2_ fold change of genes encoding the interaction partners in the AML program P1 relative to other AML cells and in the CD4^IL10^ cells after culture with sensitive AML. The plus sign separates subunits of multi-subunit receptor complexes.

To identify potential receptor-ligand interactions between AML cells and CD4^IL10^ cells, we leveraged known receptor-ligand pairs from the CellChat database^32^ (**Methods**), focusing on the genes upregulated in both AML and CD4^IL10^ cells after their interaction. For AML cells expressing the sensitivity-associated transcriptional program P1, some of the top interacting pairs were the genes encoding for: a) receptors for IFN-γ, TNF-α and lymphotoxin A ^41^ cytokines (encoded by *IFNGR1* and *TNFRSF1B*) on AML, and cytokines *IFNG, LTA* and *TNF* on CD4^IL10^ cells; and b) integrins *ITGB2* (encoding CD18*), ITGAL* (CD11a), *ITGAM* (CD11b) and *ITGAX* (CD11c) on AML cells, and adhesion proteins *ICAM1* (encoding ICAM1, also known as CD54) and *ICAM2* (CD102) on CD4^IL10^ cells (**Figure 3C**).

AML cells enriched in resistance-associated programs P3 and P4 displayed a different, less abundant interaction profile with CD4^IL10^ cells. For AML program P3, top interacting pair was between *LGALS9* (encoding galectin-9) on AML cells and *CD44* on CD4^IL10^ cells (**Figure S6B)**. For AML program P4, top interacting pair was between *ICAM1*, here expressed on AML cells, and *ITGAL* (CD11a) on CD4^IL10^ cells (**Figure S6C**).

Overall, sensitive AML were successful in activating the T cells, likely by making more cell-cell interactions, and are subsequently killed by T cells; whereas resistant AML fail to activate the T cells and evade T cell killing.

### ICAM1/LFA-1 interaction is required for AML killing by CD4^IL10^ and primary CD8^+^ T cells

To validate predicted AML/T cell interactions, we knocked out several highly expressed genes found in program P1 from the otherwise killing-sensitive AML cell line U937, and quantified the effect of the knock-out on CD4^IL10^ cell-mediated killing of U937 cells (**Methods**). Knockout of TNF-α receptor (encoded by *TNFRSF1B*) and IFN-γ receptor (*IFNGR1)* from U937 cells did not alter their sensitivity to killing. Similarly, knockout of CD18 integrin (*ITGB2*) from U937 cells did not have an effect (**Figure 4A, S7A**).

**Figure 4.**
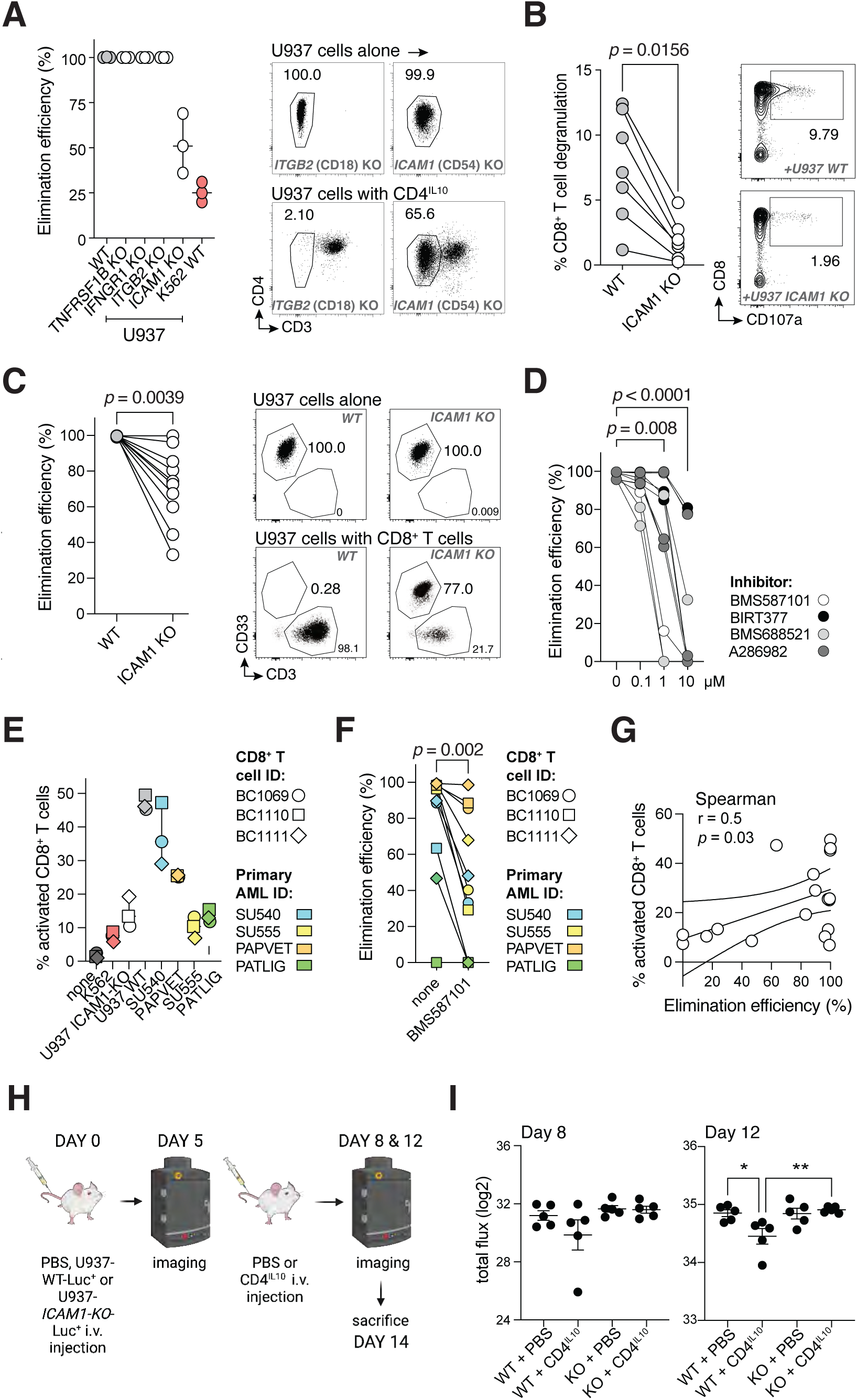
ICAM1/LFA-1 interaction is essential for AML killing by CD4^IL10^ and *ex vivo* isolated CD8^+^ T cells. **A.** Elimination efficiency of U937 cell line after CRISPR-Cas9 mediated knockout (KO) of selected genes by CD4^IL10^ cells; each dot represents a CD4^IL10^ donor (n=3). WT = wild type; U937 WT: positive control, K562 WT: negative control. Representative FACS plots from one donor against U937-*ITGB2*-KO and U937-*ICAM1*-KO are shown on the right; numbers indicate percentage in AML gate. **B.** Degranulation of CD8^+^ T cells, expressed as % CD8^+^CD107a^+^ cells within live singlet CD3^+^ T cells, against U937 WT and U937-*ICAM1*-KO cells; each circle represents a CD8^+^ T cell donor (n = 7). Representative FACS plots from one donor are shown on the right; numbers indicate percent degranulating CD8^+^ T cells. **C.** Elimination efficiency of U937 WT and U937-*ICAM1*-KO cells by CD8^+^ T cells (n = 10). Wilcoxon test, *p* < 0.05. Representative FACS plots from one donor are shown on the right; numbers indicate percentage in AML gate. **D.** Elimination efficiency of U937 cell line 24 hours after treatment with LFA-1 inhibitors at indicated concentrations; each line represents a CD4^IL10^ donor (n = 3); circles are color-coded by the inhibitor as indicated. Dunn’s post hoc test results are shown, following Friedman ANOVA (*p* < 0.0001). **E.** CD8^+^ T cells were co-cultured for 24hr with indicated target cells; activation was measured as percent of CD69^+^CD25^+/-^ cells within the live CD8^+^ T cell gate. **F.** Elimination efficiency of indicated target cell lines and primary AML cells by CD8^+^ T cells after 3 days of co-culture in the absence or presence of LFA-1 inhibitor BMS587101 (10 μM). **G.** Spearman correlation of percent activation of CD8^+^ T cells and the elimination efficiency of target cells. **H.** Timeline of mice injections; n = 10 mice per group. **I.** Graphs showing total flux obtained from imaging data on days 8 and 12. 1-way ANOVA with Bonferroni multiple comparison test; *p* = n.s. at day 8, *p* = 0.0075 at day 12.

Notably, CD18 and CD11a are subunits of the LFA-1 (lymphocyte function-associated antigen 1) protein. LFA-1, when expressed on T cells, binds to ICAM1 on target cells, enabling immune synapse formation and T cell-mediated killing of target cells^46^. CD4^IL10^ cells constitutively express high levels of LFA-1 subunits CD18^15^ (*ITGB2*; **Figure S2**) and CD11a^17^ (*ITGAL*, **Figure S6D, S6E**). *ITGB2* and *ITGAL* were not further upregulated in CD4^IL10^ cells interacting with killing-sensitive AML cells, and thus not identified by CellChat (**Figure 3C**). However, *ICAM1* is one of the top upregulated genes in AML cells expressing sensitivity-associated program P1 (**Table S3**) and found on sensitive AML samples in other programs as well (**Figure S6F**). Therefore, we also knocked-out *ICAM1* on U937 cells (U937-*ICAM1*-KO), which significantly decreased their sensitivity to CD4^IL10^ killing (**Figure 4A, Figure S7A**). Thus, ICAM1 expression on AML is one of the key mechanisms that enables AML killing by CD4^IL10^ cells.

ICAM1/LFA-1 interaction is critical for the immune synapse formation between endogenous cytotoxic cells, such as CD8^+^ T cells, and tumor cells *in vivo*^46,47^. This could help explain why patients with abundant AML programs P3 and P4, associated with low *ICAM1* expression and resistance to CD4^IL10^ killing, have poor survival. To test whether ICAM1/LFA-1 interaction governs AML killing by *ex vivo*-isolated, primary CD8^+^ T cells, we co-cultured human CD8^+^ T cells with killing-sensitive, wild-type U937 cells or U937 lacking ICAM1 (U937-*ICAM1*-KO). To bypass the diverse endogenous T cell receptors (TCR) on primary human CD8^+^ T cells, we pre-incubated U937 cells with low-dose soluble anti-CD3 antibody. Myeloid cells such as U937 cells express Fc receptors that bind the anti-CD3 antibody, providing a non-specific TCR stimulus together with their own co-stimulatory signals to T cells. When both TCR activation and co-stimulation signals are present, CD8^+^ T cells are expected to kill U937 cells. Indeed, primary human CD8^+^ T cells released cytotoxic granules and killed almost all wild-type U937 cells, whereas their degranulation and killing of U937-*ICAM1*-KO cells was significantly impaired (**Figure 4B-C**). Similar to CD4^IL10^ cells, CD8^+^ T cells did not degranulate and kill resistant, lymphoblastoid K562 cells (**Methods**). Thus, ICAM1 expression on AML is necessary for AML killing by primary CD8^+^ T cells.

To understand whether LFA-1, the ICAM1 binding partner expressed on T cells, is equally important as ICAM1 for AML sensitivity to killing, we added small molecule inhibitors of LFA-1 or ICAM1/LFA-1 interaction (BMS587101, BIRT377, BMS688521, A286982) to the AML/CD4^IL10^ cell co-culture (killing assay; **Methods**). All four agents significantly increased the resistance of wild-type U937 cells to CD4^IL10^ killing in a dose-dependent manner (**Figure 4D, Figure S7C**), further confirming the role of ICAM1/LFA-1 interaction for effective AML killing by T cells.

Next, we showed that only the killing-sensitive, wild-type U937, but not resistant U937-*ICAM1*-KO cells, potently induce primary CD8^+^ T cell activation, as measured by frequency of CD69^+^CD25^+/-^ CD8^+^ T cells after 24h co-culture (**Figure 4E; Figure S7D**). Thus, primary CD8^+^ T cell activation in response to sensitive U937 cells was aligned with CD4^IL10^ cell activation in response to sensitive primary AML cells (**Figure 3A-B**). We also co-cultured these CD8^+^ T cells with primary adult (SU540, SU555) and pediatric (PAPVET, PATLIG) AML samples, for which we previously determined to be sensitive to CD4^IL10^ cell killing. Two out of four primary AML samples induced strong CD8^+^ T cell activation after 24 hours of interaction (**Figure 4E; Figure S7D**); after 72h of interaction, three out of four primary AML samples were also killed by CD8^+^ T cells (**Figure 4F**). Sample PATLIG had low expression of CD64 and ICAM1, both of which could explain its resistance to killing, as low CD64 expression is likely not sufficient to cross-link the anti-CD3 antibody to provide a TCR stimulus (**Figure S8A-B**). Further, killing of primary AML by CD8^+^ T cells was significantly impaired by an LFA-1 inhibitor when ICAM1 was expressed on AML (**Figure 4F; Figure S7D, Figure S8A-B**), showing that LFA-1 signaling is critical for AML killing. We also observed a positive correlation between the degree of CD8^+^ T cell activation at 24h and AML cell killing (**Figure 4G**).

Finally, we tested the importance of *ICAM1* expression *in vivo*, in a Luciferase^+^ (Luc^+^) U937 cell-induced AML model. We injected U937-WT-Luc^+^ or U937-*ICAM1-KO*-Luc^+^ cells into NSG mice, and 5 days later injected CD4^IL10^ cells into half the mice per U937 group. Tumor progression was followed by imaging at day 5 (before CD4^IL10^ injection), day 8 and 12 post U937-injection (**Figure 4H-I and S9**). On days 8 and 12, we observed a reduction in tumor burden in CD4^IL10^-treated U937-WT mice, but not in *ICAM1-KO* mice; the reduction on day 12 was statistically significant (**Figure 4I**). Thus, ICAM1 expression on killing-sensitive AML cells contributes to AML clearance by CD4^IL10^ cells *in vivo*.

Collectively, our data suggest that ICAM1/LFA-1 interaction is one of the key mechanisms that governs AML elimination by cytotoxic CD4^+^ and CD8^+^ T cells, which could influence AML control in patients.

## DISCUSSION

In this study, we leveraged longitudinal single-cell transcriptomics to understand the determinants of AML sensitivity or resistance to T cell killing. We found that killing resistance-associated AML transcriptional programs are enriched in samples of AML patients with poor survival rates. Killing-sensitive AML cells expressed different transcriptional programs than resistant AML, and were able to activate the CD4^IL10^ cells through receptor-ligand interactions. A key interaction that defined AML sensitivity to killing by CD4^IL10^ cells and primary human CD8^+^ T cells was mediated by ICAM1, which binds to LFA-1 on T cells and facilitates immune synapse formation. These data add a mechanistic basis to existing studies showing that functional T cell responses are important for AML patient survival^4,5^, and suggest that ICAM1 expression on AML acts as an immune trigger, enabling T cell activation and AML killing.

CD4^IL10^ cells recognize and kill most primary AML cells *in vitro* independent of a single-antigen recognition via their TCR^14–16^, and inhibit leukemia *in vivo*^15^. These properties make them useful to study T cell-AML interaction, bypassing the requirement to isolate rare AML antigen-specific T cells from AML patients. CD200-based mechanism of resistance to T cell killing was observed both on CD4^IL10^ cells and primary human CD8^+^ T cells, and we show herein that *ICAM1* expression on AML has a comparable effect on CD4^IL10^ cell and primary, *ex vivo*-isolated human CD8^+^ T cell function. Thus, CD4^IL10^ cells can be used in relevant human models to study AML-T cell interaction.

Examination of primary AML/CD4^IL10^ interactions by longitudinal scRNA-seq identified four AML transcriptional programs: P1-P4. Program P1 was associated with AML sensitivity to killing, and was enriched for genes associated with mature monocytic cells, integrins, and adhesion proteins, such as *CD68*, *CD14*, *ITGAX* (CD11c), *ITGB2* (CD18), and *ICAM1* (encoding ICAM1 protein, also known as CD54). Additionally, P1 contained *S100A8* and *S100A9*, which are considered to contribute to myeloid differentiation of leukemic cells^50^. Expression of program P2 did not clearly correlate with sensitivity or resistance to killing. AML cells enriched in program P3 over-expressed *CD34*, which is ordinarily expressed on LSC^38^ (as well as normal hematopoietic stem cells; however primary AML bone marrow samples we tested contained ≥ 80% AML cells). Program P4 contained HLA class II genes, as well another gene described in AML LSC: *CD200*^39^, which impairs cytotoxic function of NK cells^24^, CD8^+^ T cells^23^, and CD4^IL10^ cells alike^16^. AML LSC are known for contributing to relapse and evading the immune response^39^, which could be mediated in part through their resistance to T cell killing.

We also investigated if the AML transcriptional programs correlate with patient outcomes. Strikingly, enrichment in AML programs P3 and P4 was significantly associated with poor patient survival. Providing this association is validated in independent cohorts, genes in programs P3 and P4 could be derived into useful biomarkers to identify high-risk patients suitable for a more aggressive therapeutic approach. Genetic aberrations with well-defined risk are not present in all patients^51^, leaving many AML patients without prognostic biomarkers^52,53^. For example, AML with *KMT2A* rearrangement is one of the most common AML subtypes in children, but *KMT2A* fusion partners – more than 80 of which are known thus far^28^ - determine survival outcomes^54^. Abundance of program P1, associated with killing-sensitive AML and enriched in patients with *KMT2A* rearrangement, could facilitate risk stratification of AML patients with *KMT2A* rearrangement. On the other hand, program P2 was more abundant in patients with poor survival, and also depleted from AML with *CBFB::MYH11* fusion, which is in most cases associated with good outcomes. The applicability of AML transcriptional programs in assessing AML risk will need to be validated in other suitable AML cohorts, ideally pediatric, because adult AML differs from pediatric AML on genetic, epigenetic, and transcriptional level^55–57^. However, aside from TARGET, large public RNA-seq datasets of pediatric AML are few, and not necessarily appropriate for survival analysis, e.g., when outcomes are mostly favorable^58^.

Several studies suggested the importance of functional T cell immune response for AML control^6^. In AML patients that relapse after HSCT, memory T cells appear exhausted^59^. In addition, AML cells in patients refractory to HSCT downregulate the HLA class II genes^10,11^, and deregulate co-stimulatory molecules^11^, both of which are critical to activate CD4^+^ Th cells that provide help to cytotoxic CD8^+^ T cells. Moreover, CD8^+^ T cells are affected directly by interaction with AML cells, acquiring a senescent state^8^, which abolishes their capacity to kill autologous AML cells^9^. Interestingly, *CBFB-MYH11* fusion, which is associated with good patient outcomes, can create a neoantigen recognized by CD8^+^ T cells, leading to AML control in a murine model^60^. These data support the notion that solidifying T cell/AML interactions and/or re-invigorating T cells *in vivo* could be a viable therapeutic approach for AML. Indeed, CD4^IL10^ cells can be developed as a therapy for killing-sensitive subtypes of AML.

Interaction with AML also induces transcriptional reprograming of T cells. CD4^IL10^ cells interact more with sensitive AML cells enriched in program P1, and respond by upregulating genes involved with T cell activation and memory. These genes include many integrins and adhesion proteins, which can strengthen the immunological synapse – a pre-requisite for productive T cell killing of target cells^61^. In addition, top upregulated genes in CD4^IL10^ cells co-cultured with sensitive AML encoded for proteins ordinarily secreted by cytotoxic CD4^+^ T cells in an immunological synapse, including IFN-γ, TNF-α, and LTA^62^. IFN-γ is particularly important, because it counter-acts the loss of HLA class II expression observed in AML relapse^10,11^. These data also illustrate that, despite constitutive expression of *IL10*, which is an immune regulatory cytokine, CD4^IL10^ cells predominantly activate their cytotoxic rather than immune regulatory functions when interacting with AML cells.

Killing-sensitive AML overexpress *ICAM1*, and ICAM1 binding partner LFA-1 is constitutively expressed on CD4^IL10^ cells^17^ and other cytotoxic lymphocytes. While efficient T cell killing requires multiple activation signals^63^, *ICAM1* upregulation on sensitive AML cells was notable because of the crucial role that ICAM1/LFA-1 interaction plays in the immune synapse between the target cells and cytotoxic lymphocytes^46,64,65^. Indeed, *ICAM1* silencing results in AML in escape from NK cell-mediated lysis^66^. In addition, we previously described a correlation between AML sensitivity to killing and ICAM1 expression in a small cohort of adult AML patients^15^, but we did not validate it experimentally. Here, we show that the knock-out of *ICAM1* from killing-sensitive AML cell line, U937 cells, significantly increased their resistance to killing by CD4^IL10^ cells *in vitro* and *in vivo*.

We also show that *ICAM1* expression enables AML killing by primary CD8^+^ T cells, which are key for AML control in patients. *Ex vivo*-isolated human CD8^+^ T cells degranulate significantly less in co-culture with U937-*ICAM1*-KO cells compared to wild-type cells, and U937-*ICAM1*-KO cells acquire resistance to killing by primary CD8^+^ T cells. As with CD4^IL10^ cells that are activated by killing-sensitive primary AML cells, which express high levels of *ICAM1*, *ex vivo*-isolated CD8^+^ T cells are activated by sensitive, wild-type U937 cells, but not by resistant U937-*ICAM1*-KO cells. Finally, inhibition of LFA-1, which is the T cell binding partner of ICAM1 on AML cells, prevented the killing of wild-type U937 cells and primary AML cells by *ex vivo*-isolated CD8^+^ T cells, further confirming that the requirement for ICAM1/LFA-1 interaction in AML cell killing is not restricted to CD4^IL10^ cells.

The strong impact of only one protein, ICAM1, on AML sensitivity or resistance to T cell killing was surprising. Previously, we uncovered that CD200 expression on AML confers resistance to killing, but this effect was mild, albeit significant. CD200, which is upregulated on T cell killing-resistant AML and AML LSCs, binds to the inhibitory CD200 receptor (*CD200R1*) on T cells, impairing their ability to degranulate and kill of AML cells. CD200 thus acts as an immune checkpoint, similar to CTLA-4, PD-1 and other immune checkpoints proteins^67^, preventing T cell activation. ICAM1, on the other hand, allows for T cell activation and killing to take place. As such, ICAM1 acts as an immune trigger, with an opposing function to immune checkpoint proteins. It is tempting to speculate that therapeutic enhancement of ICAM1/LFA-1 interaction between AML and T cells can enable T cell activation and tumor control. ICAM1 may play the same role in other malignancies, and it is likely that other proteins can exhibit an ICAM1-like function of an immune trigger.

In summary, we utilized scRNA-seq to reveal that: (1) expression of AML transcriptional programs (P1-P4) affect AML cells’ sensitivity or resistance to CD4^IL10^ cell-mediated killing, (2) program P1 is enriched in T cell killing-sensitive and mature myeloid-like AML, (3) programs P3 and P4, enriched in T cell killing-resistant and stem cell-like AML, are associated with poor patient survival, (4) CD4^IL10^ cells and primary CD8^+^ T cells that interact with killing-sensitive, but not resistant AML cells, acquire an activated T cell profile, and (5) disruption of only one interaction, between ICAM1 on AML cells and LFA-1 on T cells, was sufficient to significantly increase AML resistance to killing both *in vitro* and *in vivo* by CD4^IL10^ cells and *in vitro* by primary CD8^+^ T cells. TCR engagement without the immune synapse formation is not always sufficient to ensure AML killing. Overall, this study highlights the importance of AML-T cell interactions in AML immune escape, suggesting that the ability of AML tumor cells to productively engage T cells is important for AML patient survival.

## ACKNOWLEDGEMENTS

The authors would like to thank Drs. Steven Strubbe and Šimon Borna for valuable discussions; Dr. Rhonda Perriman for her help in obtaining funding and critical reading of the manuscript, Dr. Le Cong and Dr. Ravi Kumar Dinesh for the gift of GFP-Luciferase plasmid, and Michael Blanco from SFGF for expert assistance in library preparation and sequencing. Schematic figures were created with BioRender. The results published here are in part based upon data generated by the TARGET (Therapeutically Applicable Research to Generate Effective Treatments) (https://ocg.cancer.gov/programs/target) initiative, phs000465. (The data used for this analysis are available at https://portal.gdc.cancer.gov/projects.) *FUNDING:* This study was funded by NIH 1R21CA245468-01A1 (M.G.R.) and supplement 3R21CA245468-01A1S1 (B.C.T.), CURE Childhood Cancer (M.G.R.), Children’s Leukemia Research Association (M.G.R.), and Ludwig Cancer Research (M.G.R.) funding. The sequencing data was generated by SFGF with instrumentation purchased with NIH funds (S10OD025212 and 1S10OD021763).

## AUTHORSHIP CONTRIBUTIONS

### Contribution

E.C.S. designed and performed cellular immunology assays and flow cytometry, analyzed and interpreted the data and wrote the manuscript; B.A.L. designed and performed bioinformatics analysis, analyzed and interpreted the data and wrote the manuscript; A.P.B performed cell culture and in vivo experiments; B.C.T. performed cell culture experiments and FACS sorting; C.W. performed FACS sorting; R.A.F. and M.J.U. performed CRISPR/Cas9-mediated genome editing of target cells; N.L., Y.N., and R.M. assisted with sample acquisition and interpreted the data and Y.N. performed in vivo experiments; P.P.C. performed cell culture experiments; A.J.G. and R.B. designed experiments and interpreted the data; A.J.G. obtained funding; A.M.C. designed experiments, interpreted the data, obtained funding, wrote the manuscript, and co-supervised the study; M.G.R. designed experiments, interpreted the data, wrote the manuscript, obtained funding, and supervised the study.

### Lead Author

Maria Grazia Roncarolo, Department of Pediatrics, Stanford University School of Medicine, Lorry I. Lokey Stem Cell Research Building, 265 Campus Drive West, Room G3021A Stanford, California 94305-5461;

## Supplementary figure legends

**Figure S1. Flow cytometry-based killing assay. A.** Killing assay gating strategy, depicting representative dot plots of effector (CD4^IL10^ cells) and target cells (AML cells) after 3-day co-culture at 1:1 ratio. CountBright beads (Thermo Fisher) are added to each sample, and cells quantified according to manufacturer’s instructions. In the top-left dot plot, beads can be seen on the upper left corner (9.56%). After gating for single and live cells, CD45^+^CD33^+^ cells are considered to be AML cells, and NGFR^+^ cells T cells (CD4^IL10^ cells are engineered to express truncated NGFR marker gene). **B.** Killing assay results, representative flow cytometry plots. Raw cell counts recorded in AML gate are displayed on each dot plot; for elimination efficiency calculations (**Methods),** raw cell counts are adjusted using the number of added and recorded CountBright beads.

**Figure S2. CD4^IL10^ cell phenotype.** At the end of the 14-day expansion, CD4^IL10^ cells were stained with two separate panels for characteristic CD4^IL10^ cell surface proteins^5–8^. **A.** The gating strategy. **B.** Histograms showing surface protein expression on CD4^IL10^ cells from donor 2441 and **C.** from donor 072, both gated through CD3^+^CD4^+^ NGFR^+^ live single cells

**Figure S3. Single-cell RNA-seq experiment design and sorting of AML and CD4^IL10^ cells. A.** Design of the scRNA-seq experiment. Purified live CD4^IL10^ cells, sensitive AML, and resistant AML samples (n = 2 for each) were analyzed by scRNA-seq first at day 0 (at thaw for AML, at the end of expansion cycle for CD4^IL10^ cells), and then after 24h co-culture in 1:1 ratio; co-cultured cells were again purified for live cells before scRNA-seq using fluorescence-activated cell sorting (FACS). Data was generated in two independent experiments, each with all AML samples but with a different CD4^IL10^ donor. **B.** Flow cytometry plots of one resistant (top row; sample PATISD) and sensitive (bottom row; sample PARCEV) primary AML cell sample after co-culture with CD4^IL10^ cells for 24 hours at 1:1 ratio; gated through live single cells. Data shown are representative of two independent experiments.

**Figure S4. Identification of AML programs in scRNA-seq data.** UMAP plots of AML single cells per AML patient and timepoint, colored by cluster (**Methods**). **B.** Cophenetic coefficients used for selecting the number of AML programs. The highest number of AML programs (5) for which the cophenetic coefficient was >0.99 was selected. **C.** Heatmap depicting the 5 initial AML programs identified, Rows - genes, columns - average expression of genes in sample-specific clusters, and colors - the relative expression of genes. **D.** The proportion of AML single cells assigned to each AML program, grouped by their mitochondrial read content. **E.** UMAP plots of AML single cells per AML patient and timepoint, colored by AML program (**Methods**).

**Figure S5. Analysis of AML transcriptional programs in bulk RNA-seq data, related to Figure 3. A.** Heatmap depicting the expression of AML program markers across hematopoietic cell lines, grouped by the dominant program (**Supplemental Methods**). **B.** Heatmap of the AML program abundances across cell lines from hematopoietic cancers. Rows – cell lines, columns – AML programs, colors – program abundance. **C.** The abundance of AML programs in selected cell lines. **D.** Heatmap depicting the expression of AML program markers across samples from the TARGET cohort, grouped by the dominant program.

**Figure S6. Gene-set enrichment of CD4^IL10^ signatures with CD4 T cell subtypes, related to Figure 4. A.** Gene-set enrichment of the top 50 genes over-expressed in CD4^IL10^ after co-culture with sensitive and resistant AML, in gene expression signatures derived from naïve, resting memory, and activated memory CD4^+^ T cells (**Supplemental Methods**). **B-C.** Bubble plots representing the potential cell-cell interactions between AML cells in program P3 (**B**) and P4 (**C**) and CD4^IL10^ cells cultured with resistant AML (**Methods**). Bubble sizes are proportional to the average fraction of single cells in each population expressing the genes encoding the first and second interaction partners. Colors indicate the average log_2_ fold change of genes encoding the interaction partners in the indicted AML program relative to other AML cells and in the CD4^IL10^ cells after culture with resistant AML. **D-E.** Expression of *ITGB2* (CD18), *ITGAL* (CD11a) and *ICAM1* (CD54) in: (**D**) CD4^IL10^ single cells and (**E**) CD4^IL10^ cells after the expansion cycle (resting state) in an independent, previously published bulk RNA-seq dataset^7^. **F**. Expression of *ICAM1* in four primary AML samples analyzed by scRNA-seq across cells enriched in programs P1-P4.

**Figure S7. The effects of ICAM1 knock-out and ICAM1/LFA-1 interaction inhibitors on T cell degranulation and killing. A.** Flow cytometry surface staining of U937 cells (unstained: pink, wild-type: purple, knock-out: orange) for TNF-α receptor (encoded by *TNFRSF1B*), IFN-γ receptor (*IFNGR1*), CD18 subunit of LFA-1 (*ITGB2*), and CD54 adhesion protein (*ICAM1*), minimum 5 days post genome editing. HEK293FT cells, which do not express TNF-α receptor, were used as a negative control (green). **B.** Representative flow cytometry plots of CD8^+^ T cells (n = 7) co-cultured alone, with PMA+I or with K562, for 5hr to detect degranulation, expressed as % CD8^+^CD107a^+^ cells within live singlet CD3^+^ T cells. PMA+I (phorbol 12-myristate 13-acetate + ionomycin) was a positive control for degranulation. Summary of control conditions for degranulation are shown in the bottom-right insert. **C.** Representative FACS plots from one CD4^IL10^ donor co-cultured 24 hr with U937 WT cells at 1:1 ratio in the presence of indicated LFA-1 inhibitors at concentrations indicated on each plot; numbers indicate percentage in AML gate. DMSO alone is used as vehicle control. **D.** Representative FACS plots from one CD8^+^ T cell donor 24hr after interaction with indicated target cells. Activation was measured as percent CD69^+^CD25^+/-^ CD8^+^ T cells.

**Figure S8. Expression of ICAM1 and CD64 on cell lines and primary AML cells. A**. Flow cytometry surface staining for high-affinity Fc receptor CD64 and ICAM1 (encoding CD54) on U937 wild-type cells (U937), U937 ICAM1 knock out cells (CD54ko), and on four primary AML samples (adult, SU540 and SU555; pediatric, PAPVET and PATLIG), depicted as histograms of specific stainings (blue) overlayed over each cell type’s unstained control (red). Numbers on histograms indicate the geometric mean fluorescence intensity (gMFI) ratio between specific marker and unstained control. **B.** Cumulative gMFI values from A.

**Figure S9. Imaging of U937 *in vivo* tumor progression.** Images taken on days 5, 8 and 12 post PBS, U937-WT-Luc^+^ or U937-*ICAM1-KO*-Luc^+^ injections are shown.

