## Supplemental Figures and Tables for "AML/T cell interactomics uncover correlates of patient outcomes and the key role of ICAM1 in T cell killing of AML"

### Figure S1

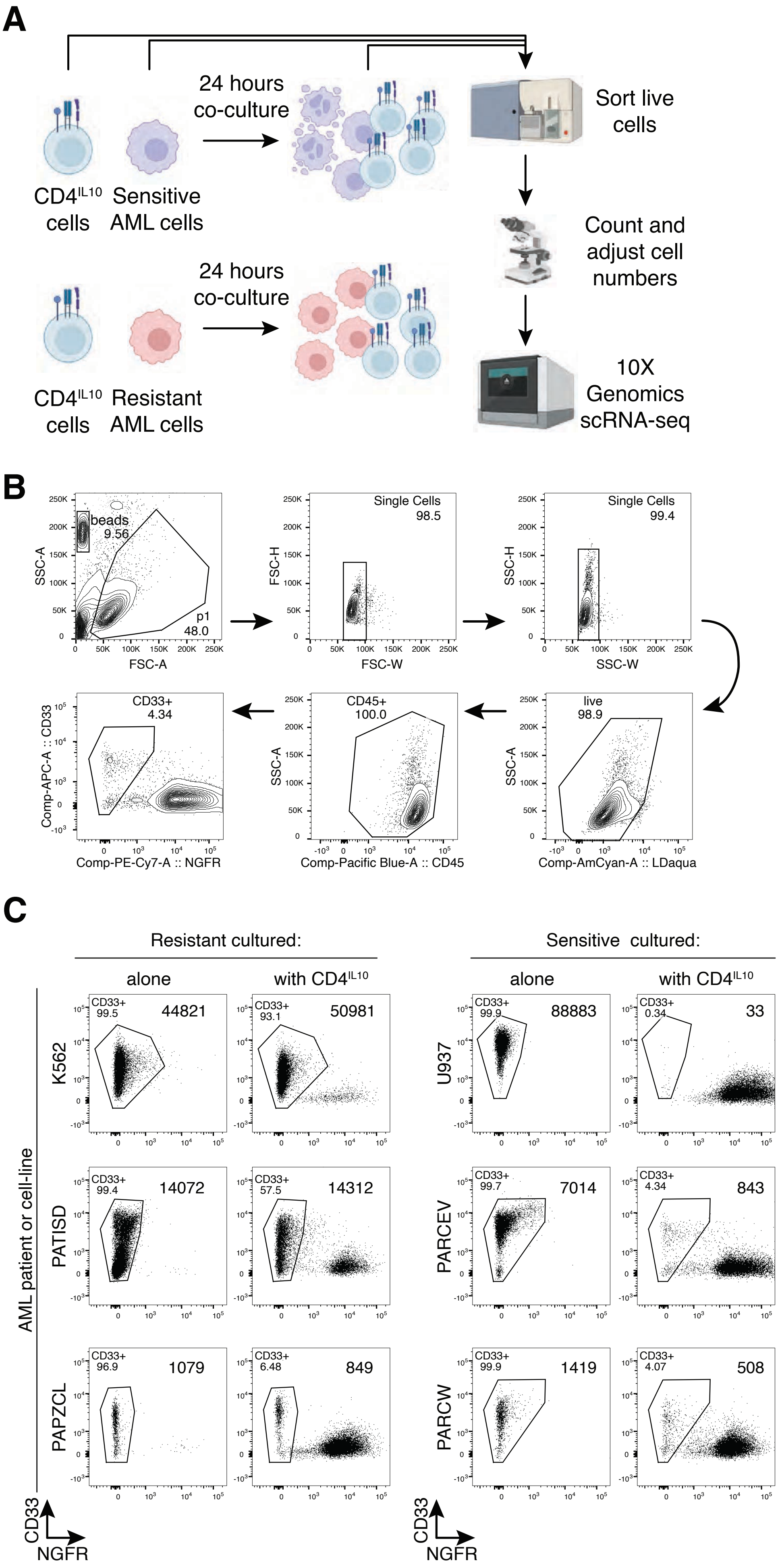
A
24 hours co-culture
Sort live cells
CD4IL10 cells
Sensitive AML cells
Count and adjust cell numbers
24 hours co-culture
10X Genomics scRNA-seq
CD4IL10 cells
Resistant AML cells
B
beads 9.56
p1 48.0
SSC-A
FSC-A
Single Cells 98.5
FSC-H
FSC-W
Single Cells 99.4
FSC-H
SSC-W
CD33+ 4.34
Comp-APC-A :: CD33
Comp-PE-Cy7-A :: NGFR
CD45+ 100.0
SSC-A
Comp-Pacific Blue-A :: CD45
live 98.9
SSC-A
Comp-AmCyan-A :: LDaqua
C
Resistant cultured:
Sensitive cultured:
alone
with CD4IL10
alone
with CD4IL10
AML patient or cell-line
K562
CD33+ 99.5
44821
CD33+ 93.1
50981
U937
CD33+ 99.9
88883
CD33+ 0.34
33
PATISD
CD33+ 99.4
14072
CD33+ 57.5
14312
PARCEV
CD33+ 99.7
7014
CD33+ 4.34
843
PAPZCL
CD33+ 96.9
1079
CD33+ 6.48
849
PARCW
CD33+ 99.9
1419
CD33+ 4.07
508
CD33
NGFR
CD33
NGFR

### Figure S2

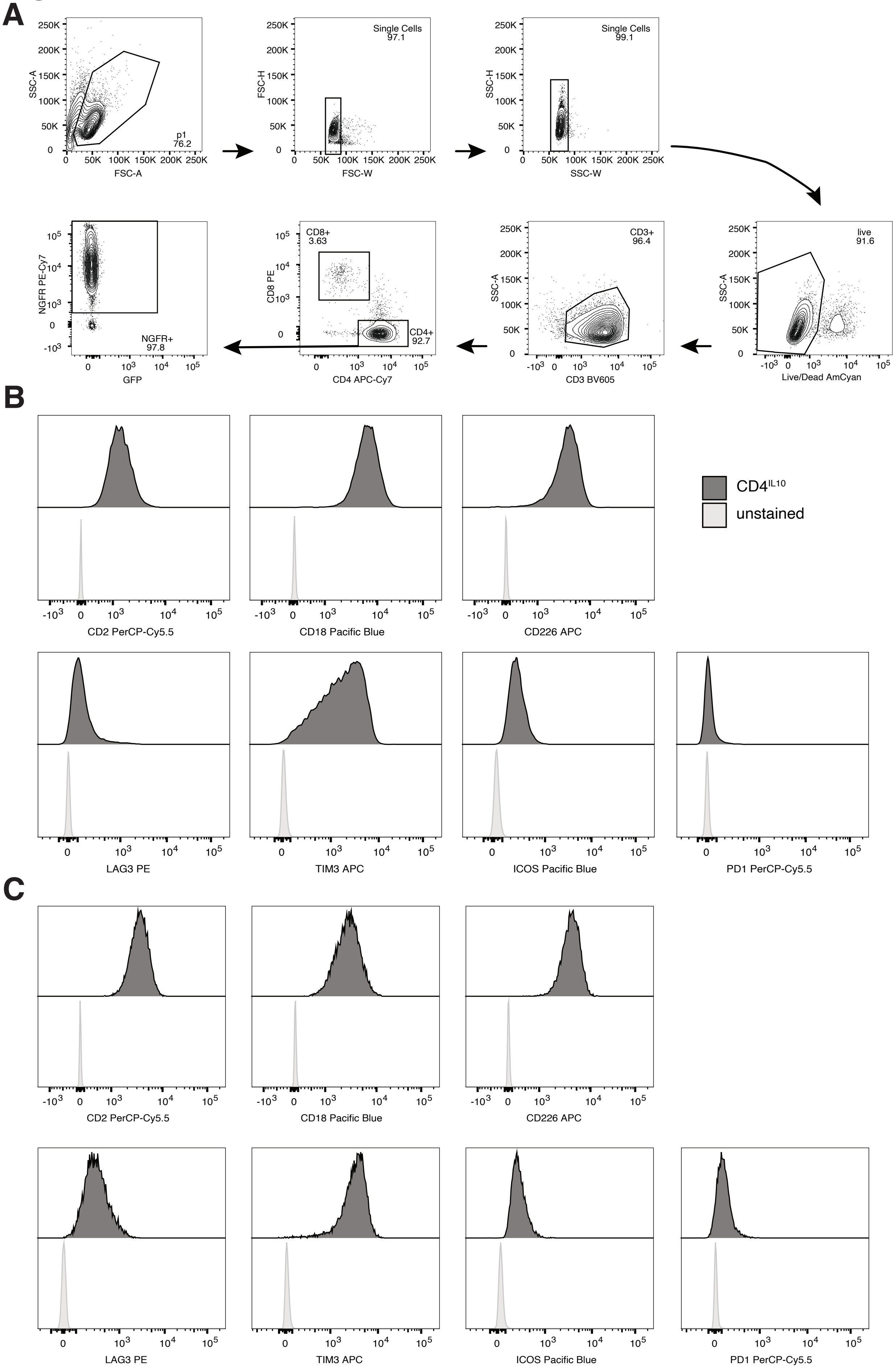

A
p1
76.2
Single Cells
97.1
Single Cells
99.1
SSC-A
FSC-A
FSC-H
FSC-W
SSC-H
SSC-W
live
91.6
Live/Dead AmCyan
CD3+
96.4
CD3 BV605
CD8+
3.63
CD4+
92.7
CD8 PE
CD4 APC-Cy7
NGFR+
97.8
NGFR PE-Cy7
GFP
B
CD4IL10
unstained
CD2 PerCP-Cy5.5
CD18 Pacific Blue
CD226 APC
LAG3 PE
TIM3 APC
ICOS Pacific Blue
PD1 PerCP-Cy5.5
C
CD2 PerCP-Cy5.5
CD18 Pacific Blue
CD226 APC
LAG3 PE
TIM3 APC
ICOS Pacific Blue
PD1 PerCP-Cy5.5

### Figure S3

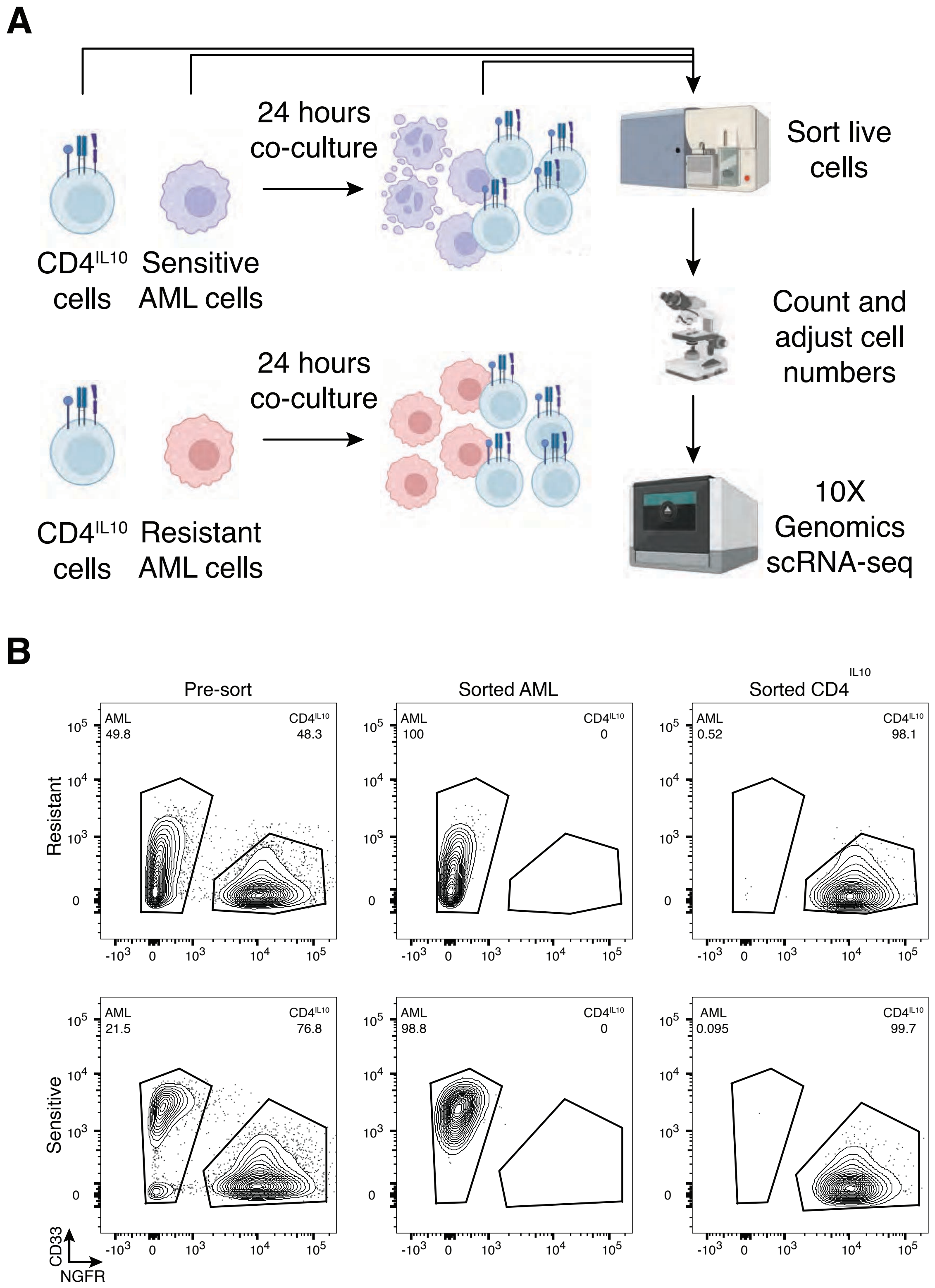
A
24 hours co-culture
Sort live cells
CD4^IL10 cells
Sensitive AML cells
Count and adjust cell numbers
24 hours co-culture
CD4^IL10 cells
Resistant AML cells
10X Genomics scRNA-seq
B
Pre-sort
Sorted AML
Sorted CD4^IL10
Resistant
Sensitive
AML 49.8
CD4^IL10 48.3
AML 100
CD4^IL10 0
AML 0.52
CD4^IL10 98.1
AML 21.5
CD4^IL10 76.8
AML 98.8
CD4^IL10 0
AML 0.095
CD4^IL10 99.7
CD33
NGFR

### Figure S4

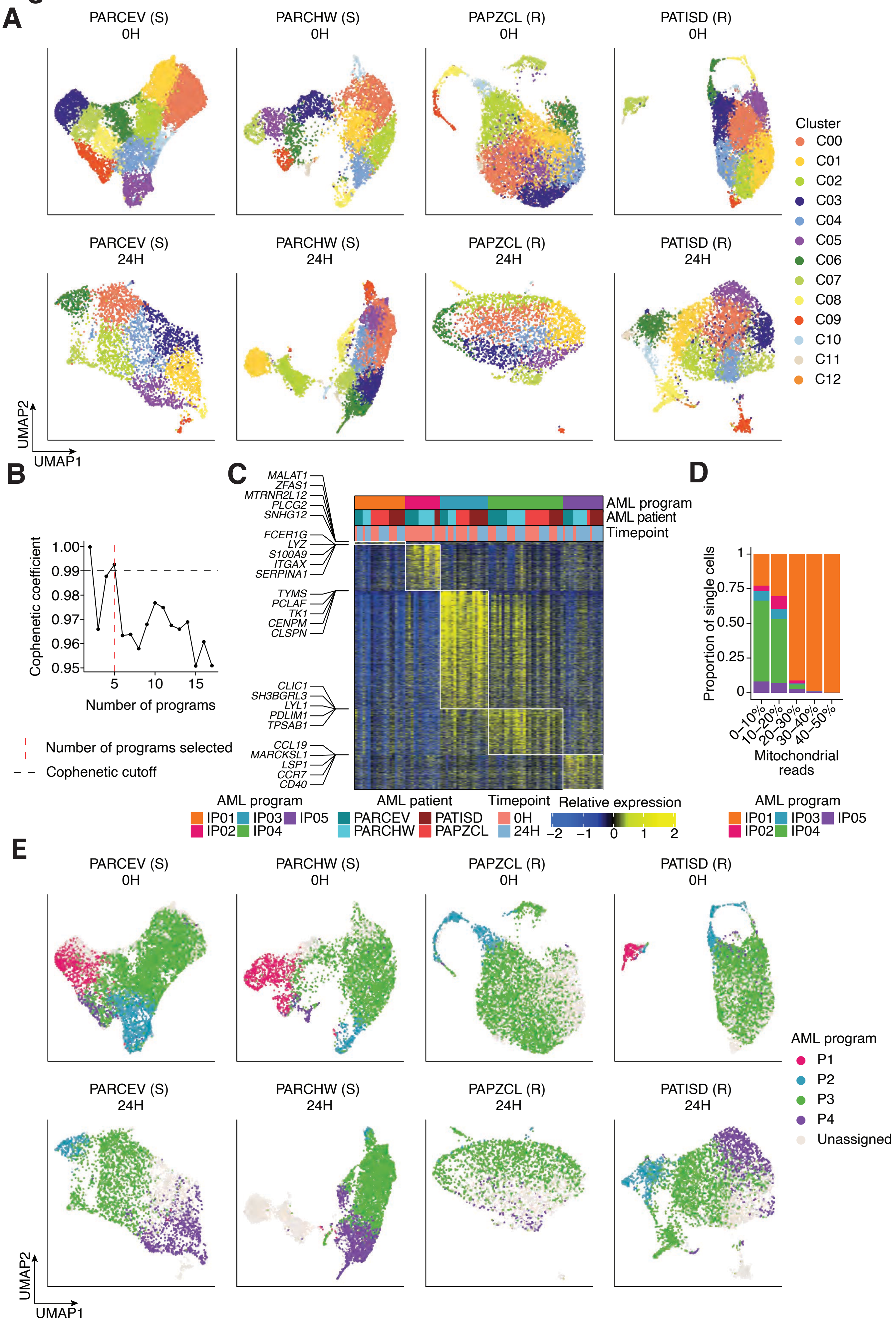
A
PARCEV (S) 0H
PARCHW (S) 0H
PAPZCL (R) 0H
PATISD (R) 0H
PARCEV (S) 24H
PARCHW (S) 24H
PAPZCL (R) 24H
PATISD (R) 24H
Cluster
C00
C01
C02
C03
C04
C05
C06
C07
C08
C09
C10
C11
C12
UMAP2
UMAP1
B
Cophenetic coefficient
Number of programs
Number of programs selected
Cophenetic cutoff
C
MALAT1
ZFAS1
MTRNR2L12
PLCG2
SNHG12
FCER1G
LYZ
S100A9
ITGAX
SERPINA1
TYMS
PCLAF
TK1
CENPM
CLSPN
CLIC1
SH3BGRL3
LYL1
PDLIM1
TPSAB1
CCL19
MARCKSL1
LSP1
CCR7
CD40
AML program
AML patient
Timepoint
IP01 IP02 IP03 IP04 IP05
PARCEV PARCHW PATISD PAPZCL
0H 24H
Relative expression
D
Proportion of single cells
Mitochondrial reads
0–10% 10–20% 20–30% 30–40% 40–50%
E
AML program
P1
P2
P3
P4
Unassigned

### Figure S5

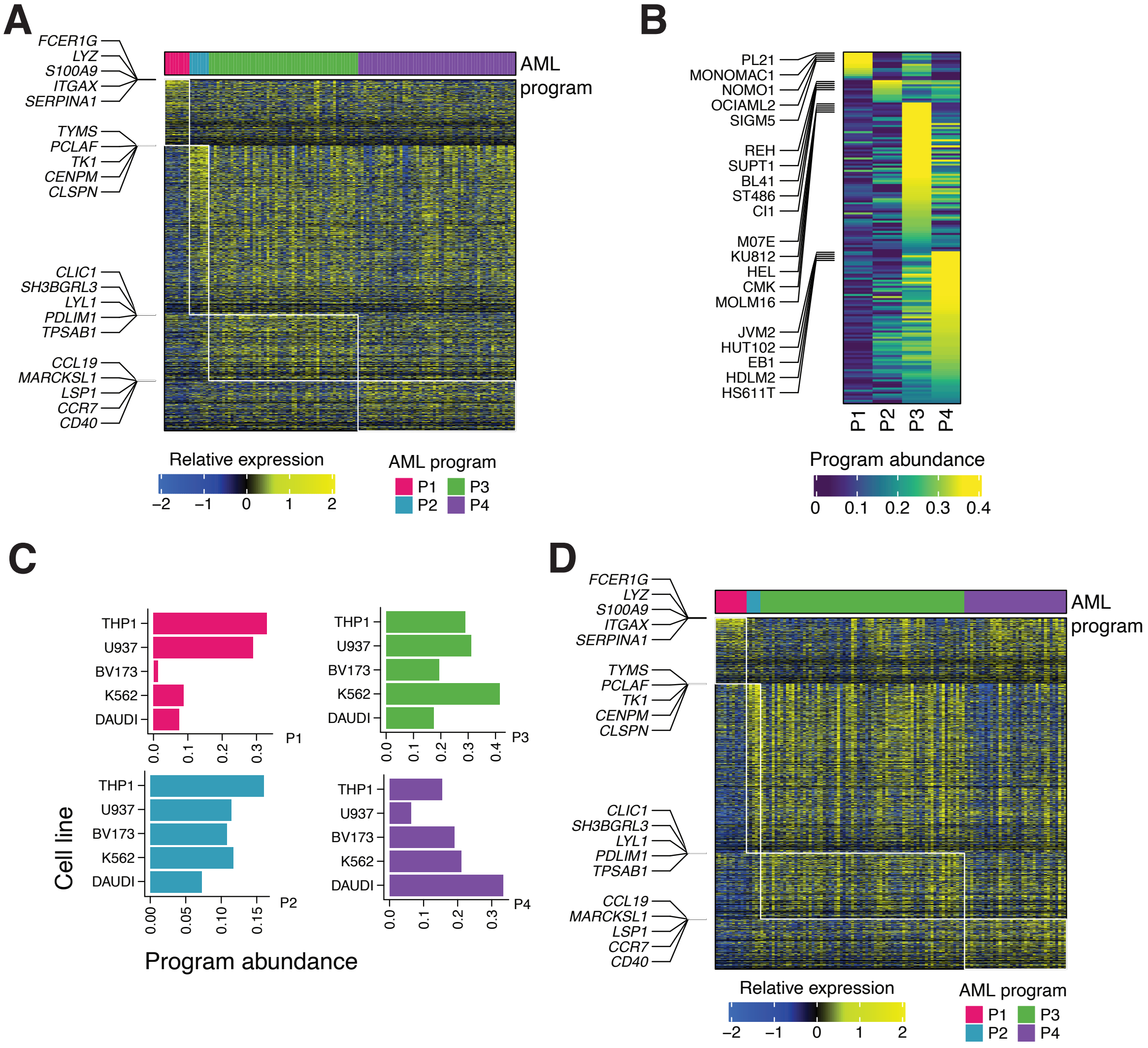

A
FCER1G
LYZ
S100A9
ITGAX
SERPINA1
TYMS
PCLAF
TK1
CENPM
CLSPN
CLIC1
SH3BGRL3
LYL1
PDLIM1
TPSAB1
CCL19
MARCKSL1
LSP1
CCR7
CD40
AML program
Relative expression
−2 −1 0 1 2
AML program
P1 P3
P2 P4
B
PL21
MONOMAC1
NOMO1
OCIAML2
SIGM5
REH
SUPT1
BL41
ST486
CI1
M07E
KU812
HEL
CMK
MOLM16
JVM2
HUT102
EB1
HDLM2
HS611T
P1 P2 P3 P4
Program abundance
0 0.1 0.2 0.3 0.4
C
THP1
U937
BV173
K562
DAUDI
P1
P2
P3
P4
0.0 0.1 0.2 0.3
0.0 0.1 0.2 0.3 0.4
0.00 0.05 0.10 0.15
Cell line
Program abundance
D
FCER1G
LYZ
S100A9
ITGAX
SERPINA1
TYMS
PCLAF
TK1
CENPM
CLSPN
CLIC1
SH3BGRL3
LYL1
PDLIM1
TPSAB1
CCL19
MARCKSL1
LSP1
CCR7
CD40
AML program
Relative expression
−2 −1 0 1 2
AML program
P1 P3
P2 P4

#### Figure S6

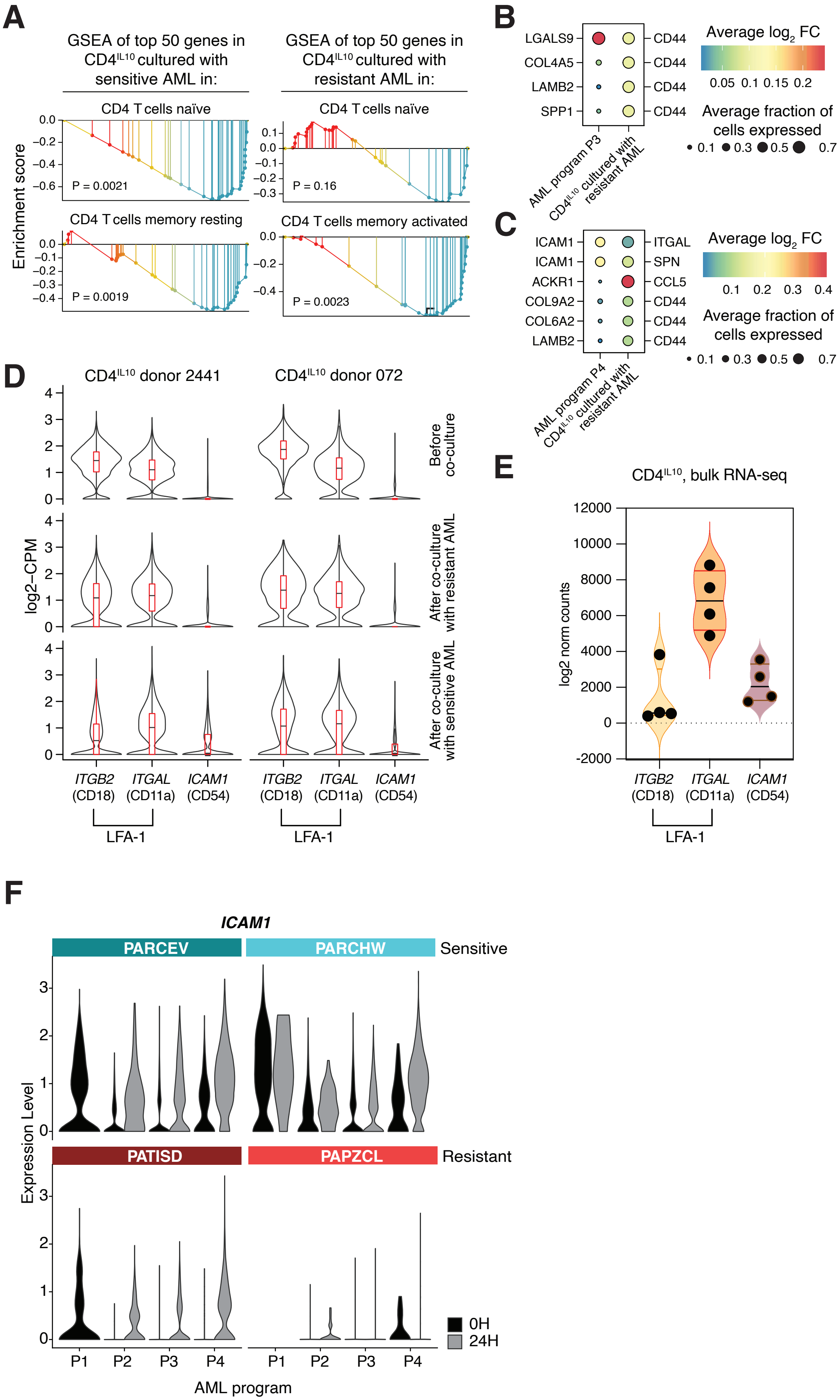

A
GSEA of top 50 genes in CD4IL10 cultured with sensitive AML in:
GSEA of top 50 genes in CD4IL10 cultured with resistant AML in:
CD4 T cells naïve
CD4 T cells memory resting
CD4 T cells memory activated
Enrichment score
P = 0.0021
P = 0.16
P = 0.0019
P = 0.0023
B
LGALS9
COL4A5
LAMB2
SPP1
CD44
AML program P3
CD4IL10 cultured with resistant AML
Average log2 FC
Average fraction of cells expressed
C
ICAM1
ACKR1
COL9A2
COL6A2
LAMB2
ITGAL
SPN
CCL5
CD44
AML program P4
D
CD4IL10 donor 2441
CD4IL10 donor 072
log2−CPM
Before co-culture
After co-culture with resistant AML
After co-culture with sensitive AML
ITGB2 (CD18)
ITGAL (CD11a)
ICAM1 (CD54)
LFA-1
E
CD4IL10, bulk RNA-seq
log2 norm counts
F
ICAM1
PARCEV
PARCHW
Sensitive
PATISD
PAPZCL
Resistant
Expression Level
0H
24H
P1
P2
P3
P4
AML program

### Figure S7

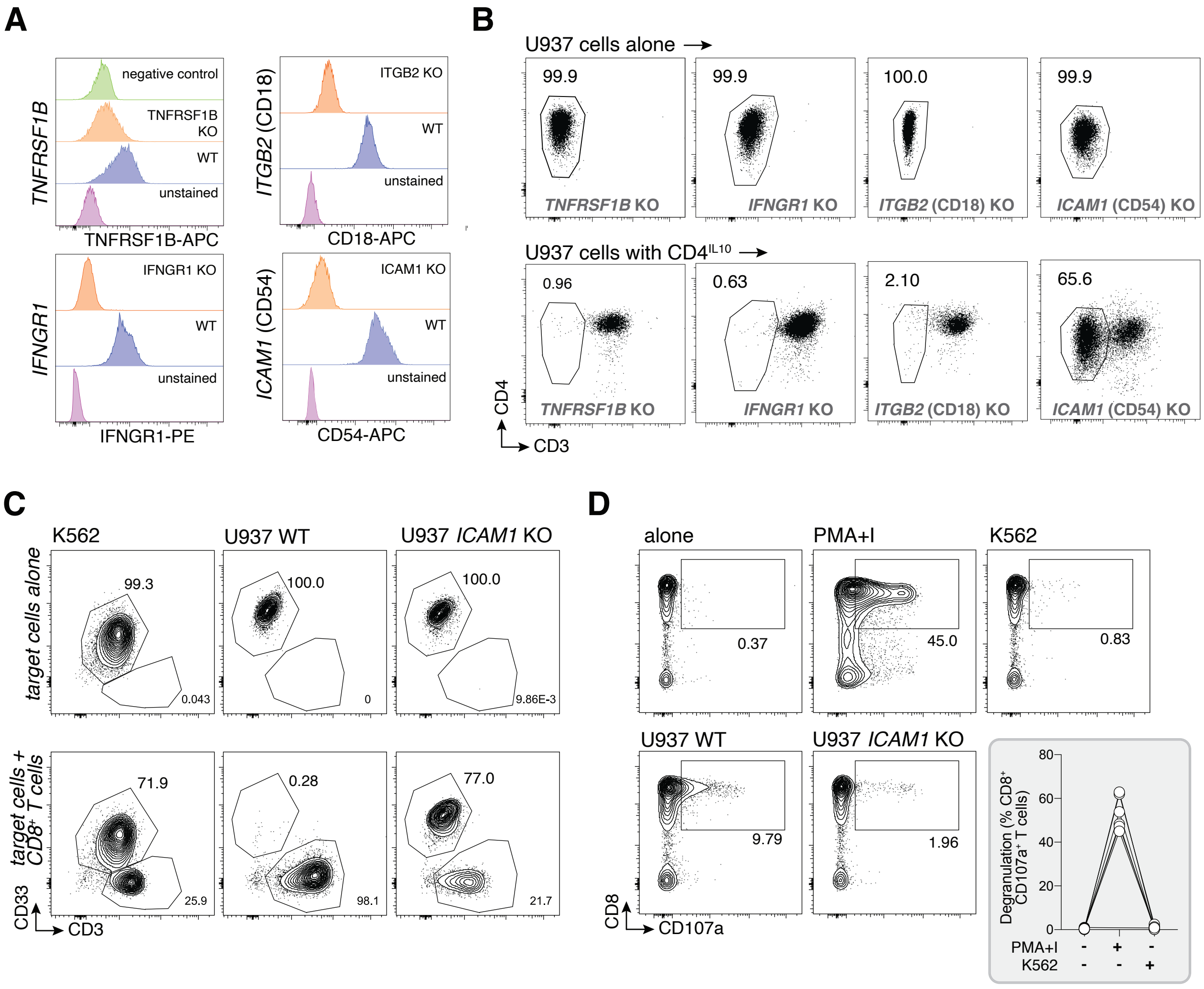
A
TNFRSF1B
negative control
TNFRSF1B KO
WT
unstained
TNFRSF1B-APC
ITGB2 (CD18)
ITGB2 KO
WT
unstained
CD18-APC
IFNGR1
IFNGR1 KO
WT
unstained
IFNGR1-PE
ICAM1 (CD54)
ICAM1 KO
WT
unstained
CD54-APC
B
U937 cells alone
99.9
99.9
100.0
99.9
TNFRSF1B KO
IFNGR1 KO
ITGB2 (CD18) KO
ICAM1 (CD54) KO
U937 cells with CD4IL10
0.96
0.63
2.10
65.6
TNFRSF1B KO
IFNGR1 KO
ITGB2 (CD18) KO
ICAM1 (CD54) KO
CD4
CD3
C
K562
U937 WT
U937 ICAM1 KO
target cells alone
99.3
0.043
100.0
0
100.0
9.86E-3
target cells + CD8+ T cells
71.9
25.9
0.28
98.1
77.0
21.7
CD33
CD3
D
alone
PMA+I
K562
0.37
45.0
0.83
U937 WT
U937 ICAM1 KO
9.79
1.96
CD8
CD107a
Degranulation (% CD8+ CD107a+ T cells)
80
60
40
20
0
PMA+I
K562
-
+
-
-
-
+

#### Figure S8

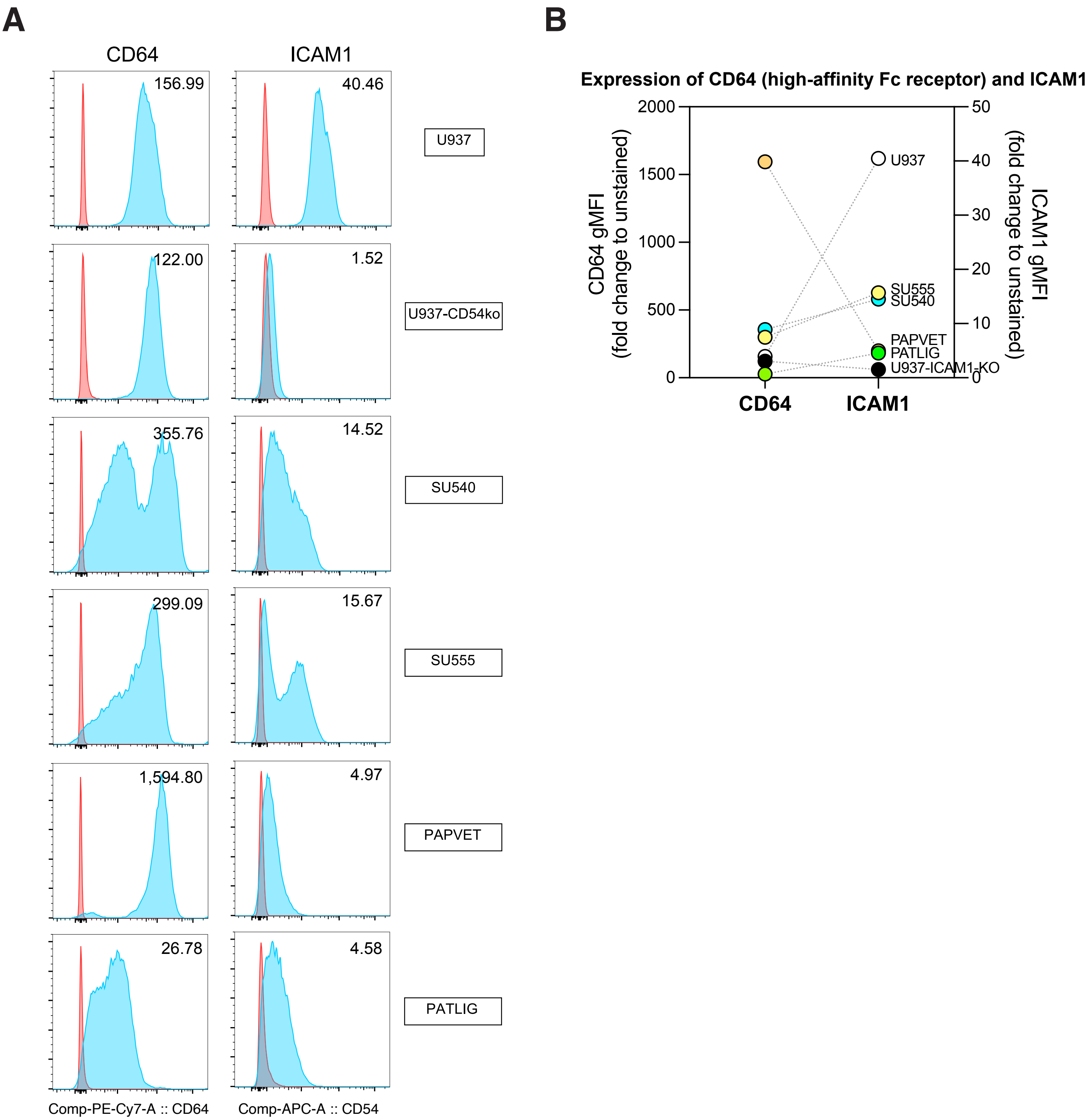
A
CD64
ICAM1
156.99
40.46
U937
122.00
1.52
U937-CD54ko
355.76
14.52
SU540
299.09
15.67
SU555
1,594.80
4.97
PAPVET
26.78
4.58
PATLIG
Comp-PE-Cy7-A :: CD64
Comp-APC-A :: CD54
B
Expression of CD64 (high-affinity Fc receptor) and ICAM1
CD64 gMFI (fold change to unstained)
ICAM1 gMFI (fold change to unstained)
2000
1500
1000
500
0
50
40
30
20
10
0
U937
SU555
SU540
PAPVET
PATLIG
U937-ICAM1-KO
CD64
ICAM1

### Figure S9

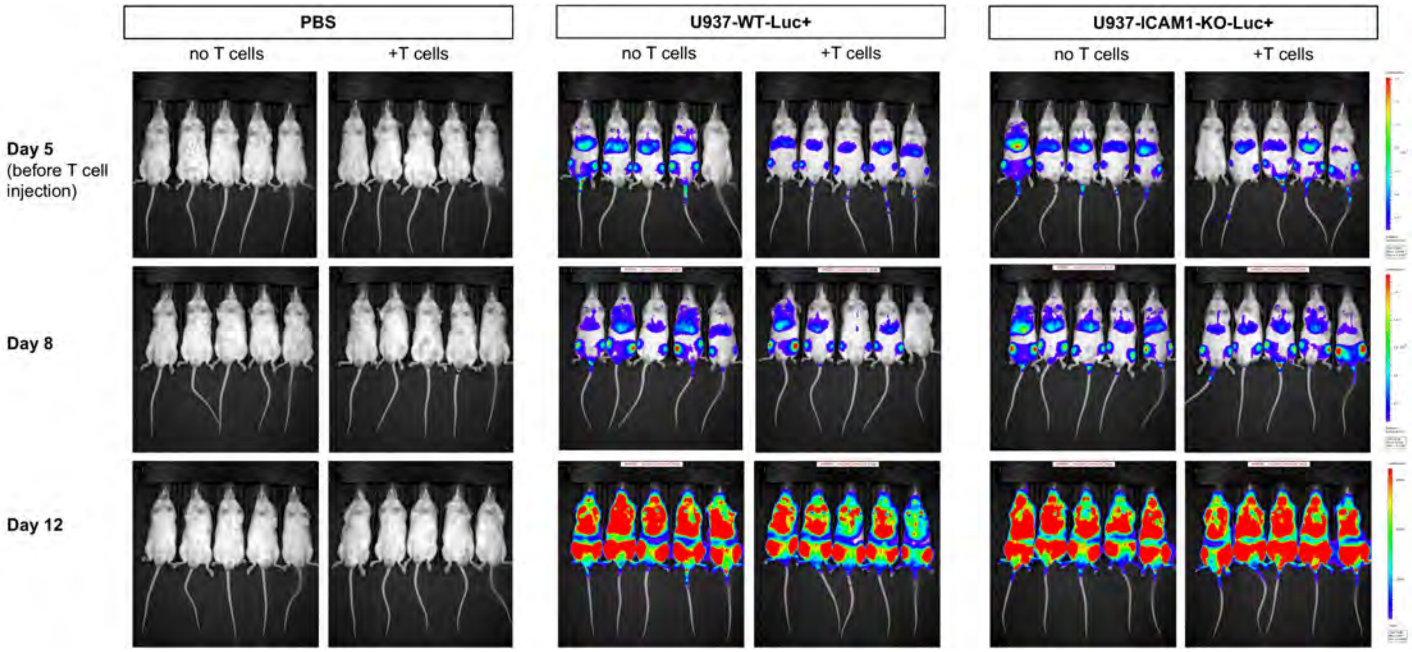

PBS
U937-WT-Luc+
U937-ICAM1-KO-Luc+
no T cells
+T cells
no T cells
+T cells
no T cells
+T cells
Day 5
(before T cell injection)
Day 8
Day 12

**Table S1.** Clinical data related to primary AML patient samples

| AML patient | Response to $CD4^{IL10}$ killing | FAB subtype | WHO classification | Cytogenetics onset | Blasts in bone marrow (%) | Age at diagnosis (days) | Risk group |
|---|---|---|---|---|---|---|---|
| PARCEV | Sensitive | M4 | acute myelomonocytic leukemia | 47,XX,+8(12) | 94 | 4152 | High |
| PARCHW | Sensitive | Unknown | AML with CBFB::MYH11 fusion | 46,XY,inv(16)(p13.1q22)(17)/46,XY(3) | 80 | 4209 | Low |
| PATISD | Resistant | NOS | AML, myelodysplasia-related | 45,XY,t(3;3)(q21;q26),-7(20) | 91 | 3901 | High |
| PAPZCL | Resistant | M0 | AML, myelodysplasia-related | 46,XY,del(5)(q13q31)(13)/46,XY(7) | 97 | 5439 | High |

Legend: AML = acute myeloid leukemia; FAB = French-American-British classification of AML; WHO = World Health Organization classification of AML, 2022 update

**Table S2.** List of antibodies used.

| Antibody | Clone | Company | Dilution | Panel | | |
|---|---|---|---|---|---|---|
| | | | | phenotyping | killing assay | sorting |
| CD2-PerCP-Cy5.5 | RPA-2.10 | Biolegend | 1:50 | + | | |
| CD3-BV605 or PerCP-Cy5.5 | OKT3 | Biolegend | 1:100 | + | + | |
| CD4-APC-Cy7 | RPA-T4 | Biolegend | 1:100 | + | + | |
| CD8-PE-Cy7 | SK1 | BD Biosciences | 1:100 | | + | |
| CD8-PE | SK1 | BD Biosciences | 1:100 | + | | |
| CD18-BV421 or APC | 6.7 | BD Biosciences | 1:50 | + | | |
| CD25-AF700 | BC96 | Biolegend | 1:100 | + | | |
| CD33-APC | AC104.3E3 | Miltenyi Biotec | 1:50 | | + | + |
| CD45-Pacific Blue | HI100 | Biolegend | 1:100 | | + | |
| CD54-APC | 15.2 | Tonbo Biosciences | 1:50 | + | | |
| CD64-PE-Cy7 | 10.1 | BD Biosciences | 1:50 | + | | |
| CD226-APC | 11A8 | Biolegend | 1:50 | + | | |
| CD271 (NGFR)-PE-Cy7 | ME20.4 | Biolegend | 1:100 | + | + | + |
| LAG3-PE | REA351 | Miltenyi Biotec | 1:33 | + | | |
| TIM3-AF647 | 7D3 | BD Biosciences | 1:50 | + | | |
| ICOS-Pacific Blue | C398.4A | Biolegend | 1:50 | + | | |
| PD1-PerCP-Cy5.5 | EH12.2H7 | Biolegend | 1:50 | + | | |
| Ghost dye violet 510 live/dead dye | | Tonbo Biosciences | 1:250 | + | + | + |
| IFNGR1-PE | GIR-208 | Biolegend | 1:50 | + | | |
| TNFRSF1B-APC | 3G7A02 | Biolegend | 1:50 | + | | |
| CD69-BV605 | FN50 | Biolegend | 1:100 | + | | |

**Table S3.** Genes differentially expressed in AML programs, related to **Figure 1**.

**Gene symbol** -= The symbol of gene for which the differential expression is assessed.
**AML program** = The AML program in which differential expression is assessed.
**Average log2 fold change** =
The weighted average LOG2 fold change of gene in cells assigned to The program indicated in column B relative to all The other cells, across all AML patient/timepoint pairs.
**Avg. percent of cells assigned to program with CPM > 0** - The weighted average percentage of cells assigned to the program with CPM > 0, across all AML patient/timepoint pairs.
**Avg. percent of cells not assigned to program with CPM > 0** =
The weighted average percentage of cells not assigned to the program with CPM > 0, across all AML patient/timepoint pairs.
**DE meta z-score** =
The meta z-score obtained by aggregating individual z-score statistics of differential expression (DE), across all AML patient/timepoint pairs, using Liptak's method (Methods).
**DE *p*-value** - The value obtained by converting the value in column **DE meta z-score** to a two-sided *p*-value.
**DE *q*-value** - The *q*-value corresponding to the *p*-value in column **DE *p*-value** after adjusted for multiple hypothesis testing using Benjamini-Hochberg method.
**Suface protein** - Flag indicating whether the gene is a surface protein, as annotated in Cell Surface Protein Atlas (*Bausch-Fluck et al. 2018*).

Note: For columns C-F, the the natural log of the number of cells assigned to the program was used as weight.

| Gene symbol | AML program | Average $log_2$ fold change | Avg. percent of cells assigned to program with CPM > 0 | Avg. percent of cells not assigned to program with CPM > 0 | DE meta z-score | DE *p*-value | DE *q*-value | Surface protein |
|---|---|---|---|---|---|---|---|---|
| *CD68* | P1 | 1.15 | 0.91 | 0.38 | 37.93 | 0.000 | 0.000 | TRUE |
| *TREM1* | P1 | 0.93 | 0.75 | 0.17 | 36.37 | 0.000 | 0.000 | TRUE |
| *GPR183* | P1 | 0.92 | 0.80 | 0.39 | 28.73 | 0.000 | 0.000 | TRUE |
| *PLAUR* | P1 | 0.91 | 0.87 | 0.59 | 26.50 | 0.000 | 0.000 | TRUE |
| *EVI2B* | P1 | 0.90 | 0.89 | 0.64 | 31.40 | 0.000 | 0.000 | TRUE |
| *FCGRT* | P1 | 0.84 | 0.82 | 0.33 | 31.53 | 0.000 | 0.000 | TRUE |
| *EMP3* | P1 | 0.84 | 0.99 | 0.88 | 27.69 | 0.000 | 0.000 | TRUE |
| *TNFRSF1B* | P1 | 0.82 | 0.65 | 0.25 | 22.74 | 0.000 | 0.000 | TRUE |
| *ITGB2* | P1 | 0.81 | 0.90 | 0.61 | 30.09 | 0.000 | 0.000 | TRUE |
| *MS4A6A* | P1 | 0.80 | 0.60 | 0.08 | 32.06 | 0.000 | 0.000 | TRUE |
| *CYBB* | P1 | 0.79 | 0.62 | 0.09 | 32.11 | 0.000 | 0.000 | TRUE |
| *CD83* | P1 | 0.79 | 0.85 | 0.63 | 24.62 | 0.000 | 0.000 | TRUE |
| *CLEC7A* | P1 | 0.77 | 0.55 | 0.08 | 30.46 | 0.000 | 0.000 | TRUE |
| *ITGAX* | P1 | 0.68 | 0.58 | 0.10 | 29.07 | 0.000 | 0.000 | TRUE |
| *EMP1* | P1 | 0.67 | 0.83 | 0.66 | 20.92 | 0.000 | 0.000 | TRUE |
| *CD44* | P1 | 0.67 | 0.99 | 0.92 | 20.86 | 0.000 | 0.000 | TRUE |
| *IL10RA* | P1 | 0.67 | 0.69 | 0.20 | 29.64 | 0.000 | 0.000 | TRUE |
| *MPEG1* | P1 | 0.66 | 0.57 | 0.06 | 32.97 | 0.000 | 0.000 | TRUE |
| *APLP2* | P1 | 0.66 | 0.93 | 0.82 | 23.09 | 0.000 | 0.000 | TRUE |
| *CD14* | P1 | 0.60 | 0.43 | 0.02 | 28.44 | 0.000 | 0.000 | TRUE |
| *PTAFR* | P1 | 0.57 | 0.59 | 0.16 | 27.27 | 0.000 | 0.000 | TRUE |
| *HBEGF* | P1 | 0.55 | 0.72 | 0.43 | 20.81 | 0.000 | 0.000 | TRUE |
| *HCAR3* | P1 | 0.54 | 0.57 | 0.15 | 23.87 | 0.000 | 0.000 | TRUE |
| *SLC11A1* | P1 | 0.54 | 0.45 | 0.06 | 26.04 | 0.000 | 0.000 | TRUE |
| *ICAM1* | P1 | 0.53 | 0.63 | 0.35 | 20.09 | 0.000 | 0.000 | TRUE |
| *TLR2* | P1 | 0.52 | 0.55 | 0.16 | 25.33 | 0.000 | 0.000 | TRUE |
| *CPD* | P1 | 0.52 | 0.69 | 0.34 | 21.43 | 0.000 | 0.000 | TRUE |
| *ATP13A3* | P1 | 0.51 | 0.72 | 0.51 | 16.97 | 0.000 | 0.000 | TRUE |
| *CD93* | P1 | 0.49 | 0.44 | 0.06 | 24.33 | 0.000 | 0.000 | TRUE |
| *PILRA* | P1 | 0.48 | 0.50 | 0.06 | 27.20 | 0.000 | 0.000 | TRUE |
| *RELT* | P1 | 0.46 | 0.64 | 0.34 | 21.45 | 0.000 | 0.000 | TRUE |
| *IFNGR2* | P1 | 0.46 | 0.63 | 0.36 | 20.09 | 0.000 | 0.000 | TRUE |
| *LRRC25* | P1 | 0.46 | 0.48 | 0.04 | 28.31 | 0.000 | 0.000 | TRUE |
| *C5AR1* | P1 | 0.45 | 0.32 | 0.01 | 23.68 | 0.000 | 0.000 | TRUE |
| *ITGA5* | P1 | 0.44 | 0.70 | 0.45 | 19.40 | 0.000 | 0.000 | TRUE |
| *IGSF6* | P1 | 0.44 | 0.46 | 0.09 | 21.03 | 0.000 | 0.000 | TRUE |
| *CD300E* | P1 | 0.44 | 0.31 | 0.01 | 24.52 | 0.000 | 0.000 | TRUE |
| *ADAM17* | P1 | 0.44 | 0.77 | 0.60 | 17.67 | 0.000 | 0.000 | TRUE |
| *SLC15A3* | P1 | 0.43 | 0.46 | 0.05 | 25.21 | 0.000 | 0.000 | TRUE |
| *EVI2A* | P1 | 0.43 | 0.61 | 0.34 | 17.16 | 0.000 | 0.000 | TRUE |
| *P2RX4* | P1 | 0.42 | 0.61 | 0.35 | 17.76 | 0.000 | 0.000 | TRUE |
| *CD48* | P1 | 0.40 | 0.53 | 0.38 | 12.58 | 0.000 | 0.000 | TRUE |
| *IL13RA1* | P1 | 0.39 | 0.50 | 0.15 | 20.60 | 0.000 | 0.000 | TRUE |
| *IL17RA* | P1 | 0.39 | 0.55 | 0.23 | 18.73 | 0.000 | 0.000 | TRUE |
| *SLC2A3* | P1 | 0.39 | 0.90 | 0.81 | 9.88 | 0.000 | 0.000 | TRUE |
| *CXCL16* | P1 | 0.39 | 0.38 | 0.05 | 21.26 | 0.000 | 0.000 | TRUE |
| *HLA-B* | P1 | 0.39 | 1.00 | 0.99 | 22.90 | 0.000 | 0.000 | TRUE |
| *CD36* | P1 | 0.39 | 0.43 | 0.10 | 23.49 | 0.000 | 0.000 | TRUE |
| *EREG* | P1 | 0.39 | 0.22 | 0.03 | 14.74 | 0.000 | 0.000 | TRUE |
| *TLR4* | P1 | 0.38 | 0.38 | 0.03 | 25.08 | 0.000 | 0.000 | TRUE |
| *MYADM* | P1 | 0.37 | 0.87 | 0.86 | 17.46 | 0.000 | 0.000 | TRUE |
| *ABCA1* | P1 | 0.37 | 0.49 | 0.25 | 18.47 | 0.000 | 0.000 | TRUE |
| *PRNP* | P1 | 0.36 | 0.71 | 0.53 | 17.48 | 0.000 | 0.000 | TRUE |
| *CD302* | P1 | 0.35 | 0.69 | 0.58 | 15.95 | 0.000 | 0.000 | TRUE |
| *FCGR2A* | P1 | 0.35 | 0.45 | 0.18 | 18.83 | 0.000 | 0.000 | TRUE |
| *PLXDC2* | P1 | 0.35 | 0.51 | 0.24 | 18.06 | 0.000 | 0.000 | TRUE |
| *LILRB2* | P1 | 0.35 | 0.34 | 0.02 | 22.92 | 0.000 | 0.000 | TRUE |
| *CD53* | P1 | 0.35 | 0.73 | 0.61 | 12.87 | 0.000 | 0.000 | TRUE |
| *SLCO3A1* | P1 | 0.35 | 0.56 | 0.36 | 17.21 | 0.000 | 0.000 | TRUE |
| *CCR1* | P1 | 0.34 | 0.35 | 0.02 | 23.38 | 0.000 | 0.000 | TRUE |
| *SLC43A2* | P1 | 0.34 | 0.40 | 0.15 | 16.51 | 0.000 | 0.000 | TRUE |
| *TCIRG1* | P1 | 0.34 | 0.54 | 0.28 | 15.37 | 0.000 | 0.000 | TRUE |
| *ITGAM* | P1 | 0.32 | 0.35 | 0.07 | 21.41 | 0.000 | 0.000 | TRUE |
| *CLEC5A* | P1 | 0.32 | 0.29 | 0.04 | 20.14 | 0.000 | 0.000 | TRUE |
| *LRP1* | P1 | 0.31 | 0.30 | 0.01 | 23.58 | 0.000 | 0.000 | TRUE |
| *ANPEP* | P1 | 0.31 | 0.47 | 0.38 | 2.47 | 0.013 | 0.030 | TRUE |
| *CD55* | P1 | 0.30 | 0.66 | 0.57 | 15.53 | 0.000 | 0.000 | TRUE |
| *HCAR2* | P1 | 0.30 | 0.41 | 0.12 | 18.48 | 0.000 | 0.000 | TRUE |
| *PTGER2* | P1 | 0.30 | 0.45 | 0.29 | 12.29 | 0.000 | 0.000 | TRUE |
| *CSF2RA* | P1 | 0.29 | 0.44 | 0.15 | 17.58 | 0.000 | 0.000 | TRUE |
| *SIRPA* | P1 | 0.29 | 0.40 | 0.13 | 18.75 | 0.000 | 0.000 | TRUE |
| *STAB1* | P1 | 0.29 | 0.44 | 0.23 | 16.19 | 0.000 | 0.000 | TRUE |
| *QSOX1* | P1 | 0.29 | 0.53 | 0.34 | 13.88 | 0.000 | 0.000 | TRUE |
| *ATP1A1* | P1 | 0.28 | 0.63 | 0.46 | 13.07 | 0.000 | 0.000 | TRUE |
| *CLEC12A* | P1 | 0.28 | 0.50 | 0.34 | 13.51 | 0.000 | 0.000 | TRUE |
| *LDLR* | P1 | 0.28 | 0.68 | 0.63 | 11.81 | 0.000 | 0.000 | TRUE |
| *ADAM9* | P1 | 0.28 | 0.49 | 0.26 | 13.25 | 0.000 | 0.000 | TRUE |
| *FCAR* | P1 | 0.27 | 0.28 | 0.01 | 22.68 | 0.000 | 0.000 | TRUE |
| *BST1* | P1 | 0.27 | 0.39 | 0.18 | 15.40 | 0.000 | 0.000 | TRUE |
| *ITGAL* | P1 | 0.27 | 0.46 | 0.25 | 10.97 | 0.000 | 0.000 | TRUE |

| | | | | | | | | |
|---|---|---|---|---|---|---|---|---|
| *CD86* | P1 | 0.26 | 0.32 | 0.07 | 19.17 | 0.000 | 0.000 | TRUE |
| *TNFSF13B* | P1 | 0.25 | 0.61 | 0.51 | 9.11 | 0.000 | 0.000 | TRUE |
| *P2RX7* | P1 | 0.25 | 0.30 | 0.10 | 13.17 | 0.000 | 0.000 | TRUE |
| *LILRB1* | P1 | 0.25 | 0.26 | 0.02 | 16.48 | 0.000 | 0.000 | TRUE |
| *SLC6A6* | P1 | 0.25 | 0.51 | 0.36 | 10.51 | 0.000 | 0.000 | TRUE |
| *SLAMF8* | P1 | 0.25 | 0.26 | 0.02 | 5.94 | 0.000 | 0.000 | TRUE |
| *LAMP1* | P1 | 0.25 | 0.74 | 0.66 | 10.56 | 0.000 | 0.000 | TRUE |
| *FCGR1A* | P1 | 0.24 | 0.35 | 0.12 | 14.41 | 0.000 | 0.000 | TRUE |
| *IFNGR1* | P1 | 0.24 | 0.58 | 0.48 | 11.62 | 0.000 | 0.000 | TRUE |
| *CXCR4* | P1 | 0.23 | 0.50 | 0.39 | 8.64 | 0.000 | 0.000 | TRUE |
| *NRP1* | P1 | 0.23 | 0.23 | 0.07 | 4.78 | 0.000 | 0.000 | TRUE |
| *IL6R* | P1 | 0.23 | 0.41 | 0.21 | 13.12 | 0.000 | 0.000 | TRUE |
| *RNF149* | P1 | 0.22 | 0.46 | 0.35 | 6.71 | 0.000 | 0.000 | TRUE |
| *LILRB3* | P1 | 0.22 | 0.26 | 0.05 | 14.12 | 0.000 | 0.000 | TRUE |
| *CSF1R* | P1 | 0.22 | 0.30 | 0.09 | 15.63 | 0.000 | 0.000 | TRUE |
| *HM13* | P1 | 0.21 | 0.67 | 0.59 | 7.45 | 0.000 | 0.000 | TRUE |
| *PTPRJ* | P1 | 0.21 | 0.30 | 0.09 | 14.17 | 0.000 | 0.000 | TRUE |
| *ADAM8* | P1 | 0.20 | 0.52 | 0.40 | 10.68 | 0.000 | 0.000 | TRUE |
| *AQP9* | P1 | 0.20 | 0.15 | 0.00 | 15.37 | 0.000 | 0.000 | TRUE |
| *CD4* | P1 | 0.19 | 0.27 | 0.09 | 14.28 | 0.000 | 0.000 | TRUE |
| *HRH2* | P1 | 0.19 | 0.29 | 0.11 | 14.76 | 0.000 | 0.000 | TRUE |
| *RNF13* | P1 | 0.19 | 0.33 | 0.23 | 7.37 | 0.000 | 0.000 | TRUE |
| *IGF2R* | P1 | 0.18 | 0.24 | 0.06 | 12.39 | 0.000 | 0.000 | TRUE |
| *ITGB1* | P1 | 0.18 | 0.66 | 0.60 | 8.04 | 0.000 | 0.000 | TRUE |
| *CD1D* | P1 | 0.18 | 0.23 | 0.03 | 15.86 | 0.000 | 0.000 | TRUE |
| *IGFLR1* | P1 | 0.18 | 0.42 | 0.31 | 8.76 | 0.000 | 0.000 | TRUE |
| *GPR132* | P1 | 0.18 | 0.26 | 0.08 | 13.21 | 0.000 | 0.000 | TRUE |
| *MRC1* | P1 | 0.18 | 0.19 | 0.05 | 9.72 | 0.000 | 0.000 | TRUE |
| *SLC31A1* | P1 | 0.18 | 0.33 | 0.17 | 11.40 | 0.000 | 0.000 | TRUE |
| *ADORA2A* | P1 | 0.18 | 0.25 | 0.08 | 8.87 | 0.000 | 0.000 | TRUE |
| *UNC93B1* | P1 | 0.18 | 0.47 | 0.33 | 8.27 | 0.000 | 0.000 | TRUE |
| *MMP14* | P1 | 0.18 | 0.15 | 0.07 | 2.60 | 0.009 | 0.021 | TRUE |
| *GPR35* | P1 | 0.17 | 0.25 | 0.07 | 13.30 | 0.000 | 0.000 | TRUE |
| *PLXNB2* | P1 | 0.17 | 0.41 | 0.28 | 9.87 | 0.000 | 0.000 | TRUE |
| *IL10RB* | P1 | 0.17 | 0.35 | 0.21 | 9.09 | 0.000 | 0.000 | TRUE |
| *SLC30A1* | P1 | 0.17 | 0.34 | 0.19 | 8.44 | 0.000 | 0.000 | TRUE |
| *MEGF9* | P1 | 0.17 | 0.28 | 0.13 | 9.61 | 0.000 | 0.000 | TRUE |
| *P2RY13* | P1 | 0.17 | 0.19 | 0.04 | 14.79 | 0.000 | 0.000 | TRUE |
| *LRP10* | P1 | 0.16 | 0.34 | 0.26 | 4.95 | 0.000 | 0.000 | TRUE |
| *TLR8* | P1 | 0.16 | 0.17 | 0.01 | 11.74 | 0.000 | 0.000 | TRUE |
| *TMED7* | P1 | 0.16 | 0.53 | 0.44 | 6.38 | 0.000 | 0.000 | TRUE |
| *SIRPB1* | P1 | 0.16 | 0.20 | 0.03 | 15.09 | 0.000 | 0.000 | TRUE |
| *PTPRC* | P1 | 0.16 | 0.91 | 0.91 | 9.48 | 0.000 | 0.000 | TRUE |
| *TSPAN14* | P1 | 0.16 | 0.27 | 0.10 | 11.39 | 0.000 | 0.000 | TRUE |
| *LILRA2* | P1 | 0.16 | 0.23 | 0.06 | 14.65 | 0.000 | 0.000 | TRUE |
| *CSF3R* | P1 | 0.16 | 0.67 | 0.65 | 7.74 | 0.000 | 0.000 | TRUE |
| *MFSD2A* | P1 | 0.15 | 0.23 | 0.06 | 13.26 | 0.000 | 0.000 | TRUE |
| *NPTN* | P1 | 0.15 | 0.33 | 0.20 | 6.33 | 0.000 | 0.000 | TRUE |
| *LILRA5* | P1 | 0.15 | 0.15 | 0.01 | 14.33 | 0.000 | 0.000 | TRUE |
| *TMEM9B* | P1 | 0.15 | 0.41 | 0.31 | 8.03 | 0.000 | 0.000 | TRUE |
| *HAVCR2* | P1 | 0.15 | 0.31 | 0.16 | 9.70 | 0.000 | 0.000 | TRUE |
| *LILRA1* | P1 | 0.15 | 0.18 | 0.02 | 12.52 | 0.000 | 0.000 | TRUE |
| *SECTM1* | P1 | 0.14 | 0.18 | 0.04 | 11.37 | 0.000 | 0.000 | TRUE |
| *NOTCH2* | P1 | 0.14 | 0.31 | 0.17 | 9.32 | 0.000 | 0.000 | TRUE |
| *CCRL2* | P1 | 0.14 | 0.18 | 0.04 | 9.09 | 0.000 | 0.000 | TRUE |
| *PECAM1* | P1 | 0.14 | 0.47 | 0.45 | 7.27 | 0.000 | 0.000 | TRUE |
| *NFAM1* | P1 | 0.14 | 0.17 | 0.02 | 15.56 | 0.000 | 0.000 | TRUE |
| *SEMA4A* | P1 | 0.14 | 0.42 | 0.30 | 7.67 | 0.000 | 0.000 | TRUE |
| *NPC1* | P1 | 0.14 | 0.22 | 0.09 | 8.72 | 0.000 | 0.000 | TRUE |
| *THBD* | P1 | 0.14 | 0.14 | 0.03 | 10.26 | 0.000 | 0.000 | TRUE |
| *ADRB2* | P1 | 0.14 | 0.19 | 0.06 | 12.17 | 0.000 | 0.000 | TRUE |
| *SEMA6B* | P1 | 0.13 | 0.22 | 0.08 | 10.36 | 0.000 | 0.000 | TRUE |
| *CD180* | P1 | 0.13 | 0.19 | 0.07 | 10.81 | 0.000 | 0.000 | TRUE |
| *SERINC1* | P1 | 0.13 | 0.58 | 0.55 | 4.30 | 0.000 | 0.000 | TRUE |
| *IL4R* | P1 | 0.13 | 0.31 | 0.16 | 8.24 | 0.000 | 0.000 | TRUE |
| *SLC44A1* | P1 | 0.13 | 0.38 | 0.34 | 7.32 | 0.000 | 0.000 | TRUE |
| *ADAM10* | P1 | 0.13 | 0.55 | 0.51 | 4.80 | 0.000 | 0.000 | TRUE |
| *MYOF* | P1 | 0.13 | 0.19 | 0.06 | 9.72 | 0.000 | 0.000 | TRUE |
| *ENTPD1* | P1 | 0.13 | 0.18 | 0.04 | 14.71 | 0.000 | 0.000 | TRUE |
| *PRLR* | P1 | 0.12 | 0.14 | 0.02 | 11.88 | 0.000 | 0.000 | TRUE |
| *TMX4* | P1 | 0.12 | 0.54 | 0.55 | 7.37 | 0.000 | 0.000 | TRUE |
| *SORT1* | P1 | 0.12 | 0.19 | 0.06 | 10.57 | 0.000 | 0.000 | TRUE |
| *CD163* | P1 | 0.12 | 0.13 | 0.00 | 14.48 | 0.000 | 0.000 | TRUE |
| *SLC36A4* | P1 | 0.12 | 0.30 | 0.21 | 6.04 | 0.000 | 0.000 | TRUE |
| *FCER1A* | P1 | 0.12 | 0.14 | 0.05 | 8.71 | 0.000 | 0.000 | TRUE |
| *MSLN* | P1 | 0.11 | 0.18 | 0.10 | 5.44 | 0.000 | 0.000 | TRUE |
| *C3AR1* | P1 | 0.11 | 0.11 | 0.02 | 8.51 | 0.000 | 0.000 | TRUE |
| *UPK3A* | P1 | 0.11 | 0.16 | 0.04 | 10.23 | 0.000 | 0.000 | TRUE |
| *PSEN1* | P1 | 0.11 | 0.36 | 0.28 | 4.84 | 0.000 | 0.000 | TRUE |
| *CSF1* | P1 | 0.11 | 0.17 | 0.09 | 9.47 | 0.000 | 0.000 | TRUE |
| *FCGR1B* | P1 | 0.11 | 0.16 | 0.06 | 9.47 | 0.000 | 0.000 | TRUE |
| *CR1* | P1 | 0.10 | 0.11 | 0.01 | 11.44 | 0.000 | 0.000 | TRUE |
| *LYPD2* | P1 | 0.10 | 0.06 | 0.00 | 5.30 | 0.000 | 0.000 | TRUE |
| *SLC7A5* | P1 | 0.10 | 0.35 | 0.28 | 5.27 | 0.000 | 0.000 | TRUE |
| *SLC24A4* | P1 | 0.10 | 0.12 | 0.02 | 10.47 | 0.000 | 0.000 | TRUE |
| *MCOLN1* | P1 | 0.10 | 0.19 | 0.11 | 8.52 | 0.000 | 0.000 | TRUE |
| *ADCY7* | P1 | 0.10 | 0.21 | 0.10 | 9.40 | 0.000 | 0.000 | TRUE |
| *VNN2* | P1 | 0.10 | 0.10 | 0.01 | 10.67 | 0.000 | 0.000 | TRUE |
| *FCGR2B* | P1 | 0.10 | 0.11 | 0.03 | 8.25 | 0.000 | 0.000 | TRUE |
| *ANO6* | P1 | 0.09 | 0.27 | 0.21 | 5.27 | 0.000 | 0.000 | TRUE |
| *SLC8A1* | P1 | 0.09 | 0.12 | 0.02 | 11.20 | 0.000 | 0.000 | TRUE |
| *CD300LF* | P1 | 0.09 | 0.19 | 0.11 | 6.41 | 0.000 | 0.000 | TRUE |
| *TTYH3* | P1 | 0.09 | 0.23 | 0.17 | 6.51 | 0.000 | 0.000 | TRUE |
| *SIRPB2* | P1 | 0.09 | 0.13 | 0.03 | 10.11 | 0.000 | 0.000 | TRUE |
| *CEACAM4* | P1 | 0.08 | 0.17 | 0.09 | 5.33 | 0.000 | 0.000 | TRUE |
| *GPR84* | P1 | 0.08 | 0.13 | 0.03 | 9.02 | 0.000 | 0.000 | TRUE |
| *CD300C* | P1 | 0.08 | 0.12 | 0.02 | 11.52 | 0.000 | 0.000 | TRUE |
| *TLR1* | P1 | 0.08 | 0.22 | 0.15 | 2.39 | 0.017 | 0.037 | TRUE |
| *SIGLEC10* | P1 | 0.08 | 0.10 | 0.02 | 9.49 | 0.000 | 0.000 | TRUE |
| *PANX1* | P1 | 0.08 | 0.16 | 0.07 | 3.37 | 0.001 | 0.002 | TRUE |
| *P2RY6* | P1 | 0.08 | 0.11 | 0.00 | 6.69 | 0.000 | 0.000 | TRUE |

| | | | | | | | | |
|---|---|---|---|---|---|---|---|---|
| *NRCAM* | P1 | 0.08 | 0.08 | 0.01 | 2.29 | 0.022 | 0.047 | TRUE |
| *SLC8B1* | P1 | 0.08 | 0.19 | 0.12 | 5.36 | 0.000 | 0.000 | TRUE |
| *OLR1* | P1 | 0.08 | 0.10 | 0.01 | 9.15 | 0.000 | 0.000 | TRUE |
| *TMEM104* | P1 | 0.08 | 0.14 | 0.07 | 6.80 | 0.000 | 0.000 | TRUE |
| *SLC11A2* | P1 | 0.07 | 0.22 | 0.17 | 3.04 | 0.002 | 0.006 | TRUE |
| *UBAC2* | P1 | 0.07 | 0.36 | 0.35 | 3.19 | 0.001 | 0.004 | TRUE |
| *MILR1* | P1 | 0.07 | 0.17 | 0.11 | 4.08 | 0.000 | 0.000 | TRUE |
| *VSTM1* | P1 | 0.07 | 0.13 | 0.09 | 4.17 | 0.000 | 0.000 | TRUE |
| *SIGLEC14* | P1 | 0.07 | 0.09 | 0.01 | 10.74 | 0.000 | 0.000 | TRUE |
| *TNFRSF10A* | P1 | 0.07 | 0.18 | 0.13 | 4.22 | 0.000 | 0.000 | TRUE |
| *MSR1* | P1 | 0.07 | 0.07 | 0.00 | 10.14 | 0.000 | 0.000 | TRUE |
| *RECK* | P1 | 0.07 | 0.09 | 0.07 | 3.97 | 0.000 | 0.000 | TRUE |
| *OSTM1* | P1 | 0.07 | 0.15 | 0.10 | 4.41 | 0.000 | 0.000 | TRUE |
| *TPCN1* | P1 | 0.06 | 0.11 | 0.06 | 6.01 | 0.000 | 0.000 | TRUE |
| *C5AR2* | P1 | 0.06 | 0.08 | 0.01 | 10.62 | 0.000 | 0.000 | TRUE |
| *PLB1* | P1 | 0.06 | 0.16 | 0.11 | 5.72 | 0.000 | 0.000 | TRUE |
| *FPR1* | P1 | 0.06 | 0.06 | 0.00 | 8.27 | 0.000 | 0.000 | TRUE |
| *TMEM154* | P1 | 0.06 | 0.14 | 0.09 | 3.70 | 0.000 | 0.001 | TRUE |
| *CPM* | P1 | 0.06 | 0.11 | 0.03 | 7.35 | 0.000 | 0.000 | TRUE |
| *FURIN* | P1 | 0.06 | 0.32 | 0.32 | 3.52 | 0.000 | 0.001 | TRUE |
| *VNN3* | P1 | 0.06 | 0.06 | 0.00 | 10.31 | 0.000 | 0.000 | TRUE |
| *SLC9A1* | P1 | 0.06 | 0.14 | 0.09 | 5.58 | 0.000 | 0.000 | TRUE |
| *MUC12* | P1 | 0.06 | 0.10 | 0.04 | 6.53 | 0.000 | 0.000 | TRUE |
| *ATG9A* | P1 | 0.06 | 0.13 | 0.08 | 5.04 | 0.000 | 0.000 | TRUE |
| *SERINC5* | P1 | 0.06 | 0.13 | 0.08 | 3.86 | 0.000 | 0.000 | TRUE |
| *SLC36A1* | P1 | 0.06 | 0.11 | 0.05 | 6.92 | 0.000 | 0.000 | TRUE |
| *CD276* | P1 | 0.06 | 0.08 | 0.01 | 4.42 | 0.000 | 0.000 | TRUE |
| *ACVR1B* | P1 | 0.06 | 0.18 | 0.12 | 4.72 | 0.000 | 0.000 | TRUE |
| *ASGR2* | P1 | 0.06 | 0.08 | 0.02 | 8.65 | 0.000 | 0.000 | TRUE |
| *TMEM8A* | P1 | 0.06 | 0.18 | 0.14 | 3.19 | 0.001 | 0.004 | TRUE |
| *FFAR2* | P1 | 0.06 | 0.07 | 0.00 | 10.66 | 0.000 | 0.000 | TRUE |
| *ASGR1* | P1 | 0.05 | 0.12 | 0.08 | 4.74 | 0.000 | 0.000 | TRUE |
| *TNFSF8* | P1 | 0.05 | 0.08 | 0.03 | 4.94 | 0.000 | 0.000 | TRUE |
| *S1PR3* | P1 | 0.05 | 0.06 | 0.01 | 8.28 | 0.000 | 0.000 | TRUE |
| *FCGR3A* | P1 | 0.05 | 0.04 | 0.00 | 5.98 | 0.000 | 0.000 | TRUE |
| *NT5E* | P1 | 0.05 | 0.07 | 0.06 | -2.31 | 0.021 | 0.045 | TRUE |
| *CD22* | P1 | 0.05 | 0.07 | 0.02 | 7.92 | 0.000 | 0.000 | TRUE |
| *GPBAR1* | P1 | 0.05 | 0.07 | 0.02 | 6.34 | 0.000 | 0.000 | TRUE |
| *SLC12A9* | P1 | 0.05 | 0.14 | 0.09 | 4.14 | 0.000 | 0.000 | TRUE |
| *SELPLG* | P1 | 0.05 | 0.15 | 0.11 | 3.29 | 0.001 | 0.003 | TRUE |
| *OPN3* | P1 | 0.05 | 0.13 | 0.07 | 5.44 | 0.000 | 0.000 | TRUE |
| *TLR6* | P1 | 0.05 | 0.12 | 0.06 | 4.21 | 0.000 | 0.000 | TRUE |
| *TMEM63B* | P1 | 0.05 | 0.10 | 0.06 | 6.57 | 0.000 | 0.000 | TRUE |
| *SLC22A15* | P1 | 0.05 | 0.06 | 0.01 | 5.82 | 0.000 | 0.000 | TRUE |
| *P2RY2* | P1 | 0.05 | 0.07 | 0.02 | 8.26 | 0.000 | 0.000 | TRUE |
| *GPR160* | P1 | 0.05 | 0.10 | 0.06 | 3.45 | 0.001 | 0.002 | TRUE |
| *TMEM158* | P1 | 0.05 | 0.06 | 0.03 | 2.68 | 0.007 | 0.017 | TRUE |
| *SIGLEC9* | P1 | 0.04 | 0.06 | 0.00 | 8.51 | 0.000 | 0.000 | TRUE |
| *IL1R1* | P1 | 0.04 | 0.05 | 0.01 | 8.01 | 0.000 | 0.000 | TRUE |
| *STS* | P1 | 0.04 | 0.07 | 0.02 | 5.53 | 0.000 | 0.000 | TRUE |
| *HEG1* | P1 | 0.04 | 0.08 | 0.02 | 3.17 | 0.002 | 0.004 | TRUE |
| *GPRC5A* | P1 | 0.04 | 0.05 | 0.01 | 7.91 | 0.000 | 0.000 | TRUE |
| *SDC2* | P1 | 0.04 | 0.06 | 0.00 | 7.94 | 0.000 | 0.000 | TRUE |
| *SLC2A9* | P1 | 0.04 | 0.07 | 0.03 | 6.14 | 0.000 | 0.000 | TRUE |
| *SLCO4A1* | P1 | 0.04 | 0.08 | 0.06 | -4.21 | 0.000 | 0.000 | TRUE |
| *HRH1* | P1 | 0.04 | 0.05 | 0.01 | 8.48 | 0.000 | 0.000 | TRUE |
| *TLR5* | P1 | 0.04 | 0.05 | 0.00 | 8.48 | 0.000 | 0.000 | TRUE |
| *CDH23* | P1 | 0.04 | 0.05 | 0.01 | 6.40 | 0.000 | 0.000 | TRUE |
| *FOLR2* | P1 | 0.03 | 0.05 | 0.01 | 6.09 | 0.000 | 0.000 | TRUE |
| *GPR65* | P1 | 0.03 | 0.06 | 0.03 | 5.21 | 0.000 | 0.000 | TRUE |
| *IL1R2* | P1 | 0.03 | 0.04 | 0.00 | 6.34 | 0.000 | 0.000 | TRUE |
| *CCR5* | P1 | 0.03 | 0.05 | 0.00 | 5.15 | 0.000 | 0.000 | TRUE |
| *LDLRAD3* | P1 | 0.03 | 0.05 | 0.01 | 3.26 | 0.001 | 0.003 | TRUE |
| *TMEM8B* | P1 | 0.03 | 0.06 | 0.03 | 4.75 | 0.000 | 0.000 | TRUE |
| *ABCC3* | P1 | 0.03 | 0.04 | 0.00 | 6.81 | 0.000 | 0.000 | TRUE |
| *SLCO4C1* | P1 | 0.03 | 0.04 | 0.02 | 5.16 | 0.000 | 0.000 | TRUE |
| *TLR7* | P1 | 0.03 | 0.04 | 0.01 | 6.28 | 0.000 | 0.000 | TRUE |
| *SIGLEC7* | P1 | 0.03 | 0.04 | 0.00 | 6.33 | 0.000 | 0.000 | TRUE |
| *SLC16A6* | P1 | 0.03 | 0.03 | 0.00 | 6.35 | 0.000 | 0.000 | TRUE |
| *CD101* | P1 | 0.03 | 0.04 | 0.01 | 6.29 | 0.000 | 0.000 | TRUE |
| *HTR7* | P1 | 0.03 | 0.05 | 0.02 | 3.61 | 0.000 | 0.001 | TRUE |
| *EPHB2* | P1 | 0.03 | 0.04 | 0.01 | 3.31 | 0.001 | 0.002 | TRUE |
| *CRTAM* | P1 | 0.03 | 0.05 | 0.02 | 5.71 | 0.000 | 0.000 | TRUE |
| *SLC37A2* | P1 | 0.03 | 0.07 | 0.04 | 2.58 | 0.010 | 0.022 | TRUE |
| *ANKH* | P1 | 0.02 | 0.06 | 0.04 | 2.82 | 0.005 | 0.012 | TRUE |
| *GDPD5* | P1 | 0.02 | 0.05 | 0.02 | 4.06 | 0.000 | 0.000 | TRUE |
| *SERINC2* | P1 | 0.02 | 0.04 | 0.02 | 4.44 | 0.000 | 0.000 | TRUE |
| *CLEC12B* | P1 | 0.02 | 0.04 | 0.01 | 3.55 | 0.000 | 0.001 | TRUE |
| *SIGLEC1* | P1 | 0.02 | 0.04 | 0.01 | 6.22 | 0.000 | 0.000 | TRUE |
| *FPR2* | P1 | 0.02 | 0.02 | 0.00 | 4.60 | 0.000 | 0.000 | TRUE |
| *DCSTAMP* | P1 | 0.02 | 0.02 | 0.00 | 2.61 | 0.009 | 0.021 | TRUE |
| *CCR2* | P1 | 0.02 | 0.03 | 0.00 | 6.54 | 0.000 | 0.000 | TRUE |
| *IL31RA* | P1 | 0.02 | 0.03 | 0.00 | 4.83 | 0.000 | 0.000 | TRUE |
| *SIGLEC5* | P1 | 0.02 | 0.04 | 0.01 | 4.26 | 0.000 | 0.000 | TRUE |
| *CD1B* | P1 | 0.02 | 0.03 | 0.00 | 3.22 | 0.001 | 0.003 | TRUE |
| *GJB2* | P1 | 0.02 | 0.03 | 0.01 | 2.98 | 0.003 | 0.007 | TRUE |
| *FPR3* | P1 | 0.02 | 0.02 | 0.00 | 3.76 | 0.000 | 0.000 | TRUE |
| *CMKLR1* | P1 | 0.02 | 0.02 | 0.00 | 3.31 | 0.001 | 0.002 | TRUE |
| *GPR141* | P1 | 0.02 | 0.05 | 0.03 | 3.36 | 0.001 | 0.002 | TRUE |
| *TREM2* | P1 | 0.02 | 0.03 | 0.01 | 2.36 | 0.018 | 0.040 | TRUE |
| *ITGB3* | P1 | 0.02 | 0.02 | 0.00 | 4.13 | 0.000 | 0.000 | TRUE |
| *NTSR1* | P1 | 0.02 | 0.03 | 0.01 | 3.74 | 0.000 | 0.001 | TRUE |
| *DAGLA* | P1 | 0.02 | 0.02 | 0.00 | 3.38 | 0.001 | 0.002 | TRUE |
| *TMEM150B* | P1 | 0.02 | 0.04 | 0.02 | 3.65 | 0.000 | 0.001 | TRUE |
| *TGFA* | P1 | 0.01 | 0.03 | 0.00 | 3.78 | 0.000 | 0.000 | TRUE |
| *OTOA* | P1 | 0.01 | 0.03 | 0.01 | 3.49 | 0.000 | 0.001 | TRUE |
| *IL1RL2* | P1 | 0.01 | 0.02 | 0.00 | 5.27 | 0.000 | 0.000 | TRUE |
| *CX3CR1* | P1 | 0.01 | 0.02 | 0.00 | 2.82 | 0.005 | 0.012 | TRUE |
| *LILRA6* | P1 | 0.01 | 0.02 | 0.00 | 5.40 | 0.000 | 0.000 | TRUE |
| *GPR18* | P1 | 0.01 | 0.03 | 0.01 | 2.33 | 0.020 | 0.043 | TRUE |
| *MMP25* | P1 | 0.01 | 0.03 | 0.02 | 3.37 | 0.001 | 0.002 | TRUE |

| | | | | | | | | |
|---|---|---|---|---|---|---|---|---|
| *SLC4A8* | P1 | 0.01 | 0.04 | 0.03 | 2.96 | 0.003 | 0.008 | TRUE |
| *LYPD3* | P1 | 0.01 | 0.02 | 0.01 | 2.36 | 0.018 | 0.039 | TRUE |
| *LILRB5* | P1 | 0.01 | 0.02 | 0.00 | 2.74 | 0.006 | 0.015 | TRUE |
| *CYSLTR2* | P1 | 0.01 | 0.06 | 0.06 | 2.94 | 0.003 | 0.008 | TRUE |
| *SLC6A8* | P1 | 0.01 | 0.03 | 0.01 | 3.37 | 0.001 | 0.002 | TRUE |
| *GABRR2* | P1 | 0.01 | 0.03 | 0.00 | 2.63 | 0.008 | 0.019 | TRUE |
| *CEACAM3* | P1 | 0.01 | 0.01 | 0.00 | 3.13 | 0.002 | 0.004 | TRUE |
| *MMP17* | P1 | 0.01 | 0.02 | 0.01 | 2.86 | 0.004 | 0.010 | TRUE |
| *NCR1* | P1 | 0.01 | 0.03 | 0.01 | 2.44 | 0.015 | 0.032 | TRUE |
| *VIPR1* | P1 | 0.01 | 0.02 | 0.00 | 3.79 | 0.000 | 0.000 | TRUE |
| *LRP3* | P1 | 0.01 | 0.02 | 0.01 | 2.29 | 0.022 | 0.047 | TRUE |
| *GRIN2C* | P1 | 0.01 | 0.03 | 0.02 | 2.53 | 0.012 | 0.026 | TRUE |
| *NIPAL4* | P1 | 0.01 | 0.02 | 0.01 | 3.53 | 0.000 | 0.001 | TRUE |
| *SCN4B* | P1 | 0.01 | 0.01 | 0.00 | 3.28 | 0.001 | 0.003 | TRUE |
| *PCSK5* | P1 | 0.01 | 0.02 | 0.00 | 2.40 | 0.016 | 0.036 | TRUE |
| *NCR3LG1* | P1 | 0.01 | 0.03 | 0.02 | 2.38 | 0.017 | 0.037 | TRUE |
| *SORCS2* | P1 | 0.01 | 0.01 | 0.00 | 2.64 | 0.008 | 0.019 | TRUE |
| *LRRC32* | P1 | 0.01 | 0.01 | 0.00 | 3.07 | 0.002 | 0.005 | TRUE |
| *LY6K* | P1 | 0.01 | 0.01 | 0.00 | 3.34 | 0.001 | 0.002 | TRUE |
| *ADAM12* | P1 | 0.01 | 0.01 | 0.00 | 2.78 | 0.005 | 0.013 | TRUE |
| *ABCA6* | P1 | 0.01 | 0.01 | 0.00 | 2.32 | 0.020 | 0.044 | TRUE |
| *PKD2L1* | P1 | 0.00 | 0.01 | 0.00 | 2.77 | 0.006 | 0.013 | TRUE |
| *SLAMF9* | P1 | 0.00 | 0.01 | 0.00 | 3.07 | 0.002 | 0.005 | TRUE |
| *SLC46A2* | P1 | 0.00 | 0.00 | 0.00 | 2.68 | 0.007 | 0.017 | TRUE |
| *OR10G3* | P1 | 0.00 | 0.01 | 0.00 | 2.61 | 0.009 | 0.021 | TRUE |
| *NEO1* | P1 | 0.00 | 0.00 | 0.00 | 2.50 | 0.012 | 0.028 | TRUE |
| *SLC1A5* | P2 | 0.33 | 0.71 | 0.29 | 30.49 | 0.000 | 0.000 | TRUE |
| *EBP* | P2 | 0.32 | 0.84 | 0.46 | 24.13 | 0.000 | 0.000 | TRUE |
| *BSG* | P2 | 0.29 | 0.96 | 0.71 | 19.24 | 0.000 | 0.000 | TRUE |
| *ATP1B3* | P2 | 0.29 | 0.85 | 0.58 | 19.00 | 0.000 | 0.000 | TRUE |
| *SLC43A3* | P2 | 0.27 | 0.60 | 0.23 | 28.75 | 0.000 | 0.000 | TRUE |
| *RPN1* | P2 | 0.27 | 0.85 | 0.51 | 21.37 | 0.000 | 0.000 | TRUE |
| *SLC3A2* | P2 | 0.25 | 0.91 | 0.67 | 17.82 | 0.000 | 0.000 | TRUE |
| *NUP210* | P2 | 0.24 | 0.58 | 0.16 | 29.47 | 0.000 | 0.000 | TRUE |
| *SSR1* | P2 | 0.24 | 0.85 | 0.55 | 17.35 | 0.000 | 0.000 | TRUE |
| *PRSS21* | P2 | 0.23 | 0.73 | 0.38 | 15.41 | 0.000 | 0.000 | TRUE |
| *LMAN2* | P2 | 0.22 | 0.93 | 0.69 | 16.00 | 0.000 | 0.000 | TRUE |
| *SLC29A1* | P2 | 0.21 | 0.44 | 0.06 | 34.55 | 0.000 | 0.000 | TRUE |
| *CD320* | P2 | 0.20 | 0.49 | 0.13 | 26.95 | 0.000 | 0.000 | TRUE |
| *SLC39A4* | P2 | 0.20 | 0.61 | 0.30 | 17.69 | 0.000 | 0.000 | TRUE |
| *EMB* | P2 | 0.19 | 0.74 | 0.51 | 14.00 | 0.000 | 0.000 | TRUE |
| *TSPAN3* | P2 | 0.18 | 0.67 | 0.33 | 18.80 | 0.000 | 0.000 | TRUE |
| *TP53I13* | P2 | 0.16 | 0.69 | 0.35 | 16.08 | 0.000 | 0.000 | TRUE |
| *CD38* | P2 | 0.15 | 0.67 | 0.41 | 13.50 | 0.000 | 0.000 | TRUE |
| *CD151* | P2 | 0.15 | 0.60 | 0.32 | 15.29 | 0.000 | 0.000 | TRUE |
| *SLC38A5* | P2 | 0.14 | 0.43 | 0.14 | 22.40 | 0.000 | 0.000 | TRUE |
| *RNF167* | P2 | 0.13 | 0.67 | 0.35 | 15.84 | 0.000 | 0.000 | TRUE |
| *SLC7A1* | P2 | 0.13 | 0.41 | 0.13 | 21.81 | 0.000 | 0.000 | TRUE |
| *DPEP3* | P2 | 0.12 | 0.64 | 0.50 | 5.03 | 0.000 | 0.000 | TRUE |
| *ITGAE* | P2 | 0.12 | 0.83 | 0.59 | 10.27 | 0.000 | 0.000 | TRUE |
| *CHPT1* | P2 | 0.12 | 0.56 | 0.27 | 15.73 | 0.000 | 0.000 | TRUE |
| *SLC2A6* | P2 | 0.11 | 0.37 | 0.24 | 6.69 | 0.000 | 0.000 | TRUE |
| *TSPAN4* | P2 | 0.11 | 0.45 | 0.19 | 17.31 | 0.000 | 0.000 | TRUE |
| *SLC19A1* | P2 | 0.11 | 0.31 | 0.07 | 22.41 | 0.000 | 0.000 | TRUE |
| *SLC16A1* | P2 | 0.11 | 0.30 | 0.07 | 22.73 | 0.000 | 0.000 | TRUE |
| *SLC52A2* | P2 | 0.11 | 0.57 | 0.32 | 12.34 | 0.000 | 0.000 | TRUE |
| *SLC1A4* | P2 | 0.10 | 0.34 | 0.10 | 18.19 | 0.000 | 0.000 | TRUE |
| *PIEZO1* | P2 | 0.10 | 0.62 | 0.36 | 13.51 | 0.000 | 0.000 | TRUE |
| *SCARB1* | P2 | 0.10 | 0.35 | 0.12 | 18.06 | 0.000 | 0.000 | TRUE |
| *IL27RA* | P2 | 0.10 | 0.46 | 0.24 | 12.91 | 0.000 | 0.000 | TRUE |
| *NETO2* | P2 | 0.10 | 0.26 | 0.07 | 18.57 | 0.000 | 0.000 | TRUE |
| *CSF2RB* | P2 | 0.10 | 0.32 | 0.19 | 7.17 | 0.000 | 0.000 | TRUE |
| *SLC5A6* | P2 | 0.10 | 0.30 | 0.09 | 18.27 | 0.000 | 0.000 | TRUE |
| *LRP8* | P2 | 0.09 | 0.35 | 0.13 | 18.27 | 0.000 | 0.000 | TRUE |
| *ADAM15* | P2 | 0.09 | 0.39 | 0.18 | 15.20 | 0.000 | 0.000 | TRUE |
| *ADCY3* | P2 | 0.09 | 0.40 | 0.17 | 16.67 | 0.000 | 0.000 | TRUE |
| *TMEM245* | P2 | 0.09 | 0.39 | 0.17 | 14.41 | 0.000 | 0.000 | TRUE |
| *INSR* | P2 | 0.08 | 0.34 | 0.11 | 16.61 | 0.000 | 0.000 | TRUE |
| *SLC37A4* | P2 | 0.08 | 0.26 | 0.06 | 19.96 | 0.000 | 0.000 | TRUE |
| *SLC39A6* | P2 | 0.08 | 0.38 | 0.17 | 14.40 | 0.000 | 0.000 | TRUE |
| *SPPL2B* | P2 | 0.08 | 0.47 | 0.25 | 12.39 | 0.000 | 0.000 | TRUE |
| *TMEM161A* | P2 | 0.07 | 0.33 | 0.12 | 15.40 | 0.000 | 0.000 | TRUE |
| *SLC43A1* | P2 | 0.07 | 0.28 | 0.09 | 17.73 | 0.000 | 0.000 | TRUE |
| *TRGC2* | P2 | 0.07 | 0.33 | 0.16 | 11.31 | 0.000 | 0.000 | TRUE |
| *SLC39A14* | P2 | 0.07 | 0.20 | 0.04 | 17.33 | 0.000 | 0.000 | TRUE |
| *PKD1* | P2 | 0.07 | 0.37 | 0.16 | 15.22 | 0.000 | 0.000 | TRUE |
| *SLC22A16* | P2 | 0.07 | 0.31 | 0.13 | 13.15 | 0.000 | 0.000 | TRUE |
| *MICB* | P2 | 0.07 | 0.26 | 0.08 | 16.59 | 0.000 | 0.000 | TRUE |
| *HYAL2* | P2 | 0.07 | 0.28 | 0.12 | 11.11 | 0.000 | 0.000 | TRUE |
| *EPCAM* | P2 | 0.06 | 0.16 | 0.05 | 11.33 | 0.000 | 0.000 | TRUE |
| *CD59* | P2 | 0.06 | 0.50 | 0.34 | 7.42 | 0.000 | 0.000 | TRUE |
| *SCAP* | P2 | 0.06 | 0.34 | 0.14 | 14.03 | 0.000 | 0.000 | TRUE |
| *ATP13A1* | P2 | 0.06 | 0.34 | 0.15 | 13.47 | 0.000 | 0.000 | TRUE |
| *GPR180* | P2 | 0.06 | 0.26 | 0.09 | 13.97 | 0.000 | 0.000 | TRUE |
| *CNNM3* | P2 | 0.06 | 0.26 | 0.09 | 15.15 | 0.000 | 0.000 | TRUE |
| *LAMP5* | P2 | 0.06 | 0.09 | 0.02 | 5.51 | 0.000 | 0.000 | TRUE |
| *TMX3* | P2 | 0.06 | 0.46 | 0.25 | 11.20 | 0.000 | 0.000 | TRUE |
| *PTK7* | P2 | 0.06 | 0.27 | 0.10 | 16.44 | 0.000 | 0.000 | TRUE |
| *CLSTN1* | P2 | 0.06 | 0.49 | 0.28 | 9.51 | 0.000 | 0.000 | TRUE |
| *LTB4R* | P2 | 0.06 | 0.28 | 0.10 | 15.88 | 0.000 | 0.000 | TRUE |
| *SLC4A7* | P2 | 0.06 | 0.31 | 0.13 | 10.80 | 0.000 | 0.000 | TRUE |
| *C11orf24* | P2 | 0.06 | 0.22 | 0.07 | 13.69 | 0.000 | 0.000 | TRUE |
| *TM9SF1* | P2 | 0.06 | 0.38 | 0.20 | 11.39 | 0.000 | 0.000 | TRUE |
| *GINM1* | P2 | 0.06 | 0.64 | 0.41 | 7.68 | 0.000 | 0.000 | TRUE |
| *SLC7A6* | P2 | 0.05 | 0.34 | 0.17 | 9.37 | 0.000 | 0.000 | TRUE |
| *ADIPOR2* | P2 | 0.05 | 0.49 | 0.28 | 10.76 | 0.000 | 0.000 | TRUE |
| *SLC12A2* | P2 | 0.05 | 0.29 | 0.14 | 10.11 | 0.000 | 0.000 | TRUE |
| *FLVCR1* | P2 | 0.05 | 0.26 | 0.12 | 11.70 | 0.000 | 0.000 | TRUE |
| *VSIG10* | P2 | 0.05 | 0.24 | 0.09 | 14.16 | 0.000 | 0.000 | TRUE |
| *KCNK5* | P2 | 0.05 | 0.21 | 0.09 | 11.71 | 0.000 | 0.000 | TRUE |
| *SLC16A7* | P2 | 0.04 | 0.35 | 0.18 | 13.08 | 0.000 | 0.000 | TRUE |

| | | | | | | | | |
|---|---|---|---|---|---|---|---|---|
| *TSPAN13* | P2 | 0.04 | 0.36 | 0.22 | 7.68 | 0.000 | 0.000 | TRUE |
| *LRP5* | P2 | 0.04 | 0.22 | 0.08 | 11.68 | 0.000 | 0.000 | TRUE |
| *FAM189B* | P2 | 0.04 | 0.21 | 0.08 | 11.74 | 0.000 | 0.000 | TRUE |
| *NRXN2* | P2 | 0.04 | 0.20 | 0.09 | 11.40 | 0.000 | 0.000 | TRUE |
| *TCTN3* | P2 | 0.04 | 0.38 | 0.21 | 9.70 | 0.000 | 0.000 | TRUE |
| *GPR27* | P2 | 0.04 | 0.31 | 0.17 | 9.29 | 0.000 | 0.000 | TRUE |
| *TMEM9* | P2 | 0.04 | 0.28 | 0.14 | 12.27 | 0.000 | 0.000 | TRUE |
| *LMBR1* | P2 | 0.04 | 0.26 | 0.12 | 12.08 | 0.000 | 0.000 | TRUE |
| *SLC20A2* | P2 | 0.04 | 0.32 | 0.15 | 9.74 | 0.000 | 0.000 | TRUE |
| *TMEM67* | P2 | 0.04 | 0.21 | 0.09 | 10.69 | 0.000 | 0.000 | TRUE |
| *TSPAN5* | P2 | 0.04 | 0.17 | 0.06 | 9.17 | 0.000 | 0.000 | TRUE |
| *FGFRL1* | P2 | 0.04 | 0.19 | 0.08 | 10.32 | 0.000 | 0.000 | TRUE |
| *SLC41A3* | P2 | 0.04 | 0.30 | 0.16 | 11.35 | 0.000 | 0.000 | TRUE |
| *SIGLEC12* | P2 | 0.04 | 0.10 | 0.02 | 11.53 | 0.000 | 0.000 | TRUE |
| *ZDHHC5* | P2 | 0.04 | 0.42 | 0.24 | 9.52 | 0.000 | 0.000 | TRUE |
| *ABCA2* | P2 | 0.04 | 0.21 | 0.09 | 12.34 | 0.000 | 0.000 | TRUE |
| *TPRA1* | P2 | 0.03 | 0.24 | 0.12 | 9.82 | 0.000 | 0.000 | TRUE |
| *DAG1* | P2 | 0.03 | 0.14 | 0.05 | 9.82 | 0.000 | 0.000 | TRUE |
| *BACE1* | P2 | 0.03 | 0.18 | 0.08 | 10.65 | 0.000 | 0.000 | TRUE |
| *QSOX2* | P2 | 0.03 | 0.21 | 0.10 | 10.76 | 0.000 | 0.000 | TRUE |
| *CHRNA5* | P2 | 0.03 | 0.09 | 0.01 | 11.71 | 0.000 | 0.000 | TRUE |
| *PILRB* | P2 | 0.03 | 0.36 | 0.21 | 7.62 | 0.000 | 0.000 | TRUE |
| *MFSD5* | P2 | 0.03 | 0.21 | 0.10 | 7.98 | 0.000 | 0.000 | TRUE |
| *DAGLB* | P2 | 0.03 | 0.28 | 0.14 | 10.02 | 0.000 | 0.000 | TRUE |
| *MRC2* | P2 | 0.03 | 0.21 | 0.11 | 10.15 | 0.000 | 0.000 | TRUE |
| *C1orf159* | P2 | 0.03 | 0.14 | 0.05 | 10.41 | 0.000 | 0.000 | TRUE |
| *SUSD3* | P2 | 0.03 | 0.17 | 0.09 | 8.89 | 0.000 | 0.000 | TRUE |
| *GPC6* | P2 | 0.03 | 0.12 | 0.04 | 4.64 | 0.000 | 0.000 | TRUE |
| *SLC12A5* | P2 | 0.03 | 0.15 | 0.06 | 9.38 | 0.000 | 0.000 | TRUE |
| *RTN4R* | P2 | 0.03 | 0.10 | 0.03 | 11.74 | 0.000 | 0.000 | TRUE |
| *SLC2A8* | P2 | 0.03 | 0.10 | 0.03 | 9.32 | 0.000 | 0.000 | TRUE |
| *FAM171A2* | P2 | 0.02 | 0.11 | 0.03 | 10.42 | 0.000 | 0.000 | TRUE |
| *CDH4* | P2 | 0.02 | 0.15 | 0.06 | 4.87 | 0.000 | 0.000 | TRUE |
| *ESAM* | P2 | 0.02 | 0.14 | 0.07 | 6.47 | 0.000 | 0.000 | TRUE |
| *KLRG1* | P2 | 0.02 | 0.14 | 0.06 | 9.51 | 0.000 | 0.000 | TRUE |
| *P2RY1* | P2 | 0.02 | 0.14 | 0.05 | 9.53 | 0.000 | 0.000 | TRUE |
| *SLC2A5* | P2 | 0.02 | 0.13 | 0.06 | 7.29 | 0.000 | 0.000 | TRUE |
| *IGSF8* | P2 | 0.02 | 0.24 | 0.13 | 8.25 | 0.000 | 0.000 | TRUE |
| *ADORA2B* | P2 | 0.02 | 0.09 | 0.03 | 8.39 | 0.000 | 0.000 | TRUE |
| *CEACAM6* | P2 | 0.02 | 0.08 | 0.03 | 3.52 | 0.000 | 0.001 | TRUE |
| *TSPAN7* | P2 | 0.02 | 0.13 | 0.06 | 6.08 | 0.000 | 0.000 | TRUE |
| *TMEM62* | P2 | 0.02 | 0.16 | 0.07 | 7.36 | 0.000 | 0.000 | TRUE |
| *TPSG1* | P2 | 0.02 | 0.04 | 0.01 | 2.47 | 0.013 | 0.030 | TRUE |
| *GPR137* | P2 | 0.02 | 0.28 | 0.16 | 9.68 | 0.000 | 0.000 | TRUE |
| *TSPAN11* | P2 | 0.02 | 0.04 | 0.00 | 4.66 | 0.000 | 0.000 | TRUE |
| *LMAN2L* | P2 | 0.02 | 0.12 | 0.05 | 9.37 | 0.000 | 0.000 | TRUE |
| *IGF1R* | P2 | 0.02 | 0.14 | 0.07 | 8.71 | 0.000 | 0.000 | TRUE |
| *PODXL2* | P2 | 0.02 | 0.10 | 0.04 | 9.14 | 0.000 | 0.000 | TRUE |
| *CELSR3* | P2 | 0.02 | 0.06 | 0.01 | 8.87 | 0.000 | 0.000 | TRUE |
| *LRP11* | P2 | 0.02 | 0.05 | 0.01 | 5.89 | 0.000 | 0.000 | TRUE |
| *LRRN1* | P2 | 0.02 | 0.09 | 0.03 | 4.60 | 0.000 | 0.000 | TRUE |
| *P2RY11* | P2 | 0.02 | 0.15 | 0.08 | 8.24 | 0.000 | 0.000 | TRUE |
| *NCR3* | P2 | 0.02 | 0.07 | 0.03 | 5.88 | 0.000 | 0.000 | TRUE |
| *DGCR2* | P2 | 0.02 | 0.14 | 0.06 | 8.93 | 0.000 | 0.000 | TRUE |
| *LRP4* | P2 | 0.02 | 0.05 | 0.01 | 5.51 | 0.000 | 0.000 | TRUE |
| *SLC26A6* | P2 | 0.02 | 0.11 | 0.05 | 7.64 | 0.000 | 0.000 | TRUE |
| *PIGO* | P2 | 0.02 | 0.14 | 0.07 | 8.06 | 0.000 | 0.000 | TRUE |
| *TMEM116* | P2 | 0.02 | 0.10 | 0.04 | 8.79 | 0.000 | 0.000 | TRUE |
| *JAM3* | P2 | 0.02 | 0.08 | 0.02 | 6.55 | 0.000 | 0.000 | TRUE |
| *ADAM22* | P2 | 0.02 | 0.07 | 0.02 | 6.12 | 0.000 | 0.000 | TRUE |
| *DCBLD2* | P2 | 0.02 | 0.07 | 0.02 | 6.75 | 0.000 | 0.000 | TRUE |
| *SLC29A2* | P2 | 0.02 | 0.06 | 0.02 | 8.03 | 0.000 | 0.000 | TRUE |
| *TYRO3* | P2 | 0.01 | 0.05 | 0.01 | 8.43 | 0.000 | 0.000 | TRUE |
| *CDH24* | P2 | 0.01 | 0.05 | 0.01 | 8.19 | 0.000 | 0.000 | TRUE |
| *SLC2A13* | P2 | 0.01 | 0.06 | 0.02 | 6.65 | 0.000 | 0.000 | TRUE |
| *SLC38A9* | P2 | 0.01 | 0.17 | 0.09 | 7.70 | 0.000 | 0.000 | TRUE |
| *ACVR2B* | P2 | 0.01 | 0.05 | 0.01 | 6.85 | 0.000 | 0.000 | TRUE |
| *ENPP6* | P2 | 0.01 | 0.02 | 0.00 | 3.23 | 0.001 | 0.003 | TRUE |
| *ABCG2* | P2 | 0.01 | 0.04 | 0.01 | 5.56 | 0.000 | 0.000 | TRUE |
| *CD19* | P2 | 0.01 | 0.05 | 0.02 | 6.00 | 0.000 | 0.000 | TRUE |
| *FGFR1* | P2 | 0.01 | 0.12 | 0.07 | 5.96 | 0.000 | 0.000 | TRUE |
| *GPC1* | P2 | 0.01 | 0.04 | 0.01 | 7.09 | 0.000 | 0.000 | TRUE |
| *VNN1* | P2 | 0.01 | 0.09 | 0.05 | 3.82 | 0.000 | 0.000 | TRUE |
| *ITGA7* | P2 | 0.01 | 0.03 | 0.01 | 4.79 | 0.000 | 0.000 | TRUE |
| *RAMP2* | P2 | 0.01 | 0.04 | 0.01 | 5.73 | 0.000 | 0.000 | TRUE |
| *RNF150* | P2 | 0.01 | 0.04 | 0.01 | 3.02 | 0.003 | 0.006 | TRUE |
| *TNFRSF8* | P2 | 0.01 | 0.05 | 0.01 | 5.02 | 0.000 | 0.000 | TRUE |
| *TCTN2* | P2 | 0.01 | 0.06 | 0.02 | 6.95 | 0.000 | 0.000 | TRUE |
| *GPR3* | P2 | 0.01 | 0.04 | 0.01 | 7.17 | 0.000 | 0.000 | TRUE |
| *CD163L1* | P2 | 0.01 | 0.03 | 0.01 | 3.05 | 0.002 | 0.006 | TRUE |
| *EPHB4* | P2 | 0.01 | 0.05 | 0.02 | 5.60 | 0.000 | 0.000 | TRUE |
| *NRG2* | P2 | 0.01 | 0.04 | 0.01 | 5.29 | 0.000 | 0.000 | TRUE |
| *ABCB9* | P2 | 0.01 | 0.06 | 0.02 | 6.70 | 0.000 | 0.000 | TRUE |
| *CLDN7* | P2 | 0.01 | 0.09 | 0.04 | 4.61 | 0.000 | 0.000 | TRUE |
| *GJA3* | P2 | 0.01 | 0.04 | 0.01 | 6.57 | 0.000 | 0.000 | TRUE |
| *ATP1A3* | P2 | 0.01 | 0.04 | 0.01 | 6.65 | 0.000 | 0.000 | TRUE |
| *SMO* | P2 | 0.01 | 0.03 | 0.01 | 6.19 | 0.000 | 0.000 | TRUE |
| *SLC46A1* | P2 | 0.01 | 0.03 | 0.01 | 7.09 | 0.000 | 0.000 | TRUE |
| *LHFPL5* | P2 | 0.01 | 0.06 | 0.03 | 4.45 | 0.000 | 0.000 | TRUE |
| *SLC22A5* | P2 | 0.01 | 0.04 | 0.02 | 6.03 | 0.000 | 0.000 | TRUE |
| *SLC2A10* | P2 | 0.01 | 0.03 | 0.01 | 3.52 | 0.000 | 0.001 | TRUE |
| *SLC16A4* | P2 | 0.01 | 0.07 | 0.03 | 5.14 | 0.000 | 0.000 | TRUE |
| *MCAM* | P2 | 0.01 | 0.03 | 0.01 | 3.57 | 0.000 | 0.001 | TRUE |
| *ASIC1* | P2 | 0.01 | 0.02 | 0.00 | 5.97 | 0.000 | 0.000 | TRUE |
| *NLGN2* | P2 | 0.01 | 0.04 | 0.01 | 6.06 | 0.000 | 0.000 | TRUE |
| *EVA1C* | P2 | 0.01 | 0.04 | 0.01 | 3.75 | 0.000 | 0.001 | TRUE |
| *LYPD5* | P2 | 0.01 | 0.02 | 0.00 | 4.80 | 0.000 | 0.000 | TRUE |
| *DLL3* | P2 | 0.01 | 0.02 | 0.00 | 6.03 | 0.000 | 0.000 | TRUE |
| *SLC15A2* | P2 | 0.01 | 0.07 | 0.04 | 6.20 | 0.000 | 0.000 | TRUE |
| *SCNN1D* | P2 | 0.01 | 0.03 | 0.01 | 4.07 | 0.000 | 0.000 | TRUE |
| *SLC19A2* | P2 | 0.01 | 0.08 | 0.04 | 5.56 | 0.000 | 0.000 | TRUE |

| | | | | | | | | |
|---|---|---|---|---|---|---|---|---|
| *GPR137C* | P2 | 0.01 | 0.03 | 0.01 | 7.27 | 0.000 | 0.000 | TRUE |
| *MAMDC4* | P2 | 0.01 | 0.04 | 0.01 | 4.78 | 0.000 | 0.000 | TRUE |
| *OXER1* | P2 | 0.01 | 0.04 | 0.01 | 4.84 | 0.000 | 0.000 | TRUE |
| *KISS1R* | P2 | 0.01 | 0.02 | 0.00 | 3.81 | 0.000 | 0.000 | TRUE |
| *PIEZO2* | P2 | 0.01 | 0.02 | 0.00 | 5.66 | 0.000 | 0.000 | TRUE |
| *LPAR4* | P2 | 0.01 | 0.06 | 0.03 | 3.50 | 0.000 | 0.001 | TRUE |
| *FOLH1* | P2 | 0.01 | 0.04 | 0.01 | 2.76 | 0.006 | 0.014 | TRUE |
| *EFNA2* | P2 | 0.01 | 0.04 | 0.02 | 2.48 | 0.013 | 0.029 | TRUE |
| *KDR* | P2 | 0.01 | 0.01 | 0.00 | 2.47 | 0.014 | 0.030 | TRUE |
| *SLC29A4* | P2 | 0.01 | 0.02 | 0.00 | 4.58 | 0.000 | 0.000 | TRUE |
| *NPHS1* | P2 | 0.01 | 0.02 | 0.00 | 3.98 | 0.000 | 0.000 | TRUE |
| *SUSD5* | P2 | 0.00 | 0.02 | 0.00 | 2.88 | 0.004 | 0.010 | TRUE |
| *EFNB2* | P2 | 0.00 | 0.05 | 0.02 | 3.64 | 0.000 | 0.001 | TRUE |
| *EPOR* | P2 | 0.00 | 0.03 | 0.01 | 5.18 | 0.000 | 0.000 | TRUE |
| *SLC2A4* | P2 | 0.00 | 0.01 | 0.00 | 3.77 | 0.000 | 0.000 | TRUE |
| *CSPG5* | P2 | 0.00 | 0.02 | 0.00 | 4.91 | 0.000 | 0.000 | TRUE |
| *CUZD1* | P2 | 0.00 | 0.02 | 0.01 | 2.52 | 0.012 | 0.026 | TRUE |
| *GPR150* | P2 | 0.00 | 0.01 | 0.00 | 2.87 | 0.004 | 0.010 | TRUE |
| *FZD7* | P2 | 0.00 | 0.04 | 0.02 | 3.99 | 0.000 | 0.000 | TRUE |
| *TSPAN6* | P2 | 0.00 | 0.01 | 0.00 | 3.02 | 0.003 | 0.006 | TRUE |
| *CHRNA10* | P2 | 0.00 | 0.04 | 0.02 | 2.62 | 0.009 | 0.020 | TRUE |
| *SEMA4F* | P2 | 0.00 | 0.02 | 0.01 | 3.98 | 0.000 | 0.000 | TRUE |
| *SLC6A9* | P2 | 0.00 | 0.02 | 0.01 | 3.81 | 0.000 | 0.000 | TRUE |
| *GPR19* | P2 | 0.00 | 0.02 | 0.00 | 5.52 | 0.000 | 0.000 | TRUE |
| *KIR3DL2* | P2 | 0.00 | 0.01 | 0.00 | 4.39 | 0.000 | 0.000 | TRUE |
| *SCN8A* | P2 | 0.00 | 0.01 | 0.00 | 3.85 | 0.000 | 0.000 | TRUE |
| *ITGB4* | P2 | 0.00 | 0.02 | 0.01 | 3.60 | 0.000 | 0.001 | TRUE |
| *MAG* | P2 | 0.00 | 0.01 | 0.00 | 3.35 | 0.001 | 0.002 | TRUE |
| *LRRC3B* | P2 | 0.00 | 0.02 | 0.00 | 2.84 | 0.005 | 0.011 | TRUE |
| *ADRA2B* | P2 | 0.00 | 0.05 | 0.03 | 3.05 | 0.002 | 0.006 | TRUE |
| *CD8B* | P2 | 0.00 | 0.01 | 0.00 | 3.04 | 0.002 | 0.006 | TRUE |
| *VSIG8* | P2 | 0.00 | 0.02 | 0.01 | 2.49 | 0.013 | 0.028 | TRUE |
| *IL12RB2* | P2 | 0.00 | 0.03 | 0.01 | 3.45 | 0.001 | 0.001 | TRUE |
| *IL2RB* | P2 | 0.00 | 0.05 | 0.02 | 3.05 | 0.002 | 0.006 | TRUE |
| *ALK* | P2 | 0.00 | 0.01 | 0.00 | 2.28 | 0.023 | 0.049 | TRUE |
| *TNFRSF10C* | P2 | 0.00 | 0.02 | 0.01 | 4.50 | 0.000 | 0.000 | TRUE |
| *EFNA3* | P2 | 0.00 | 0.03 | 0.02 | 3.22 | 0.001 | 0.003 | TRUE |
| *SELP* | P2 | 0.00 | 0.01 | 0.00 | 2.72 | 0.007 | 0.016 | TRUE |
| *CACNG4* | P2 | 0.00 | 0.02 | 0.01 | 2.50 | 0.012 | 0.028 | TRUE |
| *SLC9A3* | P2 | 0.00 | 0.01 | 0.00 | 3.33 | 0.001 | 0.002 | TRUE |
| *SLC18A2* | P2 | 0.00 | 0.02 | 0.01 | 2.32 | 0.021 | 0.044 | TRUE |
| *OR10A2* | P2 | 0.00 | 0.01 | 0.00 | 2.52 | 0.012 | 0.027 | TRUE |
| *LRFN1* | P2 | 0.00 | 0.03 | 0.02 | 3.33 | 0.001 | 0.002 | TRUE |
| *RXFP2* | P2 | 0.00 | 0.01 | 0.00 | 2.78 | 0.005 | 0.013 | TRUE |
| *TAS2R20* | P2 | 0.00 | 0.01 | 0.00 | 2.30 | 0.022 | 0.046 | TRUE |
| *EPHA1* | P2 | 0.00 | 0.01 | 0.00 | 2.62 | 0.009 | 0.020 | TRUE |
| *OR4F15* | P2 | 0.00 | 0.01 | 0.00 | 2.57 | 0.010 | 0.023 | TRUE |
| *APLP1* | P2 | 0.00 | 0.01 | 0.01 | 3.01 | 0.003 | 0.006 | TRUE |
| *OR6A2* | P2 | 0.00 | 0.01 | 0.00 | 2.60 | 0.009 | 0.021 | TRUE |
| *AMIGO1* | P2 | 0.00 | 0.01 | 0.00 | 2.28 | 0.023 | 0.048 | TRUE |
| *SLC26A4* | P2 | 0.00 | 0.01 | 0.00 | 2.39 | 0.017 | 0.037 | TRUE |
| *OR8D4* | P2 | 0.00 | 0.01 | 0.00 | 2.40 | 0.016 | 0.036 | TRUE |
| *BEST2* | P2 | 0.00 | 0.01 | 0.00 | 3.32 | 0.001 | 0.002 | TRUE |
| *ASTN2* | P2 | 0.00 | 0.01 | 0.00 | 2.88 | 0.004 | 0.010 | TRUE |
| *GFRA3* | P2 | 0.00 | 0.01 | 0.00 | 2.56 | 0.010 | 0.023 | TRUE |
| *FLRT1* | P2 | 0.00 | 0.00 | 0.00 | 2.63 | 0.009 | 0.020 | TRUE |
| *ABCA12* | P2 | 0.00 | 0.00 | 0.00 | 2.28 | 0.022 | 0.048 | TRUE |
| *SLC4A4* | P2 | 0.00 | 0.00 | 0.00 | 2.30 | 0.021 | 0.046 | TRUE |
| *OR2AG1* | P2 | 0.00 | 0.00 | 0.00 | 2.37 | 0.018 | 0.038 | TRUE |
| *VSIG10L* | P2 | 0.00 | 0.01 | 0.01 | 2.49 | 0.013 | 0.029 | TRUE |
| *SLC5A2* | P2 | 0.00 | 0.00 | 0.00 | 2.26 | 0.024 | 0.050 | TRUE |
| *CD52* | P3 | 0.45 | 0.83 | 0.62 | 24.32 | 0.000 | 0.000 | TRUE |
| *SPN* | P3 | 0.39 | 0.90 | 0.63 | 26.38 | 0.000 | 0.000 | TRUE |
| *CD69* | P3 | 0.38 | 0.80 | 0.60 | 18.50 | 0.000 | 0.000 | TRUE |
| *GYPC* | P3 | 0.31 | 0.82 | 0.68 | 17.65 | 0.000 | 0.000 | TRUE |
| *LY6E* | P3 | 0.31 | 0.83 | 0.62 | 18.51 | 0.000 | 0.000 | TRUE |
| *CD63* | P3 | 0.30 | 0.98 | 0.90 | 16.78 | 0.000 | 0.000 | TRUE |
| *CD34* | P3 | 0.29 | 0.60 | 0.38 | 22.64 | 0.000 | 0.000 | TRUE |
| *RNF130* | P3 | 0.29 | 0.89 | 0.67 | 18.56 | 0.000 | 0.000 | TRUE |
| *CD96* | P3 | 0.28 | 0.68 | 0.42 | 22.87 | 0.000 | 0.000 | TRUE |
| *ITGA4* | P3 | 0.27 | 0.82 | 0.61 | 16.57 | 0.000 | 0.000 | TRUE |
| *BST2* | P3 | 0.24 | 0.90 | 0.77 | 13.26 | 0.000 | 0.000 | TRUE |
| *HLA-DRB1* | P3 | 0.23 | 0.90 | 0.81 | 10.82 | 0.000 | 0.000 | TRUE |
| *TM7SF3* | P3 | 0.23 | 0.67 | 0.40 | 21.86 | 0.000 | 0.000 | TRUE |
| *HLA-DMA* | P3 | 0.22 | 0.88 | 0.70 | 11.78 | 0.000 | 0.000 | TRUE |
| *ITM2B* | P3 | 0.22 | 0.99 | 0.91 | 11.01 | 0.000 | 0.000 | TRUE |
| *ATP6V0A2* | P3 | 0.21 | 0.54 | 0.37 | 16.56 | 0.000 | 0.000 | TRUE |
| *CD47* | P3 | 0.21 | 0.92 | 0.76 | 12.94 | 0.000 | 0.000 | TRUE |
| *IL2RG* | P3 | 0.20 | 0.85 | 0.66 | 14.38 | 0.000 | 0.000 | TRUE |
| *CD82* | P3 | 0.19 | 0.82 | 0.65 | 13.03 | 0.000 | 0.000 | TRUE |
| *TMEM219* | P3 | 0.19 | 0.85 | 0.65 | 13.79 | 0.000 | 0.000 | TRUE |
| *HLA-A* | P3 | 0.19 | 1.00 | 0.98 | 12.27 | 0.000 | 0.000 | TRUE |
| *SYPL1* | P3 | 0.18 | 0.70 | 0.49 | 16.16 | 0.000 | 0.000 | TRUE |
| *CD37* | P3 | 0.18 | 0.84 | 0.66 | 14.71 | 0.000 | 0.000 | TRUE |
| *ICAM3* | P3 | 0.17 | 0.66 | 0.49 | 17.19 | 0.000 | 0.000 | TRUE |
| *SPINT2* | P3 | 0.17 | 0.74 | 0.58 | 12.59 | 0.000 | 0.000 | TRUE |
| *M6PR* | P3 | 0.16 | 0.73 | 0.55 | 15.05 | 0.000 | 0.000 | TRUE |
| *SELL* | P3 | 0.16 | 0.45 | 0.31 | 13.88 | 0.000 | 0.000 | TRUE |
| *ATRAID* | P3 | 0.16 | 0.69 | 0.48 | 15.46 | 0.000 | 0.000 | TRUE |
| *ITM2C* | P3 | 0.16 | 0.63 | 0.46 | 12.71 | 0.000 | 0.000 | TRUE |
| *F2R* | P3 | 0.16 | 0.29 | 0.12 | 18.78 | 0.000 | 0.000 | TRUE |
| *IL1RL1* | P3 | 0.15 | 0.15 | 0.05 | 12.20 | 0.000 | 0.000 | TRUE |
| *STT3B* | P3 | 0.15 | 0.80 | 0.66 | 12.25 | 0.000 | 0.000 | TRUE |
| *HLA-E* | P3 | 0.15 | 0.99 | 0.93 | 8.29 | 0.000 | 0.000 | TRUE |
| *S1PR4* | P3 | 0.14 | 0.55 | 0.36 | 16.82 | 0.000 | 0.000 | TRUE |
| *TMEM179B* | P3 | 0.14 | 0.59 | 0.41 | 13.75 | 0.000 | 0.000 | TRUE |
| *PTTG1IP* | P3 | 0.14 | 0.73 | 0.56 | 11.33 | 0.000 | 0.000 | TRUE |
| *HLA-C* | P3 | 0.14 | 1.00 | 0.96 | 5.81 | 0.000 | 0.000 | TRUE |
| *ICAM2* | P3 | 0.14 | 0.53 | 0.35 | 15.00 | 0.000 | 0.000 | TRUE |
| *CD9* | P3 | 0.14 | 0.32 | 0.17 | 15.27 | 0.000 | 0.000 | TRUE |
| *KIT* | P3 | 0.13 | 0.33 | 0.18 | 15.17 | 0.000 | 0.000 | TRUE |

| | | | | | | | | |
|---|---|---|---|---|---|---|---|---|
| *SORL1* | P3 | 0.12 | 0.46 | 0.31 | 13.87 | 0.000 | 0.000 | TRUE |
| *SERINC3* | P3 | 0.12 | 0.61 | 0.43 | 13.39 | 0.000 | 0.000 | TRUE |
| *ABCC4* | P3 | 0.12 | 0.24 | 0.11 | 14.47 | 0.000 | 0.000 | TRUE |
| *HLA-DRB5* | P3 | 0.12 | 0.68 | 0.59 | 8.62 | 0.000 | 0.000 | TRUE |
| *SLC9A7* | P3 | 0.12 | 0.45 | 0.29 | 13.62 | 0.000 | 0.000 | TRUE |
| *PROM1* | P3 | 0.12 | 0.36 | 0.19 | 16.37 | 0.000 | 0.000 | TRUE |
| *TGOLN2* | P3 | 0.12 | 0.81 | 0.66 | 8.29 | 0.000 | 0.000 | TRUE |
| *NPR3* | P3 | 0.11 | 0.26 | 0.13 | 14.61 | 0.000 | 0.000 | TRUE |
| *SLC39A8* | P3 | 0.11 | 0.45 | 0.30 | 12.64 | 0.000 | 0.000 | TRUE |
| *TNFRSF14* | P3 | 0.11 | 0.63 | 0.46 | 11.61 | 0.000 | 0.000 | TRUE |
| *ABCC1* | P3 | 0.11 | 0.51 | 0.34 | 12.42 | 0.000 | 0.000 | TRUE |
| *CD164* | P3 | 0.11 | 0.91 | 0.78 | 6.78 | 0.000 | 0.000 | TRUE |
| *CD84* | P3 | 0.11 | 0.36 | 0.22 | 13.01 | 0.000 | 0.000 | TRUE |
| *CD79B* | P3 | 0.11 | 0.31 | 0.16 | 18.13 | 0.000 | 0.000 | TRUE |
| *TFPI* | P3 | 0.11 | 0.46 | 0.32 | 9.97 | 0.000 | 0.000 | TRUE |
| *NTRK1* | P3 | 0.10 | 0.15 | 0.03 | 15.53 | 0.000 | 0.000 | TRUE |
| *CD33* | P3 | 0.10 | 0.40 | 0.27 | 11.14 | 0.000 | 0.000 | TRUE |
| *IL1RAP* | P3 | 0.10 | 0.44 | 0.31 | 11.74 | 0.000 | 0.000 | TRUE |
| *IL9R* | P3 | 0.10 | 0.23 | 0.10 | 14.38 | 0.000 | 0.000 | TRUE |
| *ITGA2B* | P3 | 0.09 | 0.15 | 0.05 | 13.35 | 0.000 | 0.000 | TRUE |
| *PTPRA* | P3 | 0.09 | 0.65 | 0.50 | 8.80 | 0.000 | 0.000 | TRUE |
| *MFSD12* | P3 | 0.09 | 0.44 | 0.31 | 10.07 | 0.000 | 0.000 | TRUE |
| *F11R* | P3 | 0.09 | 0.50 | 0.35 | 11.81 | 0.000 | 0.000 | TRUE |
| *LAMP2* | P3 | 0.09 | 0.76 | 0.62 | 7.40 | 0.000 | 0.000 | TRUE |
| *BTN3A2* | P3 | 0.09 | 0.37 | 0.24 | 12.37 | 0.000 | 0.000 | TRUE |
| *PKD2* | P3 | 0.09 | 0.32 | 0.20 | 11.21 | 0.000 | 0.000 | TRUE |
| *SPNS3* | P3 | 0.09 | 0.31 | 0.24 | 6.55 | 0.000 | 0.000 | TRUE |
| *APP* | P3 | 0.08 | 0.48 | 0.35 | 8.83 | 0.000 | 0.000 | TRUE |
| *PIGT* | P3 | 0.08 | 0.43 | 0.30 | 11.14 | 0.000 | 0.000 | TRUE |
| *TGFBR2* | P3 | 0.08 | 0.39 | 0.26 | 11.59 | 0.000 | 0.000 | TRUE |
| *CD244* | P3 | 0.08 | 0.28 | 0.16 | 14.64 | 0.000 | 0.000 | TRUE |
| *SIGIRR* | P3 | 0.08 | 0.43 | 0.30 | 11.15 | 0.000 | 0.000 | TRUE |
| *P2RX1* | P3 | 0.08 | 0.36 | 0.22 | 13.99 | 0.000 | 0.000 | TRUE |
| *ATP1B1* | P3 | 0.08 | 0.31 | 0.20 | 12.12 | 0.000 | 0.000 | TRUE |
| *P2RY8* | P3 | 0.08 | 0.34 | 0.23 | 9.93 | 0.000 | 0.000 | TRUE |
| *SEMA7A* | P3 | 0.08 | 0.24 | 0.14 | 10.31 | 0.000 | 0.000 | TRUE |
| *CD46* | P3 | 0.08 | 0.74 | 0.60 | 6.83 | 0.000 | 0.000 | TRUE |
| *SLC40A1* | P3 | 0.07 | 0.14 | 0.07 | 7.93 | 0.000 | 0.000 | TRUE |
| *SLC39A10* | P3 | 0.07 | 0.43 | 0.31 | 8.59 | 0.000 | 0.000 | TRUE |
| *CD300A* | P3 | 0.07 | 0.24 | 0.14 | 11.67 | 0.000 | 0.000 | TRUE |
| *SLC12A6* | P3 | 0.07 | 0.47 | 0.36 | 7.92 | 0.000 | 0.000 | TRUE |
| *TRPV2* | P3 | 0.07 | 0.26 | 0.15 | 12.64 | 0.000 | 0.000 | TRUE |
| *SV2A* | P3 | 0.07 | 0.25 | 0.15 | 10.92 | 0.000 | 0.000 | TRUE |
| *FAM171A1* | P3 | 0.07 | 0.22 | 0.11 | 9.67 | 0.000 | 0.000 | TRUE |
| *SLC17A9* | P3 | 0.06 | 0.25 | 0.16 | 11.93 | 0.000 | 0.000 | TRUE |
| *GLIPR1* | P3 | 0.06 | 0.49 | 0.44 | 4.14 | 0.000 | 0.000 | TRUE |
| *RNFT1* | P3 | 0.06 | 0.28 | 0.16 | 10.49 | 0.000 | 0.000 | TRUE |
| *TMCO3* | P3 | 0.06 | 0.52 | 0.41 | 8.29 | 0.000 | 0.000 | TRUE |
| *TM4SF1* | P3 | 0.06 | 0.22 | 0.16 | 7.16 | 0.000 | 0.000 | TRUE |
| *TMIGD2* | P3 | 0.06 | 0.21 | 0.12 | 9.96 | 0.000 | 0.000 | TRUE |
| *GPR174* | P3 | 0.06 | 0.34 | 0.22 | 11.54 | 0.000 | 0.000 | TRUE |
| *TNFRSF1A* | P3 | 0.06 | 0.43 | 0.33 | 7.17 | 0.000 | 0.000 | TRUE |
| *CLN3* | P3 | 0.05 | 0.35 | 0.24 | 8.69 | 0.000 | 0.000 | TRUE |
| *ITFG1* | P3 | 0.05 | 0.32 | 0.22 | 9.70 | 0.000 | 0.000 | TRUE |
| *ATP2B4* | P3 | 0.05 | 0.30 | 0.21 | 6.66 | 0.000 | 0.000 | TRUE |
| *SEMA4D* | P3 | 0.05 | 0.34 | 0.24 | 8.05 | 0.000 | 0.000 | TRUE |
| *AMN* | P3 | 0.05 | 0.21 | 0.13 | 9.12 | 0.000 | 0.000 | TRUE |
| *EPHB6* | P3 | 0.05 | 0.21 | 0.12 | 10.81 | 0.000 | 0.000 | TRUE |
| *SUSD1* | P3 | 0.05 | 0.29 | 0.20 | 9.55 | 0.000 | 0.000 | TRUE |
| *NRROS* | P3 | 0.05 | 0.35 | 0.24 | 6.68 | 0.000 | 0.000 | TRUE |
| *TMEM106B* | P3 | 0.05 | 0.21 | 0.12 | 10.02 | 0.000 | 0.000 | TRUE |
| *TSPAN31* | P3 | 0.05 | 0.29 | 0.19 | 9.42 | 0.000 | 0.000 | TRUE |
| *GP1BB* | P3 | 0.05 | 0.12 | 0.05 | 8.97 | 0.000 | 0.000 | TRUE |
| *ITGA9* | P3 | 0.05 | 0.17 | 0.09 | 11.10 | 0.000 | 0.000 | TRUE |
| *NCSTN* | P3 | 0.05 | 0.29 | 0.20 | 7.71 | 0.000 | 0.000 | TRUE |
| *CDCP1* | P3 | 0.05 | 0.17 | 0.12 | 6.51 | 0.000 | 0.000 | TRUE |
| *STIM1* | P3 | 0.05 | 0.31 | 0.21 | 9.27 | 0.000 | 0.000 | TRUE |
| *SLC37A1* | P3 | 0.05 | 0.34 | 0.25 | 6.20 | 0.000 | 0.000 | TRUE |
| *PLXNA1* | P3 | 0.05 | 0.12 | 0.07 | 5.55 | 0.000 | 0.000 | TRUE |
| *TMEM63A* | P3 | 0.04 | 0.22 | 0.13 | 8.36 | 0.000 | 0.000 | TRUE |
| *RXFP1* | P3 | 0.04 | 0.13 | 0.07 | 5.58 | 0.000 | 0.000 | TRUE |
| *LTBR* | P3 | 0.04 | 0.33 | 0.24 | 7.58 | 0.000 | 0.000 | TRUE |
| *CTNS* | P3 | 0.04 | 0.18 | 0.11 | 7.22 | 0.000 | 0.000 | TRUE |
| *NOTCH1* | P3 | 0.04 | 0.29 | 0.21 | 8.89 | 0.000 | 0.000 | TRUE |
| *ICAM4* | P3 | 0.04 | 0.20 | 0.13 | 8.21 | 0.000 | 0.000 | TRUE |
| *KCNMB3* | P3 | 0.04 | 0.13 | 0.07 | 8.55 | 0.000 | 0.000 | TRUE |
| *UMODL1* | P3 | 0.04 | 0.16 | 0.11 | 6.28 | 0.000 | 0.000 | TRUE |
| *LRRC37B* | P3 | 0.04 | 0.23 | 0.16 | 5.56 | 0.000 | 0.000 | TRUE |
| *LRP6* | P3 | 0.04 | 0.17 | 0.09 | 3.98 | 0.000 | 0.000 | TRUE |
| *ERMP1* | P3 | 0.04 | 0.15 | 0.09 | 7.26 | 0.000 | 0.000 | TRUE |
| *SIGLEC6* | P3 | 0.04 | 0.08 | 0.03 | 6.64 | 0.000 | 0.000 | TRUE |
| *TMEM150A* | P3 | 0.04 | 0.14 | 0.07 | 8.87 | 0.000 | 0.000 | TRUE |
| *TM9SF4* | P3 | 0.04 | 0.24 | 0.16 | 9.27 | 0.000 | 0.000 | TRUE |
| *TXNDC15* | P3 | 0.04 | 0.22 | 0.15 | 10.13 | 0.000 | 0.000 | TRUE |
| *SLC24A3* | P3 | 0.04 | 0.09 | 0.02 | 6.78 | 0.000 | 0.000 | TRUE |
| *GPR12* | P3 | 0.04 | 0.09 | 0.04 | 3.29 | 0.001 | 0.003 | TRUE |
| *C3orf80* | P3 | 0.04 | 0.15 | 0.10 | 7.42 | 0.000 | 0.000 | TRUE |
| *TNFRSF18* | P3 | 0.04 | 0.23 | 0.18 | 6.99 | 0.000 | 0.000 | TRUE |
| *ATP13A2* | P3 | 0.04 | 0.15 | 0.09 | 8.53 | 0.000 | 0.000 | TRUE |
| *ADCY6* | P3 | 0.03 | 0.13 | 0.07 | 9.29 | 0.000 | 0.000 | TRUE |
| *THSD7A* | P3 | 0.03 | 0.08 | 0.02 | 3.78 | 0.000 | 0.000 | TRUE |
| *RYK* | P3 | 0.03 | 0.24 | 0.17 | 7.32 | 0.000 | 0.000 | TRUE |
| *GRIK5* | P3 | 0.03 | 0.12 | 0.06 | 8.83 | 0.000 | 0.000 | TRUE |
| *CD7* | P3 | 0.03 | 0.13 | 0.08 | 5.56 | 0.000 | 0.000 | TRUE |
| *HTR1F* | P3 | 0.03 | 0.13 | 0.07 | 9.76 | 0.000 | 0.000 | TRUE |
| *GJA1* | P3 | 0.03 | 0.09 | 0.05 | 7.51 | 0.000 | 0.000 | TRUE |
| *SUCNR1* | P3 | 0.03 | 0.18 | 0.13 | 5.29 | 0.000 | 0.000 | TRUE |
| *CASD1* | P3 | 0.03 | 0.16 | 0.10 | 7.97 | 0.000 | 0.000 | TRUE |
| *MFSD6* | P3 | 0.03 | 0.20 | 0.13 | 8.60 | 0.000 | 0.000 | TRUE |
| *SVOPL* | P3 | 0.03 | 0.12 | 0.07 | 7.86 | 0.000 | 0.000 | TRUE |
| *CLEC14A* | P3 | 0.03 | 0.15 | 0.10 | 5.62 | 0.000 | 0.000 | TRUE |

| | | | | | | | | |
|---|---|---|---|---|---|---|---|---|
| *SLC33A1* | P3 | 0.03 | 0.23 | 0.16 | 7.58 | 0.000 | 0.000 | TRUE |
| *CHRNB1* | P3 | 0.03 | 0.23 | 0.17 | 7.38 | 0.000 | 0.000 | TRUE |
| *NIPAL2* | P3 | 0.03 | 0.11 | 0.06 | 8.49 | 0.000 | 0.000 | TRUE |
| *ACE* | P3 | 0.03 | 0.09 | 0.04 | 7.48 | 0.000 | 0.000 | TRUE |
| *GP1BA* | P3 | 0.03 | 0.08 | 0.04 | 5.23 | 0.000 | 0.000 | TRUE |
| *LPAR5* | P3 | 0.03 | 0.07 | 0.03 | 8.35 | 0.000 | 0.000 | TRUE |
| *SPNS2* | P3 | 0.03 | 0.16 | 0.11 | 5.57 | 0.000 | 0.000 | TRUE |
| *ROBO3* | P3 | 0.03 | 0.18 | 0.13 | 5.61 | 0.000 | 0.000 | TRUE |
| *BACE2* | P3 | 0.03 | 0.11 | 0.07 | 3.90 | 0.000 | 0.000 | TRUE |
| *SLC46A3* | P3 | 0.03 | 0.11 | 0.06 | 4.95 | 0.000 | 0.000 | TRUE |
| *SLC26A2* | P3 | 0.03 | 0.17 | 0.12 | 6.36 | 0.000 | 0.000 | TRUE |
| *SLC39A9* | P3 | 0.03 | 0.18 | 0.12 | 6.62 | 0.000 | 0.000 | TRUE |
| *LRFN4* | P3 | 0.03 | 0.14 | 0.09 | 7.04 | 0.000 | 0.000 | TRUE |
| *CD79A* | P3 | 0.03 | 0.12 | 0.08 | 5.78 | 0.000 | 0.000 | TRUE |
| *FAM174A* | P3 | 0.03 | 0.12 | 0.07 | 8.63 | 0.000 | 0.000 | TRUE |
| *CLDN15* | P3 | 0.02 | 0.18 | 0.12 | 6.83 | 0.000 | 0.000 | TRUE |
| *BTN3A3* | P3 | 0.02 | 0.14 | 0.09 | 7.73 | 0.000 | 0.000 | TRUE |
| *SLC44A2* | P3 | 0.02 | 0.13 | 0.08 | 5.18 | 0.000 | 0.000 | TRUE |
| *PSEN2* | P3 | 0.02 | 0.09 | 0.05 | 4.23 | 0.000 | 0.000 | TRUE |
| *SEMA6A* | P3 | 0.02 | 0.03 | 0.01 | 4.06 | 0.000 | 0.000 | TRUE |
| *ZFYVE27* | P3 | 0.02 | 0.18 | 0.13 | 5.34 | 0.000 | 0.000 | TRUE |
| *TIE1* | P3 | 0.02 | 0.07 | 0.03 | 8.34 | 0.000 | 0.000 | TRUE |
| *NAGPA* | P3 | 0.02 | 0.15 | 0.11 | 6.38 | 0.000 | 0.000 | TRUE |
| *BTN3A1* | P3 | 0.02 | 0.15 | 0.10 | 6.25 | 0.000 | 0.000 | TRUE |
| *MFSD8* | P3 | 0.02 | 0.15 | 0.10 | 6.82 | 0.000 | 0.000 | TRUE |
| *F2RL3* | P3 | 0.02 | 0.05 | 0.01 | 7.62 | 0.000 | 0.000 | TRUE |
| *HLA-DQA2* | P3 | 0.02 | 0.10 | 0.07 | 3.66 | 0.000 | 0.001 | TRUE |
| *NTNG2* | P3 | 0.02 | 0.11 | 0.07 | 6.41 | 0.000 | 0.000 | TRUE |
| *ACVR1* | P3 | 0.02 | 0.11 | 0.07 | 7.35 | 0.000 | 0.000 | TRUE |
| *TSPAN2* | P3 | 0.02 | 0.08 | 0.05 | 5.28 | 0.000 | 0.000 | TRUE |
| *ABCC5* | P3 | 0.02 | 0.12 | 0.08 | 5.82 | 0.000 | 0.000 | TRUE |
| *TRAT1* | P3 | 0.02 | 0.08 | 0.03 | 5.23 | 0.000 | 0.000 | TRUE |
| *CDH26* | P3 | 0.02 | 0.08 | 0.05 | 3.83 | 0.000 | 0.000 | TRUE |
| *TNFRSF25* | P3 | 0.02 | 0.13 | 0.10 | 5.31 | 0.000 | 0.000 | TRUE |
| *FLT3LG* | P3 | 0.02 | 0.08 | 0.04 | 6.35 | 0.000 | 0.000 | TRUE |
| *TMEM25* | P3 | 0.02 | 0.07 | 0.04 | 5.42 | 0.000 | 0.000 | TRUE |
| *IL18R1* | P3 | 0.02 | 0.17 | 0.12 | 2.58 | 0.010 | 0.022 | TRUE |
| *S1PR1* | P3 | 0.02 | 0.06 | 0.03 | 3.37 | 0.001 | 0.002 | TRUE |
| *DPEP2* | P3 | 0.02 | 0.10 | 0.06 | 7.83 | 0.000 | 0.000 | TRUE |
| *IL12RB1* | P3 | 0.02 | 0.11 | 0.07 | 4.44 | 0.000 | 0.000 | TRUE |
| *IL17RE* | P3 | 0.02 | 0.08 | 0.05 | 7.50 | 0.000 | 0.000 | TRUE |
| *CDH9* | P3 | 0.02 | 0.04 | 0.02 | 2.33 | 0.020 | 0.042 | TRUE |
| *SLC12A4* | P3 | 0.02 | 0.10 | 0.07 | 4.28 | 0.000 | 0.000 | TRUE |
| *SLC2A11* | P3 | 0.02 | 0.09 | 0.06 | 6.52 | 0.000 | 0.000 | TRUE |
| *TBXA2R* | P3 | 0.02 | 0.04 | 0.01 | 6.80 | 0.000 | 0.000 | TRUE |
| *NAALADL1* | P3 | 0.02 | 0.06 | 0.03 | 4.36 | 0.000 | 0.000 | TRUE |
| *ACP2* | P3 | 0.02 | 0.09 | 0.06 | 4.02 | 0.000 | 0.000 | TRUE |
| *ATRN* | P3 | 0.02 | 0.13 | 0.09 | 4.36 | 0.000 | 0.000 | TRUE |
| *LMBRD2* | P3 | 0.02 | 0.09 | 0.06 | 5.90 | 0.000 | 0.000 | TRUE |
| *MEGF8* | P3 | 0.01 | 0.08 | 0.05 | 6.62 | 0.000 | 0.000 | TRUE |
| *TSPAN17* | P3 | 0.01 | 0.11 | 0.08 | 5.14 | 0.000 | 0.000 | TRUE |
| *MICA* | P3 | 0.01 | 0.12 | 0.09 | 4.75 | 0.000 | 0.000 | TRUE |
| *NRG4* | P3 | 0.01 | 0.08 | 0.06 | 3.01 | 0.003 | 0.006 | TRUE |
| *PLXNA3* | P3 | 0.01 | 0.09 | 0.06 | 5.79 | 0.000 | 0.000 | TRUE |
| *GPR146* | P3 | 0.01 | 0.08 | 0.05 | 6.13 | 0.000 | 0.000 | TRUE |
| *ZP3* | P3 | 0.01 | 0.07 | 0.04 | 4.86 | 0.000 | 0.000 | TRUE |
| *ICAM5* | P3 | 0.01 | 0.05 | 0.02 | 5.61 | 0.000 | 0.000 | TRUE |
| *SLC16A5* | P3 | 0.01 | 0.06 | 0.03 | 5.97 | 0.000 | 0.000 | TRUE |
| *TMEM182* | P3 | 0.01 | 0.05 | 0.03 | 6.80 | 0.000 | 0.000 | TRUE |
| *KIRREL3* | P3 | 0.01 | 0.03 | 0.00 | 6.09 | 0.000 | 0.000 | TRUE |
| *SLC35A5* | P3 | 0.01 | 0.09 | 0.06 | 5.60 | 0.000 | 0.000 | TRUE |
| *SLC29A3* | P3 | 0.01 | 0.08 | 0.05 | 4.13 | 0.000 | 0.000 | TRUE |
| *TMEM204* | P3 | 0.01 | 0.05 | 0.03 | 4.28 | 0.000 | 0.000 | TRUE |
| *MEGF10* | P3 | 0.01 | 0.04 | 0.02 | 2.76 | 0.006 | 0.014 | TRUE |
| *IL17RC* | P3 | 0.01 | 0.09 | 0.06 | 6.18 | 0.000 | 0.000 | TRUE |
| *SLC45A4* | P3 | 0.01 | 0.08 | 0.05 | 4.84 | 0.000 | 0.000 | TRUE |
| *TMEM140* | P3 | 0.01 | 0.14 | 0.10 | 3.43 | 0.001 | 0.002 | TRUE |
| *LTK* | P3 | 0.01 | 0.04 | 0.02 | 5.75 | 0.000 | 0.000 | TRUE |
| *LCT* | P3 | 0.01 | 0.04 | 0.03 | 3.17 | 0.002 | 0.004 | TRUE |
| *AQP2* | P3 | 0.01 | 0.01 | 0.00 | 3.18 | 0.001 | 0.004 | TRUE |
| *HLA-G* | P3 | 0.01 | 0.08 | 0.06 | 2.43 | 0.015 | 0.033 | TRUE |
| *DPEP1* | P3 | 0.01 | 0.07 | 0.05 | 3.50 | 0.000 | 0.001 | TRUE |
| *SLAMF6* | P3 | 0.01 | 0.06 | 0.03 | 4.43 | 0.000 | 0.000 | TRUE |
| *ABCC10* | P3 | 0.01 | 0.06 | 0.04 | 3.87 | 0.000 | 0.000 | TRUE |
| *MFSD2B* | P3 | 0.01 | 0.04 | 0.02 | 4.97 | 0.000 | 0.000 | TRUE |
| *PTGER1* | P3 | 0.01 | 0.03 | 0.01 | 3.77 | 0.000 | 0.000 | TRUE |
| *MFAP3L* | P3 | 0.01 | 0.02 | 0.00 | 4.28 | 0.000 | 0.000 | TRUE |
| *TMEM132E* | P3 | 0.01 | 0.05 | 0.03 | 2.70 | 0.007 | 0.016 | TRUE |
| *CADM2* | P3 | 0.01 | 0.03 | 0.01 | 4.38 | 0.000 | 0.000 | TRUE |
| *TREML2* | P3 | 0.01 | 0.03 | 0.02 | 2.69 | 0.007 | 0.017 | TRUE |
| *GABRD* | P3 | 0.01 | 0.03 | 0.02 | 2.56 | 0.010 | 0.024 | TRUE |
| *EDA2R* | P3 | 0.01 | 0.02 | 0.01 | 4.75 | 0.000 | 0.000 | TRUE |
| *CNNM2* | P3 | 0.01 | 0.08 | 0.06 | 3.73 | 0.000 | 0.001 | TRUE |
| *SLC5A10* | P3 | 0.01 | 0.02 | 0.01 | 2.61 | 0.009 | 0.021 | TRUE |
| *LRRC24* | P3 | 0.01 | 0.05 | 0.03 | 3.56 | 0.000 | 0.001 | TRUE |
| *TLR3* | P3 | 0.01 | 0.03 | 0.02 | 3.22 | 0.001 | 0.003 | TRUE |
| *MMEL1* | P3 | 0.01 | 0.04 | 0.03 | 4.40 | 0.000 | 0.000 | TRUE |
| *GPC2* | P3 | 0.01 | 0.03 | 0.01 | 3.47 | 0.001 | 0.001 | TRUE |
| *DUOX2* | P3 | 0.01 | 0.01 | 0.00 | 2.69 | 0.007 | 0.017 | TRUE |
| *ITGB5* | P3 | 0.01 | 0.02 | 0.01 | 3.85 | 0.000 | 0.000 | TRUE |
| *SLC38A11* | P3 | 0.01 | 0.01 | 0.00 | 4.25 | 0.000 | 0.000 | TRUE |
| *PTH2R* | P3 | 0.01 | 0.03 | 0.02 | 4.40 | 0.000 | 0.000 | TRUE |
| *MALRD1* | P3 | 0.00 | 0.04 | 0.03 | 2.57 | 0.010 | 0.023 | TRUE |
| *THSD1* | P3 | 0.00 | 0.03 | 0.02 | 2.85 | 0.004 | 0.011 | TRUE |
| *EFNA4* | P3 | 0.00 | 0.05 | 0.04 | 3.35 | 0.001 | 0.002 | TRUE |
| *MANSC1* | P3 | 0.00 | 0.03 | 0.02 | 4.22 | 0.000 | 0.000 | TRUE |
| *TAS2R14* | P3 | 0.00 | 0.04 | 0.03 | 3.16 | 0.002 | 0.004 | TRUE |
| *SLC14A1* | P3 | 0.00 | 0.04 | 0.03 | 3.03 | 0.002 | 0.006 | TRUE |
| *OR2C3* | P3 | 0.00 | 0.01 | 0.01 | 2.49 | 0.013 | 0.029 | TRUE |
| *CD24* | P3 | 0.00 | 0.02 | 0.01 | 2.66 | 0.008 | 0.018 | TRUE |
| *UNC5B* | P3 | 0.00 | 0.01 | 0.01 | 2.69 | 0.007 | 0.017 | TRUE |

| | | | | | | | | |
|---|---|---|---|---|---|---|---|---|
| *DISP2* | P3 | 0.00 | 0.02 | 0.01 | 2.53 | 0.011 | 0.025 | TRUE |
| *PTGDR2* | P3 | 0.00 | 0.01 | 0.00 | 3.23 | 0.001 | 0.003 | TRUE |
| *XKR3* | P3 | 0.00 | 0.01 | 0.01 | 2.75 | 0.006 | 0.014 | TRUE |
| *BCAN* | P3 | 0.00 | 0.01 | 0.00 | 3.20 | 0.001 | 0.003 | TRUE |
| *SLC52A3* | P3 | 0.00 | 0.01 | 0.01 | 3.80 | 0.000 | 0.000 | TRUE |
| *PCDHGB2* | P3 | 0.00 | 0.01 | 0.00 | 2.76 | 0.006 | 0.014 | TRUE |
| *PLXNB1* | P3 | 0.00 | 0.02 | 0.02 | 2.90 | 0.004 | 0.009 | TRUE |
| *AMHR2* | P3 | 0.00 | 0.00 | 0.00 | 3.36 | 0.001 | 0.002 | TRUE |
| *SLC26A1* | P3 | 0.00 | 0.01 | 0.01 | 3.18 | 0.001 | 0.004 | TRUE |
| *SLC2A14* | P3 | 0.00 | 0.02 | 0.01 | 4.01 | 0.000 | 0.000 | TRUE |
| *SCN4A* | P3 | 0.00 | 0.01 | 0.00 | 2.64 | 0.008 | 0.019 | TRUE |
| *LGR6* | P3 | 0.00 | 0.01 | 0.01 | 2.60 | 0.009 | 0.021 | TRUE |
| *IL5RA* | P3 | 0.00 | 0.01 | 0.00 | 2.45 | 0.014 | 0.032 | TRUE |
| *SLC10A1* | P3 | 0.00 | 0.01 | 0.00 | 3.37 | 0.001 | 0.002 | TRUE |
| *DUOXA1* | P3 | 0.00 | 0.01 | 0.00 | 3.16 | 0.002 | 0.004 | TRUE |
| *CHRNA7* | P3 | 0.00 | 0.01 | 0.01 | 2.46 | 0.014 | 0.031 | TRUE |
| *OR52H1* | P3 | 0.00 | 0.00 | 0.00 | 2.59 | 0.010 | 0.022 | TRUE |
| *GLP1R* | P3 | 0.00 | 0.00 | 0.00 | 2.40 | 0.017 | 0.036 | TRUE |
| *ADCY2* | P3 | 0.00 | 0.01 | 0.00 | 2.31 | 0.021 | 0.045 | TRUE |
| *HRH4* | P3 | 0.00 | 0.00 | 0.00 | 2.72 | 0.006 | 0.015 | TRUE |
| *GPR171* | P3 | 0.00 | 0.10 | 0.07 | 3.32 | 0.001 | 0.002 | TRUE |
| *NIPAL1* | P3 | 0.00 | 0.00 | 0.00 | 2.28 | 0.023 | 0.048 | TRUE |
| *LAMP3* | P4 | 0.73 | 0.52 | 0.07 | 29.79 | 0.000 | 0.000 | TRUE |
| *CCR7* | P4 | 0.49 | 0.45 | 0.23 | 24.27 | 0.000 | 0.000 | TRUE |
| *CD40* | P4 | 0.44 | 0.59 | 0.36 | 25.95 | 0.000 | 0.000 | TRUE |
| *HLA-DPA1* | P4 | 0.37 | 0.91 | 0.83 | 12.17 | 0.000 | 0.000 | TRUE |
| *HLA-DPB1* | P4 | 0.32 | 0.91 | 0.85 | 8.87 | 0.000 | 0.000 | TRUE |
| *HLA-DQB1* | P4 | 0.32 | 0.77 | 0.64 | 13.32 | 0.000 | 0.000 | TRUE |
| *CD74* | P4 | 0.32 | 0.97 | 0.94 | 8.64 | 0.000 | 0.000 | TRUE |
| *SLAMF7* | P4 | 0.30 | 0.31 | 0.10 | 19.56 | 0.000 | 0.000 | TRUE |
| *HLA-DRA* | P4 | 0.29 | 0.95 | 0.93 | 10.40 | 0.000 | 0.000 | TRUE |
| *HLA-DQA1* | P4 | 0.29 | 0.71 | 0.52 | 14.50 | 0.000 | 0.000 | TRUE |
| *VCAM1* | P4 | 0.28 | 0.41 | 0.17 | 15.93 | 0.000 | 0.000 | TRUE |
| *IL2RA* | P4 | 0.27 | 0.45 | 0.26 | 18.00 | 0.000 | 0.000 | TRUE |
| *ANTXR2* | P4 | 0.26 | 0.67 | 0.41 | 18.78 | 0.000 | 0.000 | TRUE |
| *SLAMF1* | P4 | 0.25 | 0.39 | 0.14 | 19.40 | 0.000 | 0.000 | TRUE |
| *HLA-DMB* | P4 | 0.25 | 0.55 | 0.34 | 15.32 | 0.000 | 0.000 | TRUE |
| *ADAM28* | P4 | 0.24 | 0.47 | 0.23 | 20.47 | 0.000 | 0.000 | TRUE |
| *AREG* | P4 | 0.24 | 0.36 | 0.27 | 6.96 | 0.000 | 0.000 | TRUE |
| *PTGER4* | P4 | 0.24 | 0.74 | 0.55 | 16.71 | 0.000 | 0.000 | TRUE |
| *LY75* | P4 | 0.23 | 0.51 | 0.30 | 18.20 | 0.000 | 0.000 | TRUE |
| *SLC38A1* | P4 | 0.23 | 0.77 | 0.58 | 11.80 | 0.000 | 0.000 | TRUE |
| *NRP2* | P4 | 0.23 | 0.29 | 0.06 | 19.87 | 0.000 | 0.000 | TRUE |
| *SPPL2A* | P4 | 0.22 | 0.70 | 0.49 | 16.53 | 0.000 | 0.000 | TRUE |
| *ALCAM* | P4 | 0.22 | 0.51 | 0.28 | 17.94 | 0.000 | 0.000 | TRUE |
| *SLC5A3* | P4 | 0.21 | 0.51 | 0.38 | 10.04 | 0.000 | 0.000 | TRUE |
| *IL7R* | P4 | 0.20 | 0.25 | 0.08 | 17.90 | 0.000 | 0.000 | TRUE |
| *TMEM123* | P4 | 0.20 | 0.89 | 0.78 | 10.58 | 0.000 | 0.000 | TRUE |
| *PTGIR* | P4 | 0.20 | 0.47 | 0.35 | 11.67 | 0.000 | 0.000 | TRUE |
| *TSPAN33* | P4 | 0.20 | 0.41 | 0.20 | 18.52 | 0.000 | 0.000 | TRUE |
| *IL3RA* | P4 | 0.18 | 0.69 | 0.55 | 11.31 | 0.000 | 0.000 | TRUE |
| *CD274* | P4 | 0.18 | 0.32 | 0.15 | 16.72 | 0.000 | 0.000 | TRUE |
| *SLCO5A1* | P4 | 0.18 | 0.35 | 0.12 | 17.34 | 0.000 | 0.000 | TRUE |
| *CD58* | P4 | 0.17 | 0.57 | 0.39 | 15.67 | 0.000 | 0.000 | TRUE |
| *TNFRSF9* | P4 | 0.16 | 0.39 | 0.23 | 14.68 | 0.000 | 0.000 | TRUE |
| *FLT3* | P4 | 0.16 | 0.42 | 0.27 | 14.28 | 0.000 | 0.000 | TRUE |
| *HLA-F* | P4 | 0.16 | 0.58 | 0.46 | 10.66 | 0.000 | 0.000 | TRUE |
| *ITGB8* | P4 | 0.15 | 0.18 | 0.05 | 14.42 | 0.000 | 0.000 | TRUE |
| *MPZL1* | P4 | 0.14 | 0.51 | 0.32 | 14.01 | 0.000 | 0.000 | TRUE |
| *IL21R* | P4 | 0.14 | 0.35 | 0.17 | 15.78 | 0.000 | 0.000 | TRUE |
| *FAS* | P4 | 0.13 | 0.37 | 0.21 | 13.18 | 0.000 | 0.000 | TRUE |
| *IL15RA* | P4 | 0.13 | 0.41 | 0.26 | 13.46 | 0.000 | 0.000 | TRUE |
| *LDLRAD4* | P4 | 0.13 | 0.35 | 0.25 | 7.29 | 0.000 | 0.000 | TRUE |
| *IL6ST* | P4 | 0.13 | 0.48 | 0.30 | 12.65 | 0.000 | 0.000 | TRUE |
| *SUCO* | P4 | 0.12 | 0.59 | 0.42 | 11.07 | 0.000 | 0.000 | TRUE |
| *RHBDF2* | P4 | 0.12 | 0.35 | 0.18 | 13.92 | 0.000 | 0.000 | TRUE |
| *TM9SF3* | P4 | 0.12 | 0.83 | 0.70 | 6.78 | 0.000 | 0.000 | TRUE |
| *PMEPA1* | P4 | 0.12 | 0.22 | 0.06 | 16.66 | 0.000 | 0.000 | TRUE |
| *IFNAR2* | P4 | 0.12 | 0.52 | 0.33 | 13.02 | 0.000 | 0.000 | TRUE |
| *PVR* | P4 | 0.12 | 0.23 | 0.08 | 15.24 | 0.000 | 0.000 | TRUE |
| *SLC15A4* | P4 | 0.11 | 0.33 | 0.17 | 13.07 | 0.000 | 0.000 | TRUE |
| *CD80* | P4 | 0.11 | 0.17 | 0.02 | 17.18 | 0.000 | 0.000 | TRUE |
| *LRP12* | P4 | 0.11 | 0.46 | 0.33 | 11.46 | 0.000 | 0.000 | TRUE |
| *CD109* | P4 | 0.11 | 0.38 | 0.27 | 8.37 | 0.000 | 0.000 | TRUE |
| *LNPEP* | P4 | 0.11 | 0.60 | 0.44 | 8.30 | 0.000 | 0.000 | TRUE |
| *ICOSLG* | P4 | 0.11 | 0.29 | 0.14 | 13.28 | 0.000 | 0.000 | TRUE |
| *CD1C* | P4 | 0.11 | 0.10 | 0.01 | 9.21 | 0.000 | 0.000 | TRUE |
| *CD200* | P4 | 0.11 | 0.22 | 0.09 | 12.92 | 0.000 | 0.000 | TRUE |
| *ADAM19* | P4 | 0.10 | 0.23 | 0.09 | 14.67 | 0.000 | 0.000 | TRUE |
| *ITGB7* | P4 | 0.10 | 0.20 | 0.10 | 7.25 | 0.000 | 0.000 | TRUE |
| *RELL1* | P4 | 0.10 | 0.41 | 0.29 | 10.36 | 0.000 | 0.000 | TRUE |
| *CD70* | P4 | 0.10 | 0.19 | 0.08 | 11.80 | 0.000 | 0.000 | TRUE |
| *SCARB2* | P4 | 0.10 | 0.50 | 0.35 | 9.44 | 0.000 | 0.000 | TRUE |
| *TMEM30A* | P4 | 0.10 | 0.61 | 0.46 | 7.71 | 0.000 | 0.000 | TRUE |
| *TM9SF2* | P4 | 0.09 | 0.68 | 0.53 | 6.68 | 0.000 | 0.000 | TRUE |
| *IFNAR1* | P4 | 0.09 | 0.57 | 0.43 | 8.71 | 0.000 | 0.000 | TRUE |
| *CLDND1* | P4 | 0.09 | 0.62 | 0.47 | 8.01 | 0.000 | 0.000 | TRUE |
| *PLXND1* | P4 | 0.09 | 0.36 | 0.22 | 11.62 | 0.000 | 0.000 | TRUE |
| *HLA-DOB* | P4 | 0.09 | 0.15 | 0.03 | 16.07 | 0.000 | 0.000 | TRUE |
| *ECE1* | P4 | 0.09 | 0.20 | 0.08 | 13.02 | 0.000 | 0.000 | TRUE |
| *SSTR2* | P4 | 0.08 | 0.24 | 0.13 | 10.02 | 0.000 | 0.000 | TRUE |
| *LPAR6* | P4 | 0.08 | 0.26 | 0.15 | 9.21 | 0.000 | 0.000 | TRUE |
| *TMEM106A* | P4 | 0.08 | 0.38 | 0.24 | 9.59 | 0.000 | 0.000 | TRUE |
| *JAG1* | P4 | 0.08 | 0.16 | 0.06 | 10.50 | 0.000 | 0.000 | TRUE |
| *BTN2A2* | P4 | 0.08 | 0.34 | 0.21 | 8.91 | 0.000 | 0.000 | TRUE |
| *GJA4* | P4 | 0.07 | 0.09 | 0.02 | 9.72 | 0.000 | 0.000 | TRUE |
| *CYSLTR1* | P4 | 0.07 | 0.32 | 0.21 | 7.41 | 0.000 | 0.000 | TRUE |
| *LMBRD1* | P4 | 0.07 | 0.39 | 0.27 | 7.81 | 0.000 | 0.000 | TRUE |
| *ANO9* | P4 | 0.07 | 0.16 | 0.05 | 11.42 | 0.000 | 0.000 | TRUE |
| *HLA-DOA* | P4 | 0.07 | 0.23 | 0.13 | 10.07 | 0.000 | 0.000 | TRUE |
| *TMEM87A* | P4 | 0.06 | 0.55 | 0.42 | 6.31 | 0.000 | 0.000 | TRUE |

| | | | | | | | | |
|---|---|---|---|---|---|---|---|---|
| *GRAMD1B* | P4 | 0.06 | 0.19 | 0.08 | 10.75 | 0.000 | 0.000 | TRUE |
| *TGFBR1* | P4 | 0.06 | 0.35 | 0.24 | 6.12 | 0.000 | 0.000 | TRUE |
| *SLC41A2* | P4 | 0.06 | 0.15 | 0.04 | 14.41 | 0.000 | 0.000 | TRUE |
| *LMBR1L* | P4 | 0.06 | 0.37 | 0.25 | 7.89 | 0.000 | 0.000 | TRUE |
| *DLL4* | P4 | 0.06 | 0.08 | 0.01 | 10.23 | 0.000 | 0.000 | TRUE |
| *CLEC2D* | P4 | 0.06 | 0.21 | 0.10 | 10.66 | 0.000 | 0.000 | TRUE |
| *TNFSF4* | P4 | 0.06 | 0.18 | 0.11 | 7.32 | 0.000 | 0.000 | TRUE |
| *ACHE* | P4 | 0.06 | 0.12 | 0.03 | 11.79 | 0.000 | 0.000 | TRUE |
| *ITGAV* | P4 | 0.06 | 0.22 | 0.12 | 10.84 | 0.000 | 0.000 | TRUE |
| *ITGA1* | P4 | 0.06 | 0.23 | 0.14 | 6.53 | 0.000 | 0.000 | TRUE |
| *CD1E* | P4 | 0.06 | 0.07 | 0.00 | 6.80 | 0.000 | 0.000 | TRUE |
| *CRLF2* | P4 | 0.05 | 0.17 | 0.11 | 6.97 | 0.000 | 0.000 | TRUE |
| *BTN2A1* | P4 | 0.05 | 0.30 | 0.18 | 8.00 | 0.000 | 0.000 | TRUE |
| *GPR157* | P4 | 0.05 | 0.09 | 0.02 | 11.02 | 0.000 | 0.000 | TRUE |
| *GPR137B* | P4 | 0.05 | 0.11 | 0.03 | 12.20 | 0.000 | 0.000 | TRUE |
| *GPR107* | P4 | 0.05 | 0.28 | 0.18 | 9.51 | 0.000 | 0.000 | TRUE |
| *CALCRL* | P4 | 0.05 | 0.15 | 0.08 | 8.08 | 0.000 | 0.000 | TRUE |
| *SCARF1* | P4 | 0.05 | 0.26 | 0.17 | 7.80 | 0.000 | 0.000 | TRUE |
| *LILRA4* | P4 | 0.05 | 0.06 | 0.00 | 8.09 | 0.000 | 0.000 | TRUE |
| *P2RY10* | P4 | 0.05 | 0.14 | 0.07 | 11.03 | 0.000 | 0.000 | TRUE |
| *BMPR2* | P4 | 0.05 | 0.20 | 0.11 | 10.32 | 0.000 | 0.000 | TRUE |
| *MR1* | P4 | 0.05 | 0.29 | 0.21 | 7.15 | 0.000 | 0.000 | TRUE |
| *PDGFRA* | P4 | 0.05 | 0.14 | 0.05 | 8.81 | 0.000 | 0.000 | TRUE |
| *PTCRA* | P4 | 0.05 | 0.04 | 0.00 | 5.79 | 0.000 | 0.000 | TRUE |
| *PLXNC1* | P4 | 0.05 | 0.22 | 0.13 | 8.52 | 0.000 | 0.000 | TRUE |
| *CRIM1* | P4 | 0.04 | 0.14 | 0.08 | 7.87 | 0.000 | 0.000 | TRUE |
| *TNFRSF11A* | P4 | 0.04 | 0.09 | 0.01 | 13.79 | 0.000 | 0.000 | TRUE |
| *SLC2A1* | P4 | 0.04 | 0.24 | 0.17 | 5.34 | 0.000 | 0.000 | TRUE |
| *ITGA6* | P4 | 0.04 | 0.26 | 0.18 | 6.63 | 0.000 | 0.000 | TRUE |
| *MS4A1* | P4 | 0.04 | 0.04 | 0.00 | 8.50 | 0.000 | 0.000 | TRUE |
| *LYSMD3* | P4 | 0.04 | 0.33 | 0.25 | 4.21 | 0.000 | 0.000 | TRUE |
| *ENPP4* | P4 | 0.04 | 0.13 | 0.08 | 3.80 | 0.000 | 0.000 | TRUE |
| *LAG3* | P4 | 0.04 | 0.22 | 0.15 | 2.93 | 0.003 | 0.008 | TRUE |
| *TPBG* | P4 | 0.04 | 0.18 | 0.11 | 5.84 | 0.000 | 0.000 | TRUE |
| *NIPAL3* | P4 | 0.04 | 0.22 | 0.15 | 6.16 | 0.000 | 0.000 | TRUE |
| *TMEM87B* | P4 | 0.04 | 0.30 | 0.22 | 6.66 | 0.000 | 0.000 | TRUE |
| *NEGR1* | P4 | 0.04 | 0.08 | 0.04 | 3.69 | 0.000 | 0.001 | TRUE |
| *SSPN* | P4 | 0.04 | 0.11 | 0.04 | 8.05 | 0.000 | 0.000 | TRUE |
| *SDK2* | P4 | 0.03 | 0.09 | 0.03 | 6.25 | 0.000 | 0.000 | TRUE |
| *ADCY9* | P4 | 0.03 | 0.11 | 0.05 | 6.59 | 0.000 | 0.000 | TRUE |
| *GPRC5C* | P4 | 0.03 | 0.20 | 0.15 | 3.95 | 0.000 | 0.000 | TRUE |
| *P2RY14* | P4 | 0.03 | 0.12 | 0.06 | 7.10 | 0.000 | 0.000 | TRUE |
| *MFSD11* | P4 | 0.03 | 0.25 | 0.17 | 6.28 | 0.000 | 0.000 | TRUE |
| *SLC8A3* | P4 | 0.03 | 0.09 | 0.02 | 8.42 | 0.000 | 0.000 | TRUE |
| *ACVR2A* | P4 | 0.03 | 0.08 | 0.02 | 8.94 | 0.000 | 0.000 | TRUE |
| *PAM* | P4 | 0.03 | 0.12 | 0.06 | 8.41 | 0.000 | 0.000 | TRUE |
| *TSPAN15* | P4 | 0.03 | 0.05 | 0.01 | 9.26 | 0.000 | 0.000 | TRUE |
| *ERVK13-1* | P4 | 0.03 | 0.16 | 0.10 | 6.37 | 0.000 | 0.000 | TRUE |
| *ACVRL1* | P4 | 0.03 | 0.08 | 0.03 | 8.00 | 0.000 | 0.000 | TRUE |
| *PRRT3* | P4 | 0.03 | 0.10 | 0.05 | 5.52 | 0.000 | 0.000 | TRUE |
| *SCN1B* | P4 | 0.03 | 0.09 | 0.04 | 7.58 | 0.000 | 0.000 | TRUE |
| *SLC6A12* | P4 | 0.03 | 0.05 | 0.00 | 9.14 | 0.000 | 0.000 | TRUE |
| *KIAA1324L* | P4 | 0.03 | 0.12 | 0.07 | 5.97 | 0.000 | 0.000 | TRUE |
| *FLVCR2* | P4 | 0.03 | 0.07 | 0.02 | 7.67 | 0.000 | 0.000 | TRUE |
| *PTPRM* | P4 | 0.03 | 0.12 | 0.08 | 2.99 | 0.003 | 0.007 | TRUE |
| *CHL1* | P4 | 0.03 | 0.05 | 0.00 | 5.13 | 0.000 | 0.000 | TRUE |
| *ACKR1* | P4 | 0.03 | 0.06 | 0.02 | 5.83 | 0.000 | 0.000 | TRUE |
| *EMP2* | P4 | 0.03 | 0.08 | 0.04 | 5.90 | 0.000 | 0.000 | TRUE |
| *CXCR3* | P4 | 0.03 | 0.07 | 0.03 | 6.46 | 0.000 | 0.000 | TRUE |
| *CD226* | P4 | 0.02 | 0.06 | 0.02 | 6.84 | 0.000 | 0.000 | TRUE |
| *SIDT2* | P4 | 0.02 | 0.13 | 0.09 | 4.00 | 0.000 | 0.000 | TRUE |
| *KCNMB1* | P4 | 0.02 | 0.08 | 0.04 | 6.50 | 0.000 | 0.000 | TRUE |
| *SLC12A7* | P4 | 0.02 | 0.19 | 0.14 | 3.65 | 0.000 | 0.001 | TRUE |
| *MC2R* | P4 | 0.02 | 0.05 | 0.00 | 5.13 | 0.000 | 0.000 | TRUE |
| *CCR6* | P4 | 0.02 | 0.04 | 0.00 | 6.00 | 0.000 | 0.000 | TRUE |
| *LY9* | P4 | 0.02 | 0.04 | 0.01 | 4.37 | 0.000 | 0.000 | TRUE |
| *MPZL3* | P4 | 0.02 | 0.08 | 0.03 | 5.05 | 0.000 | 0.000 | TRUE |
| *GABBR1* | P4 | 0.02 | 0.08 | 0.03 | 8.36 | 0.000 | 0.000 | TRUE |
| *ABCA5* | P4 | 0.02 | 0.09 | 0.05 | 3.80 | 0.000 | 0.000 | TRUE |
| *PGAP1* | P4 | 0.02 | 0.12 | 0.07 | 4.98 | 0.000 | 0.000 | TRUE |
| *CLSTN3* | P4 | 0.02 | 0.15 | 0.09 | 5.30 | 0.000 | 0.000 | TRUE |
| *DPP4* | P4 | 0.02 | 0.09 | 0.03 | 6.94 | 0.000 | 0.000 | TRUE |
| *PLXNA4* | P4 | 0.02 | 0.04 | 0.01 | 2.87 | 0.004 | 0.010 | TRUE |
| *TTYH2* | P4 | 0.02 | 0.09 | 0.05 | 6.32 | 0.000 | 0.000 | TRUE |
| *ABCA7* | P4 | 0.02 | 0.10 | 0.06 | 2.91 | 0.004 | 0.009 | TRUE |
| *PDCD1LG2* | P4 | 0.02 | 0.07 | 0.02 | 6.87 | 0.000 | 0.000 | TRUE |
| *AVPR1B* | P4 | 0.02 | 0.05 | 0.00 | 3.49 | 0.000 | 0.001 | TRUE |
| *LRRC37A2* | P4 | 0.02 | 0.11 | 0.06 | 4.23 | 0.000 | 0.000 | TRUE |
| *FZD6* | P4 | 0.02 | 0.17 | 0.12 | 2.40 | 0.016 | 0.036 | TRUE |
| *SEMA4C* | P4 | 0.02 | 0.06 | 0.02 | 7.12 | 0.000 | 0.000 | TRUE |
| *SLC41A1* | P4 | 0.02 | 0.09 | 0.04 | 7.13 | 0.000 | 0.000 | TRUE |
| *EFNB1* | P4 | 0.02 | 0.10 | 0.06 | 5.26 | 0.000 | 0.000 | TRUE |
| *SIDT1* | P4 | 0.02 | 0.07 | 0.04 | 4.31 | 0.000 | 0.000 | TRUE |
| *GPR34* | P4 | 0.02 | 0.03 | 0.01 | 2.74 | 0.006 | 0.014 | TRUE |
| *LYPD6B* | P4 | 0.02 | 0.05 | 0.00 | 5.13 | 0.000 | 0.000 | TRUE |
| *LRIG2* | P4 | 0.02 | 0.14 | 0.09 | 3.65 | 0.000 | 0.001 | TRUE |
| *NPFFR1* | P4 | 0.02 | 0.03 | 0.00 | 5.73 | 0.000 | 0.000 | TRUE |
| *MPZL2* | P4 | 0.02 | 0.06 | 0.03 | 2.59 | 0.009 | 0.022 | TRUE |
| *CEACAM19* | P4 | 0.02 | 0.05 | 0.01 | 5.30 | 0.000 | 0.000 | TRUE |
| *CACNG8* | P4 | 0.02 | 0.09 | 0.06 | 3.55 | 0.000 | 0.001 | TRUE |
| *SLC28A1* | P4 | 0.02 | 0.03 | 0.00 | 3.51 | 0.000 | 0.001 | TRUE |
| *SLC9A6* | P4 | 0.02 | 0.10 | 0.07 | 2.85 | 0.004 | 0.010 | TRUE |
| *HLA-DQB2* | P4 | 0.02 | 0.05 | 0.02 | 4.26 | 0.000 | 0.000 | TRUE |
| *GFRA2* | P4 | 0.02 | 0.03 | 0.00 | 3.51 | 0.000 | 0.001 | TRUE |
| *CDH1* | P4 | 0.02 | 0.02 | 0.00 | 3.08 | 0.002 | 0.005 | TRUE |
| *GPR55* | P4 | 0.02 | 0.03 | 0.01 | 5.95 | 0.000 | 0.000 | TRUE |
| *CLDN12* | P4 | 0.01 | 0.06 | 0.03 | 5.73 | 0.000 | 0.000 | TRUE |
| *SLC26A9* | P4 | 0.01 | 0.03 | 0.00 | 6.12 | 0.000 | 0.000 | TRUE |
| *CEACAM21* | P4 | 0.01 | 0.07 | 0.04 | 4.50 | 0.000 | 0.000 | TRUE |
| *PTGFRN* | P4 | 0.01 | 0.04 | 0.01 | 5.27 | 0.000 | 0.000 | TRUE |
| *IL18RAP* | P4 | 0.01 | 0.04 | 0.01 | 2.68 | 0.007 | 0.017 | TRUE |

| | | | | | | | | |
|---|---|---|---|---|---|---|---|---|
| *TMPRSS13* | P4 | 0.01 | 0.04 | 0.01 | 5.42 | 0.000 | 0.000 | TRUE |
| *ITGAD* | P4 | 0.01 | 0.03 | 0.00 | 3.13 | 0.002 | 0.004 | TRUE |
| *LAYN* | P4 | 0.01 | 0.04 | 0.02 | 3.99 | 0.000 | 0.000 | TRUE |
| *TNFRSF21* | P4 | 0.01 | 0.06 | 0.03 | 2.94 | 0.003 | 0.008 | TRUE |
| *TMEM231* | P4 | 0.01 | 0.06 | 0.03 | 2.85 | 0.004 | 0.011 | TRUE |
| *CNR2* | P4 | 0.01 | 0.04 | 0.02 | 3.22 | 0.001 | 0.003 | TRUE |
| *HCAR1* | P4 | 0.01 | 0.06 | 0.03 | 2.79 | 0.005 | 0.013 | TRUE |
| *TNFSF15* | P4 | 0.01 | 0.02 | 0.01 | 5.65 | 0.000 | 0.000 | TRUE |
| *AXL* | P4 | 0.01 | 0.04 | 0.01 | 3.84 | 0.000 | 0.000 | TRUE |
| *FZD4* | P4 | 0.01 | 0.02 | 0.00 | 4.31 | 0.000 | 0.000 | TRUE |
| *TMEM132A* | P4 | 0.01 | 0.03 | 0.01 | 5.67 | 0.000 | 0.000 | TRUE |
| *AJAP1* | P4 | 0.01 | 0.02 | 0.00 | 2.81 | 0.005 | 0.012 | TRUE |
| *PIGR* | P4 | 0.01 | 0.02 | 0.00 | 3.25 | 0.001 | 0.003 | TRUE |
| *EGF* | P4 | 0.01 | 0.04 | 0.02 | 2.74 | 0.006 | 0.014 | TRUE |
| *PANX2* | P4 | 0.01 | 0.17 | 0.12 | 4.64 | 0.000 | 0.000 | TRUE |
| *IGSF3* | P4 | 0.01 | 0.02 | 0.00 | 4.60 | 0.000 | 0.000 | TRUE |
| *FKRP* | P4 | 0.01 | 0.10 | 0.07 | 4.08 | 0.000 | 0.000 | TRUE |
| *ART3* | P4 | 0.01 | 0.03 | 0.01 | 3.12 | 0.002 | 0.005 | TRUE |
| *CXCR5* | P4 | 0.01 | 0.03 | 0.01 | 6.09 | 0.000 | 0.000 | TRUE |
| *PTPRS* | P4 | 0.01 | 0.04 | 0.02 | 3.02 | 0.002 | 0.006 | TRUE |
| *BTLA* | P4 | 0.01 | 0.02 | 0.00 | 3.28 | 0.001 | 0.003 | TRUE |
| *SCARF2* | P4 | 0.01 | 0.02 | 0.00 | 3.15 | 0.002 | 0.004 | TRUE |
| *CEACAM1* | P4 | 0.01 | 0.04 | 0.02 | 5.43 | 0.000 | 0.000 | TRUE |
| *BCAM* | P4 | 0.01 | 0.04 | 0.02 | 2.60 | 0.009 | 0.021 | TRUE |
| *GPR82* | P4 | 0.01 | 0.03 | 0.01 | 3.15 | 0.002 | 0.004 | TRUE |
| *PTCHD4* | P4 | 0.01 | 0.01 | 0.00 | 2.83 | 0.005 | 0.011 | TRUE |
| *SLC8A2* | P4 | 0.01 | 0.02 | 0.00 | 3.71 | 0.000 | 0.001 | TRUE |
| *ACKR3* | P4 | 0.01 | 0.02 | 0.01 | 4.78 | 0.000 | 0.000 | TRUE |
| *DRD4* | P4 | 0.01 | 0.02 | 0.00 | 2.74 | 0.006 | 0.015 | TRUE |
| *GJC2* | P4 | 0.01 | 0.02 | 0.00 | 2.45 | 0.014 | 0.031 | TRUE |
| *LSMEM1* | P4 | 0.01 | 0.05 | 0.03 | 3.12 | 0.002 | 0.005 | TRUE |
| *FCRL5* | P4 | 0.01 | 0.01 | 0.00 | 2.30 | 0.022 | 0.046 | TRUE |
| *PTPRF* | P4 | 0.01 | 0.02 | 0.00 | 3.87 | 0.000 | 0.000 | TRUE |
| *SEMA5B* | P4 | 0.01 | 0.02 | 0.00 | 2.96 | 0.003 | 0.008 | TRUE |
| *PROCR* | P4 | 0.01 | 0.06 | 0.04 | 3.00 | 0.003 | 0.007 | TRUE |
| *GALR2* | P4 | 0.01 | 0.02 | 0.00 | 3.78 | 0.000 | 0.000 | TRUE |
| *TMEM255A* | P4 | 0.01 | 0.02 | 0.00 | 5.23 | 0.000 | 0.000 | TRUE |
| *CNNM4* | P4 | 0.01 | 0.11 | 0.08 | 3.43 | 0.001 | 0.002 | TRUE |
| *SLC22A13* | P4 | 0.01 | 0.02 | 0.00 | 4.74 | 0.000 | 0.000 | TRUE |
| *CD5* | P4 | 0.01 | 0.03 | 0.02 | 3.13 | 0.002 | 0.005 | TRUE |
| *PLVAP* | P4 | 0.01 | 0.02 | 0.00 | 5.10 | 0.000 | 0.000 | TRUE |
| *EFNA5* | P4 | 0.01 | 0.02 | 0.00 | 2.96 | 0.003 | 0.007 | TRUE |
| *SLC23A1* | P4 | 0.01 | 0.02 | 0.01 | 3.01 | 0.003 | 0.006 | TRUE |
| *DUOX1* | P4 | 0.01 | 0.03 | 0.02 | 4.00 | 0.000 | 0.000 | TRUE |
| *CHRNA6* | P4 | 0.01 | 0.03 | 0.01 | 3.32 | 0.001 | 0.002 | TRUE |
| *GRIN3B* | P4 | 0.01 | 0.02 | 0.00 | 2.76 | 0.006 | 0.014 | TRUE |
| *TLR10* | P4 | 0.01 | 0.02 | 0.00 | 3.37 | 0.001 | 0.002 | TRUE |
| *HEPHL1* | P4 | 0.01 | 0.01 | 0.00 | 2.80 | 0.005 | 0.012 | TRUE |
| *EPHA2* | P4 | 0.01 | 0.01 | 0.00 | 3.83 | 0.000 | 0.000 | TRUE |
| *RTN4RL1* | P4 | 0.01 | 0.02 | 0.01 | 2.84 | 0.004 | 0.011 | TRUE |
| *OXTR* | P4 | 0.01 | 0.02 | 0.00 | 4.00 | 0.000 | 0.000 | TRUE |
| *TNFSF18* | P4 | 0.01 | 0.01 | 0.00 | 2.78 | 0.005 | 0.013 | TRUE |
| *GP5* | P4 | 0.01 | 0.01 | 0.00 | 3.12 | 0.002 | 0.005 | TRUE |
| *SLC4A5* | P4 | 0.01 | 0.04 | 0.02 | 2.92 | 0.003 | 0.008 | TRUE |
| *LRP1B* | P4 | 0.01 | 0.01 | 0.00 | 3.03 | 0.002 | 0.006 | TRUE |
| *MFAP3* | P4 | 0.01 | 0.05 | 0.04 | 2.39 | 0.017 | 0.036 | TRUE |
| *OMG* | P4 | 0.01 | 0.02 | 0.00 | 3.65 | 0.000 | 0.001 | TRUE |
| *NTNG1* | P4 | 0.01 | 0.02 | 0.00 | 3.96 | 0.000 | 0.000 | TRUE |
| *DLL1* | P4 | 0.01 | 0.03 | 0.01 | 2.39 | 0.017 | 0.037 | TRUE |
| *CDH2* | P4 | 0.01 | 0.02 | 0.00 | 2.93 | 0.003 | 0.008 | TRUE |
| *GPR153* | P4 | 0.01 | 0.01 | 0.00 | 4.45 | 0.000 | 0.000 | TRUE |
| *PARM1* | P4 | 0.01 | 0.01 | 0.00 | 2.44 | 0.015 | 0.032 | TRUE |
| *TMEFF1* | P4 | 0.01 | 0.02 | 0.01 | 3.65 | 0.000 | 0.001 | TRUE |
| *PTPRN* | P4 | 0.01 | 0.01 | 0.00 | 2.30 | 0.022 | 0.046 | TRUE |
| *CSPG4* | P4 | 0.01 | 0.01 | 0.00 | 3.73 | 0.000 | 0.001 | TRUE |
| *LHFPL4* | P4 | 0.01 | 0.03 | 0.02 | 2.35 | 0.019 | 0.041 | TRUE |
| *CLDN9* | P4 | 0.01 | 0.01 | 0.00 | 2.63 | 0.009 | 0.020 | TRUE |
| *GGT7* | P4 | 0.01 | 0.06 | 0.05 | 2.53 | 0.011 | 0.025 | TRUE |
| *SLC37A3* | P4 | 0.01 | 0.05 | 0.03 | 2.32 | 0.020 | 0.044 | TRUE |
| *ITGA3* | P4 | 0.01 | 0.03 | 0.02 | 3.49 | 0.000 | 0.001 | TRUE |
| *CD6* | P4 | 0.01 | 0.04 | 0.03 | 4.09 | 0.000 | 0.000 | TRUE |
| *ADAM11* | P4 | 0.01 | 0.02 | 0.01 | 3.78 | 0.000 | 0.000 | TRUE |
| *NOTCH3* | P4 | 0.01 | 0.02 | 0.01 | 2.84 | 0.005 | 0.011 | TRUE |
| *SLC12A3* | P4 | 0.01 | 0.01 | 0.00 | 4.09 | 0.000 | 0.000 | TRUE |
| *NLGN3* | P4 | 0.00 | 0.03 | 0.01 | 2.57 | 0.010 | 0.023 | TRUE |
| *SYP* | P4 | 0.00 | 0.03 | 0.02 | 2.61 | 0.009 | 0.021 | TRUE |
| *VASN* | P4 | 0.00 | 0.01 | 0.00 | 2.79 | 0.005 | 0.013 | TRUE |
| *CNTNAP2* | P4 | 0.00 | 0.01 | 0.00 | 2.45 | 0.014 | 0.032 | TRUE |
| *DDR1* | P4 | 0.00 | 0.01 | 0.00 | 3.88 | 0.000 | 0.000 | TRUE |
| *AOC3* | P4 | 0.00 | 0.01 | 0.00 | 2.91 | 0.004 | 0.009 | TRUE |
| *CD8A* | P4 | 0.00 | 0.01 | 0.00 | 3.75 | 0.000 | 0.001 | TRUE |
| *NLGN4Y* | P4 | 0.00 | 0.01 | 0.00 | 3.37 | 0.001 | 0.002 | TRUE |
| *CDON* | P4 | 0.00 | 0.01 | 0.00 | 4.13 | 0.000 | 0.000 | TRUE |
| *RNF43* | P4 | 0.00 | 0.06 | 0.04 | 2.36 | 0.018 | 0.039 | TRUE |
| *SLC6A16* | P4 | 0.00 | 0.01 | 0.00 | 3.09 | 0.002 | 0.005 | TRUE |
| *FAM171B* | P4 | 0.00 | 0.02 | 0.01 | 2.72 | 0.007 | 0.015 | TRUE |
| *EFNB3* | P4 | 0.00 | 0.02 | 0.01 | 2.50 | 0.012 | 0.028 | TRUE |
| *PTPRK* | P4 | 0.00 | 0.01 | 0.00 | 3.16 | 0.002 | 0.004 | TRUE |
| *SIGLEC15* | P4 | 0.00 | 0.03 | 0.02 | 2.31 | 0.021 | 0.045 | TRUE |
| *GPR155* | P4 | 0.00 | 0.03 | 0.02 | 3.24 | 0.001 | 0.003 | TRUE |
| *GPR173* | P4 | 0.00 | 0.02 | 0.01 | 2.34 | 0.019 | 0.041 | TRUE |
| *PCDH12* | P4 | 0.00 | 0.01 | 0.00 | 3.09 | 0.002 | 0.005 | TRUE |
| *SLC1A2* | P4 | 0.00 | 0.02 | 0.01 | 2.43 | 0.015 | 0.033 | TRUE |
| *UPK2* | P4 | 0.00 | 0.01 | 0.00 | 2.82 | 0.005 | 0.011 | TRUE |
| *ILDR1* | P4 | 0.00 | 0.01 | 0.00 | 2.91 | 0.004 | 0.009 | TRUE |
| *TAS1R3* | P4 | 0.00 | 0.01 | 0.00 | 2.95 | 0.003 | 0.008 | TRUE |
| *CDH13* | P4 | 0.00 | 0.01 | 0.00 | 2.44 | 0.015 | 0.032 | TRUE |
| *GPC5* | P4 | 0.00 | 0.01 | 0.00 | 2.39 | 0.017 | 0.037 | TRUE |
| *CDHR3* | P4 | 0.00 | 0.01 | 0.00 | 2.89 | 0.004 | 0.009 | TRUE |
| *CD248* | P4 | 0.00 | 0.02 | 0.01 | 3.05 | 0.002 | 0.006 | TRUE |
| *LRRTM2* | P4 | 0.00 | 0.02 | 0.01 | 2.94 | 0.003 | 0.008 | TRUE |

| | | | | | | | | |
|---|---|---|---|---|---|---|---|---|
| *SLC6A7* | P4 | 0.00 | 0.01 | 0.00 | 3.56 | 0.000 | 0.001 | TRUE |
| *GPM6B* | P4 | 0.00 | 0.05 | 0.04 | 2.69 | 0.007 | 0.017 | TRUE |
| *JAG2* | P4 | 0.00 | 0.01 | 0.00 | 2.76 | 0.006 | 0.014 | TRUE |
| *CADM1* | P4 | 0.00 | 0.01 | 0.00 | 2.32 | 0.021 | 0.044 | TRUE |
| *TRABD2A* | P4 | 0.00 | 0.02 | 0.01 | 2.92 | 0.004 | 0.009 | TRUE |
| *SEZ6* | P4 | 0.00 | 0.03 | 0.02 | 2.55 | 0.011 | 0.024 | TRUE |
| *DLK2* | P4 | 0.00 | 0.01 | 0.00 | 2.37 | 0.018 | 0.039 | TRUE |
| *TLR9* | P4 | 0.00 | 0.01 | 0.01 | 3.42 | 0.001 | 0.002 | TRUE |
| *ROR2* | P4 | 0.00 | 0.01 | 0.00 | 3.57 | 0.000 | 0.001 | TRUE |
| *PCDHGB7* | P4 | 0.00 | 0.01 | 0.00 | 3.80 | 0.000 | 0.000 | TRUE |
| *MST1R* | P4 | 0.00 | 0.01 | 0.00 | 2.65 | 0.008 | 0.019 | TRUE |
| *ATP2B2* | P4 | 0.00 | 0.01 | 0.00 | 2.62 | 0.009 | 0.020 | TRUE |
| *IGDCC4* | P4 | 0.00 | 0.01 | 0.00 | 2.58 | 0.010 | 0.023 | TRUE |
| *TIGIT* | P4 | 0.00 | 0.01 | 0.01 | 3.27 | 0.001 | 0.003 | TRUE |
| *CD27* | P4 | 0.00 | 0.01 | 0.00 | 2.68 | 0.007 | 0.017 | TRUE |
| *TGFBR3* | P4 | 0.00 | 0.01 | 0.00 | 3.34 | 0.001 | 0.002 | TRUE |
| *PDCD1* | P4 | 0.00 | 0.01 | 0.00 | 2.27 | 0.023 | 0.049 | TRUE |
| *CNTNAP1* | P4 | 0.00 | 0.01 | 0.00 | 2.96 | 0.003 | 0.008 | TRUE |
| *ABCB1* | P4 | 0.00 | 0.01 | 0.01 | 2.95 | 0.003 | 0.008 | TRUE |
| *PCDHGB6* | P4 | 0.00 | 0.01 | 0.00 | 2.93 | 0.003 | 0.008 | TRUE |
| *LIFR* | P4 | 0.00 | 0.01 | 0.00 | 2.27 | 0.023 | 0.050 | TRUE |
| *ERVW-1* | P4 | 0.00 | 0.00 | 0.00 | 2.37 | 0.018 | 0.039 | TRUE |
| *OR1I1* | P4 | 0.00 | 0.01 | 0.00 | 2.83 | 0.005 | 0.011 | TRUE |
| *SLC26A5* | P4 | 0.00 | 0.00 | 0.00 | 2.69 | 0.007 | 0.017 | TRUE |
| *CD160* | P4 | 0.00 | 0.02 | 0.02 | 2.40 | 0.016 | 0.036 | TRUE |
| *IL17RD* | P4 | 0.00 | 0.00 | 0.00 | 3.46 | 0.001 | 0.001 | TRUE |

**Table S4.** Abundance of AML programs in hematopoietic cell lines, related to **Figure 2**.

| Cell line | AML program | | | |
|---|---|---|---|---|
| | P1 | P2 | P3 | P4 |
| PL21 | 0.42 | 0.03 | 0.15 | 0.02 |
| MONOMAC1 | 0.41 | 0.04 | 0.28 | 0.10 |
| NOMO1 | 0.41 | 0.04 | 0.22 | 0.08 |
| OCIAML2 | 0.40 | 0.06 | 0.26 | 0.11 |
| SIGM5 | 0.38 | 0.10 | 0.14 | 0.02 |
| OCIAML3 | 0.37 | 0.00 | 0.13 | 0.05 |
| OCIAML5 | 0.35 | 0.04 | 0.32 | 0.08 |
| EOL1 | 0.35 | 0.03 | 0.28 | 0.08 |
| THP1 | 0.33 | 0.16 | 0.29 | 0.15 |
| SKM1 | 0.32 | 0.14 | 0.27 | 0.11 |
| MV411 | 0.31 | 0.08 | 0.20 | 0.01 |
| MOLM13 | 0.28 | 0.07 | 0.25 | 0.02 |
| PLB985 | 0.20 | 0.00 | 0.07 | 0.01 |
| HL60 | 0.11 | 0.04 | 0.08 | 0.00 |
| REH | 0.02 | 0.37 | 0.29 | 0.27 |
| SUPT1 | 0.01 | 0.34 | 0.31 | 0.30 |
| BL41 | 0.05 | 0.33 | 0.13 | 0.18 |
| ST486 | 0.01 | 0.32 | 0.26 | 0.30 |
| CI1 | 0.09 | 0.31 | 0.16 | 0.25 |
| GA10 | 0.02 | 0.30 | 0.11 | 0.22 |
| RAJI | 0.04 | 0.30 | 0.13 | 0.21 |
| P3HR1 | 0.08 | 0.27 | 0.18 | 0.26 |
| MINO | 0.10 | 0.26 | 0.23 | 0.25 |
| BL70 | 0.02 | 0.26 | 0.07 | 0.14 |
| 697 | 0.03 | 0.20 | 0.14 | 0.19 |
| M07E | 0.07 | 0.04 | 0.68 | 0.07 |
| KU812 | 0.05 | 0.08 | 0.61 | 0.07 |
| HEL | 0.09 | 0.07 | 0.61 | 0.07 |
| CMK | 0.14 | 0.05 | 0.59 | 0.08 |
| MOLM16 | 0.07 | 0.08 | 0.59 | 0.10 |
| SET2 | 0.10 | 0.07 | 0.57 | 0.14 |
| KASUMI6 | 0.13 | 0.01 | 0.55 | 0.10 |
| KG1 | 0.08 | 0.19 | 0.53 | 0.17 |
| HEL9217 | 0.05 | 0.21 | 0.52 | 0.13 |
| NCO2 | 0.06 | 0.07 | 0.50 | 0.13 |
| KARPAS422 | 0.09 | 0.08 | 0.49 | 0.34 |
| TF1 | 0.10 | 0.19 | 0.46 | 0.11 |
| MOLT16 | 0.09 | 0.04 | 0.46 | 0.30 |
| OCIM1 | 0.07 | 0.02 | 0.46 | 0.15 |
| MUTZ3 | 0.26 | 0.00 | 0.45 | 0.24 |
| KO52 | 0.09 | 0.07 | 0.45 | 0.08 |
| GDM1 | 0.10 | 0.05 | 0.44 | 0.21 |
| KCL22 | 0.09 | 0.19 | 0.44 | 0.22 |
| CMLT1 | 0.07 | 0.05 | 0.44 | 0.36 |
| P31FUJ | 0.26 | 0.13 | 0.42 | 0.20 |
| JURLMK1 | 0.12 | 0.06 | 0.42 | 0.14 |
| LP1 | 0.07 | 0.00 | 0.42 | 0.41 |
| SKNO1 | 0.02 | 0.21 | 0.42 | 0.08 |

| | | | | |
|---|---|---|---|---|
| K562 | 0.09 | 0.12 | 0.42 | 0.21 |
| LOUCY | 0.03 | 0.24 | 0.41 | 0.29 |
| MEG01 | 0.07 | 0.07 | 0.40 | 0.06 |
| RPMI8402 | 0.00 | 0.21 | 0.40 | 0.25 |
| AML193 | 0.28 | 0.16 | 0.39 | 0.09 |
| CA46 | 0.03 | 0.08 | 0.39 | 0.38 |
| HNT34 | 0.08 | 0.01 | 0.39 | 0.19 |
| MOLT13 | 0.03 | 0.21 | 0.39 | 0.31 |
| F36P | 0.12 | 0.01 | 0.38 | 0.12 |
| PEER | 0.00 | 0.27 | 0.38 | 0.14 |
| MONOMAC6 | 0.27 | 0.19 | 0.37 | 0.15 |
| SKMM2 | 0.06 | 0.07 | 0.36 | 0.21 |
| DND41 | 0.04 | 0.20 | 0.36 | 0.32 |
| A4FUK | 0.11 | 0.13 | 0.36 | 0.25 |
| JURKAT | 0.04 | 0.22 | 0.35 | 0.31 |
| P12ICHIKAWA | 0.02 | 0.27 | 0.35 | 0.26 |
| SUPT11 | 0.05 | 0.23 | 0.35 | 0.32 |
| A3KAW | 0.11 | 0.01 | 0.35 | 0.24 |
| LAMA84 | 0.12 | 0.01 | 0.34 | 0.09 |
| MOLP8 | 0.11 | 0.02 | 0.34 | 0.31 |
| OPM2 | 0.10 | 0.02 | 0.34 | 0.28 |
| EJM | 0.12 | 0.13 | 0.34 | 0.33 |
| KMS27 | 0.11 | 0.09 | 0.34 | 0.20 |
| BCP1 | 0.08 | 0.13 | 0.34 | 0.23 |
| HUT78 | 0.06 | 0.28 | 0.34 | 0.29 |
| ALLSIL | 0.02 | 0.26 | 0.33 | 0.26 |
| EM2 | 0.21 | 0.17 | 0.33 | 0.08 |
| KMS28BM | 0.07 | 0.13 | 0.32 | 0.23 |
| JJN3 | 0.09 | 0.13 | 0.32 | 0.27 |
| KASUMI1 | 0.09 | 0.09 | 0.32 | 0.11 |
| ME1 | 0.04 | 0.02 | 0.31 | 0.03 |
| U937 | 0.29 | 0.11 | 0.31 | 0.06 |
| KMS34 | 0.07 | 0.19 | 0.31 | 0.20 |
| MOLT3 | 0.02 | 0.15 | 0.30 | 0.29 |
| MOLP2 | 0.15 | 0.01 | 0.30 | 0.21 |
| NB4 | 0.18 | 0.24 | 0.30 | 0.04 |
| MOLM6 | 0.14 | 0.01 | 0.30 | 0.04 |
| JK1 | 0.11 | 0.00 | 0.29 | 0.09 |
| KHM1B | 0.08 | 0.09 | 0.29 | 0.13 |
| KMS21BM | 0.10 | 0.03 | 0.29 | 0.19 |
| KMS12BM | 0.05 | 0.20 | 0.28 | 0.18 |
| KYO1 | 0.05 | 0.04 | 0.26 | 0.04 |
| RL | 0.07 | 0.19 | 0.26 | 0.19 |
| KOPN8 | 0.13 | 0.16 | 0.25 | 0.18 |
| NCIH929 | 0.09 | 0.03 | 0.24 | 0.15 |
| AMO1 | 0.07 | 0.13 | 0.22 | 0.16 |
| BV173 | 0.01 | 0.11 | 0.19 | 0.19 |
| RPMI8226 | 0.12 | 0.11 | 0.18 | 0.16 |
| RS411 | 0.02 | 0.10 | 0.16 | 0.01 |
| OCIMY7 | 0.05 | 0.00 | 0.15 | 0.11 |
| JVM2 | 0.17 | 0.00 | 0.16 | 0.51 |
| HUT102 | 0.14 | 0.00 | 0.21 | 0.50 |

| | | | | |
|---|---|---|---|---|
| EB1 | 0.06 | 0.03 | 0.34 | 0.50 |
| HDLM2 | 0.09 | 0.09 | 0.25 | 0.49 |
| HS611T | 0.11 | 0.11 | 0.21 | 0.48 |
| EHEB | 0.16 | 0.00 | 0.12 | 0.48 |
| JVM3 | 0.16 | 0.00 | 0.14 | 0.46 |
| GRANTA519 | 0.08 | 0.07 | 0.24 | 0.45 |
| SR786 | 0.14 | 0.07 | 0.30 | 0.43 |
| L1236 | 0.11 | 0.10 | 0.27 | 0.43 |
| SUPM2 | 0.16 | 0.05 | 0.23 | 0.40 |
| L363 | 0.05 | 0.12 | 0.39 | 0.40 |
| MHHCALL4 | 0.06 | 0.14 | 0.33 | 0.40 |
| DEL | 0.18 | 0.01 | 0.24 | 0.39 |
| SUPB15 | 0.04 | 0.24 | 0.27 | 0.39 |
| KMS26 | 0.05 | 0.13 | 0.35 | 0.39 |
| MHHCALL3 | 0.05 | 0.24 | 0.27 | 0.39 |
| KMH2 | 0.10 | 0.07 | 0.13 | 0.37 |
| PF382 | 0.03 | 0.23 | 0.26 | 0.37 |
| SUDHL5 | 0.05 | 0.26 | 0.18 | 0.36 |
| MUTZ5 | 0.04 | 0.07 | 0.21 | 0.36 |
| KARPAS299 | 0.23 | 0.04 | 0.21 | 0.36 |
| TOLEDO | 0.02 | 0.32 | 0.22 | 0.36 |
| L540 | 0.12 | 0.05 | 0.23 | 0.36 |
| MJ | 0.06 | 0.23 | 0.29 | 0.36 |
| OPM1 | 0.07 | 0.05 | 0.18 | 0.36 |
| MEC1 | 0.11 | 0.07 | 0.19 | 0.35 |
| PFEIFFER | 0.03 | 0.20 | 0.30 | 0.35 |
| KASUMI2 | 0.05 | 0.17 | 0.28 | 0.35 |
| HDMYZ | 0.10 | 0.06 | 0.31 | 0.35 |
| BDCM | 0.09 | 0.01 | 0.13 | 0.34 |
| NALM1 | 0.03 | 0.11 | 0.29 | 0.34 |
| KMS20 | 0.10 | 0.20 | 0.34 | 0.34 |
| OCILY3 | 0.09 | 0.12 | 0.11 | 0.34 |
| U266B1 | 0.09 | 0.17 | 0.32 | 0.34 |
| NALM6 | 0.03 | 0.21 | 0.22 | 0.34 |
| HUNS1 | 0.05 | 0.17 | 0.15 | 0.34 |
| REC1 | 0.10 | 0.12 | 0.27 | 0.34 |
| HT | 0.07 | 0.23 | 0.28 | 0.33 |
| DAUDI | 0.08 | 0.07 | 0.17 | 0.33 |
| NALM19 | 0.03 | 0.25 | 0.31 | 0.33 |
| KMS11 | 0.09 | 0.21 | 0.29 | 0.33 |
| JM1 | 0.01 | 0.29 | 0.20 | 0.33 |
| C8166 | 0.10 | 0.12 | 0.18 | 0.32 |
| MHHCALL2 | 0.04 | 0.27 | 0.28 | 0.32 |
| KE37 | 0.03 | 0.23 | 0.29 | 0.32 |
| HPBALL | 0.03 | 0.19 | 0.26 | 0.32 |
| TALL1 | 0.01 | 0.22 | 0.30 | 0.32 |
| KMM1 | 0.04 | 0.25 | 0.30 | 0.32 |
| RI1 | 0.08 | 0.18 | 0.18 | 0.32 |
| SUDHL6 | 0.04 | 0.20 | 0.22 | 0.32 |
| INA6 | 0.08 | 0.10 | 0.29 | 0.31 |
| DB | 0.03 | 0.18 | 0.31 | 0.31 |
| SEM | 0.07 | 0.24 | 0.28 | 0.30 |

| | | | | |
|---|---|---|---|---|
| NUDUL1 | 0.07 | 0.18 | 0.18 | 0.30 |
| NAMALWA | 0.04 | 0.21 | 0.26 | 0.29 |
| KE97 | 0.06 | 0.12 | 0.16 | 0.29 |
| L428 | 0.05 | 0.12 | 0.15 | 0.29 |
| MM1S | 0.14 | 0.22 | 0.20 | 0.28 |
| HH | 0.07 | 0.21 | 0.24 | 0.28 |
| SUDHL1 | 0.10 | 0.13 | 0.26 | 0.27 |
| KARPAS620 | 0.11 | 0.14 | 0.25 | 0.27 |
| WSUDLCL2 | 0.05 | 0.06 | 0.22 | 0.26 |
| MC116 | 0.04 | 0.19 | 0.17 | 0.25 |
| OCILY19 | 0.04 | 0.05 | 0.19 | 0.24 |
| KMS18 | 0.06 | 0.02 | 0.16 | 0.23 |
| DOHH2 | 0.08 | 0.04 | 0.13 | 0.21 |
| RCHACV | 0.06 | 0.20 | 0.20 | 0.21 |
| KIJK | 0.18 | 0.10 | 0.16 | 0.21 |
| OCIMY5 | 0.07 | 0.03 | 0.19 | 0.20 |
| SUDHL8 | 0.01 | 0.19 | 0.16 | 0.20 |
| JEKO1 | 0.09 | 0.15 | 0.17 | 0.19 |
| SUDHL10 | 0.07 | 0.10 | 0.18 | 0.18 |
| NUDHL1 | 0.16 | 0.00 | 0.16 | 0.17 |
| SUDHL4 | 0.08 | 0.03 | 0.15 | 0.15 |

**Table S5.** $CD4^{IL10}$ marker genes significantly changing in $CD4^{IL10}$ cells due to co-culture with sensitive or resistant blasts, related to **Figure 3**.

**Gene symbol** - The symbol of gene for which the differential expression is assessed.

**Average log2 fold change** - The weighted average log2 fold change of gene in CD4IL10 cells after 24 hour co-culture with AML sensitive or resistant to killing (**Methods**).

**DE meta z-score** - The meta z-score obtained by aggregating individual differntial expression (DE) z-score statistics of CD4IL10 cells after 24 hour co-culture with AML sensitive or resistant to killing, using Liptak's method, across CD4IL10 donors (**Methods**).

**-log10 DE *p*-value** - The -log10 of the p-value obtained by converting the value in column **DE meta z-score** to a two-sided *p*-value. Sign indicates the dirrection of differential expression, <0 - under-expressed, >0 - over-expressed.

**-log10 DE *q*-value** - The **-lo**g1**0** *q*-value corresponding to the *p*-value in column **-log10 DE *p*-value**, after adjusted for multiple hypothesis testing using Benjamini-Hochberg method. Sign indicates the dirrection of differential expression, <0 - under-expressed, >0 - over-expressed.

**AML association** - Flag indicating wgether the gene is significantly over-expressed in CD4IL10 co-cutured with either sensitive or resistant AML.

**Suface protein** - Flag indicating whether the gene is a surface protein, as annotated in Cell Surface Protein Atlas (*Bausch-Fluck et al. 2018*).

Note: For columns B,C,F,G the the natural log of the number of CD4IL10 cells from each CD4IL10 donor before and after co-culture with each AML patient were used as weights.

| | $CD4^{IL10}$ co-cutured with sensitive AML | | | | $CD4^{IL10}$ co-cutured with resistant AML | | | | | |
|---|---|---|---|---|---|---|---|---|---|---|
| **Gene symbol** | **Average $\log_2$ fold change** | **DE meta z-score** | **$-\log_{10}$ DE *p*-value** | **$-\log_{10}$ DE *q*-value** | **Average $\log_2$ fold change** | **DE meta z-score** | **$-\log_{10}$ DE *p*-value** | **$-\log_{10}$ DE *q*-value** | **AML association** | **Surface protein** |
| IL2RA | 0.85 | 23.40 | 120.42 | 118.08 | -0.26 | -12.72 | -36.32 | -34.73 | Sensitive | TRUE |
| TNFRSF18 | 0.72 | 23.95 | 126.03 | 123.66 | -0.07 | -5.51 | -7.44 | -6.53 | Sensitive | TRUE |
| TNFRSF4 | 0.62 | 17.40 | 67.12 | 65.19 | -0.14 | -8.37 | -16.24 | -15.06 | Sensitive | TRUE |
| HAVCR2 | 0.41 | 15.70 | 54.84 | 53.03 | -0.07 | -3.37 | -3.12 | -2.47 | Sensitive | TRUE |
| SLAMF7 | 0.38 | 13.02 | 38.02 | 36.40 | 0.09 | 0.59 | 0.26 | 0.03 | Sensitive | TRUE |
| IL3RA | 0.37 | 13.31 | 39.69 | 38.05 | 0.06 | -2.31 | -1.68 | -1.17 | Sensitive | TRUE |
| CSF1 | 0.34 | 11.62 | 30.48 | 29.00 | -0.04 | -5.91 | -8.46 | -7.51 | Sensitive | TRUE |
| CD38 | 0.33 | 11.67 | 30.73 | 29.25 | -0.13 | -11.33 | -29.01 | -27.56 | Sensitive | TRUE |
| IL12RB2 | 0.30 | 14.96 | 49.90 | 48.13 | -0.03 | -5.67 | -7.83 | -6.91 | Sensitive | TRUE |
| CTLA4 | 0.29 | 14.42 | 46.40 | 44.67 | -0.01 | -4.71 | -5.60 | -4.79 | Sensitive | TRUE |
| SLAMF1 | 0.28 | 11.11 | 27.93 | 26.49 | 0.04 | -1.25 | -0.67 | -0.32 | Sensitive | TRUE |
| CD83 | 0.27 | 21.85 | 105.12 | 102.88 | 0.04 | 2.37 | 1.75 | 1.23 | Sensitive | TRUE |
| ADAM19 | 0.25 | 12.05 | 32.72 | 31.20 | -0.20 | -8.09 | -15.24 | -14.07 | Sensitive | TRUE |
| TCTN3 | 0.25 | 9.08 | 18.98 | 17.73 | 0.06 | -1.06 | -0.54 | -0.22 | Sensitive | TRUE |
| ICAM1 | 0.25 | 18.16 | 73.00 | 71.02 | 0.03 | 0.58 | 0.25 | 0.03 | Sensitive | TRUE |
| CD70 | 0.24 | 11.41 | 29.44 | 27.98 | 0.04 | -2.69 | -2.15 | -1.59 | Sensitive | TRUE |
| LRP8 | 0.19 | 11.94 | 32.12 | 30.61 | -0.02 | -4.30 | -4.77 | -4.00 | Sensitive | TRUE |
| SPPL2A | 0.18 | 6.78 | 10.92 | 9.88 | 0.04 | 0.23 | 0.09 | 0.00 | Sensitive | TRUE |
| PGAP1 | 0.18 | 10.75 | 26.24 | 24.84 | 0.05 | 0.98 | 0.49 | 0.20 | Sensitive | TRUE |
| CSF2RB | 0.18 | 6.38 | 9.75 | 8.75 | 0.00 | -3.88 | -3.99 | -3.27 | Sensitive | TRUE |
| TNFRSF8 | 0.16 | 11.74 | 31.10 | 29.60 | 0.01 | -3.73 | -3.72 | -3.02 | Sensitive | TRUE |
| SLC38A5 | 0.15 | 8.67 | 17.36 | 16.15 | 0.02 | -0.09 | -0.03 | 0.00 | Sensitive | TRUE |
| SLC7A5 | 0.14 | 7.48 | 13.12 | 12.01 | -0.03 | -5.57 | -7.60 | -6.69 | Sensitive | TRUE |
| FASLG | 0.14 | 11.69 | 30.83 | 29.34 | 0.04 | 2.37 | 1.75 | 1.23 | Sensitive | TRUE |
| ATP6V0A2 | 0.13 | 6.54 | 10.21 | 9.19 | 0.04 | -0.13 | -0.05 | 0.00 | Sensitive | TRUE |
| CD274 | 0.12 | 11.75 | 31.15 | 29.65 | 0.02 | 0.86 | 0.41 | 0.14 | Sensitive | TRUE |

| | | | | | | | | | | |
|---|---|---|---|---|---|---|---|---|---|---|
| SEMA4A | 0.12 | 13.93 | 43.36 | 41.67 | 0.02 | 2.01 | 1.35 | 0.88 | Sensitive | TRUE |
| FURIN | 0.12 | 4.85 | 5.91 | 5.07 | -0.15 | -7.73 | -13.98 | -12.85 | Sensitive | TRUE |
| NOTCH1 | 0.11 | 5.27 | 6.86 | 5.98 | 0.01 | -0.84 | -0.39 | -0.13 | Sensitive | TRUE |
| HEG1 | 0.11 | 5.83 | 8.25 | 7.31 | 0.01 | -0.54 | -0.23 | -0.03 | Sensitive | TRUE |
| PTGIR | 0.10 | 13.80 | 42.61 | 40.93 | 0.01 | 1.99 | 1.33 | 0.87 | Sensitive | TRUE |
| ECE1 | 0.10 | 5.01 | 6.27 | 5.42 | 0.04 | -0.19 | -0.07 | 0.00 | Sensitive | TRUE |
| TMEM178B | 0.09 | 5.97 | 8.63 | 7.67 | 0.02 | -1.00 | -0.50 | -0.20 | Sensitive | TRUE |
| IL15RA | 0.09 | 8.45 | 16.55 | 15.36 | 0.02 | -0.84 | -0.40 | -0.13 | Sensitive | TRUE |
| GPR137 | 0.08 | 6.68 | 10.61 | 9.58 | 0.03 | 0.37 | 0.15 | 0.00 | Sensitive | TRUE |
| NOTCH2 | 0.08 | 2.50 | 1.91 | 1.36 | -0.04 | -4.67 | -5.53 | -4.72 | Sensitive | TRUE |
| APLP2 | 0.07 | 2.69 | 2.15 | 1.59 | -0.02 | -4.60 | -5.37 | -4.57 | Sensitive | TRUE |
| IL6ST | 0.06 | 2.64 | 2.09 | 1.53 | -0.01 | -5.85 | -8.31 | -7.37 | Sensitive | TRUE |
| CCR4 | 0.05 | 6.22 | 9.30 | 8.32 | 0.00 | -2.52 | -1.94 | -1.39 | Sensitive | TRUE |
| NETO2 | 0.05 | 6.82 | 11.05 | 10.01 | -0.01 | -3.33 | -3.06 | -2.41 | Sensitive | TRUE |
| MFSD12 | 0.05 | 3.26 | 2.95 | 2.31 | 0.01 | -1.80 | -1.14 | -0.71 | Sensitive | TRUE |
| PKD2 | 0.05 | 5.31 | 6.96 | 6.08 | 0.01 | -0.59 | -0.26 | -0.03 | Sensitive | TRUE |
| ERVK13-1 | 0.05 | 2.72 | 2.19 | 1.62 | 0.03 | 1.31 | 0.72 | 0.36 | Sensitive | TRUE |
| ACP2 | 0.05 | 3.03 | 2.61 | 2.00 | 0.02 | 0.60 | 0.26 | 0.03 | Sensitive | TRUE |
| SLC9A7 | 0.05 | 5.06 | 6.39 | 5.53 | 0.00 | -2.23 | -1.59 | -1.08 | Sensitive | TRUE |
| ICAM2 | 0.05 | 4.06 | 4.30 | 3.56 | 0.03 | -0.17 | -0.06 | 0.00 | Sensitive | TRUE |
| SLC39A14 | 0.04 | 6.85 | 11.13 | 10.08 | 0.00 | -1.49 | -0.86 | -0.47 | Sensitive | TRUE |
| SLC41A2 | 0.04 | 3.23 | 2.91 | 2.27 | -0.01 | -2.89 | -2.41 | -1.82 | Sensitive | TRUE |
| MFSD2A | 0.04 | 9.06 | 18.89 | 17.64 | 0.00 | 1.48 | 0.85 | 0.47 | Sensitive | TRUE |
| CD86 | 0.04 | 7.93 | 14.65 | 13.50 | 0.00 | 0.46 | 0.19 | 0.03 | Sensitive | TRUE |
| TSPAN33 | 0.04 | 7.16 | 12.11 | 11.03 | 0.00 | -0.08 | -0.03 | 0.00 | Sensitive | TRUE |
| LAYN | 0.03 | 3.99 | 4.19 | 3.46 | 0.00 | -0.79 | -0.36 | -0.11 | Sensitive | TRUE |
| ADCY3 | 0.03 | 2.64 | 2.08 | 1.52 | -0.01 | -2.40 | -1.79 | -1.26 | Sensitive | TRUE |
| NCSTN | 0.03 | 2.84 | 2.35 | 1.77 | 0.02 | 0.78 | 0.36 | 0.11 | Sensitive | TRUE |
| IL6R | 0.03 | 7.54 | 13.33 | 12.22 | 0.00 | 1.53 | 0.90 | 0.50 | Sensitive | TRUE |
| PDCD1 | 0.03 | 4.56 | 5.30 | 4.50 | 0.02 | 2.44 | 1.83 | 1.30 | Sensitive | TRUE |
| NTRK1 | 0.03 | 5.70 | 7.92 | 6.99 | 0.01 | 1.48 | 0.86 | 0.47 | Sensitive | TRUE |
| PVR | 0.03 | 3.62 | 3.53 | 2.84 | 0.00 | -4.22 | -4.61 | -3.85 | Sensitive | TRUE |
| CD68 | 0.03 | 4.69 | 5.57 | 4.75 | 0.01 | 1.17 | 0.62 | 0.28 | Sensitive | TRUE |
| FKRP | 0.03 | 2.65 | 2.09 | 1.53 | 0.01 | -1.25 | -0.67 | -0.32 | Sensitive | TRUE |
| P2RY11 | 0.03 | 3.11 | 2.73 | 2.11 | 0.01 | 0.78 | 0.36 | 0.11 | Sensitive | TRUE |
| SEMA7A | 0.03 | 5.45 | 7.30 | 6.40 | 0.01 | 1.47 | 0.85 | 0.46 | Sensitive | TRUE |
| SLC2A6 | 0.03 | 8.20 | 15.62 | 14.45 | 0.00 | 2.31 | 1.69 | 1.17 | Sensitive | TRUE |
| SLC43A2 | 0.02 | 6.56 | 10.27 | 9.25 | 0.01 | 2.11 | 1.46 | 0.98 | Sensitive | TRUE |
| SLC30A1 | 0.02 | 3.62 | 3.52 | 2.84 | 0.01 | -0.66 | -0.29 | -0.06 | Sensitive | TRUE |
| QSOX2 | 0.02 | 2.61 | 2.04 | 1.49 | 0.00 | -0.65 | -0.29 | -0.05 | Sensitive | TRUE |
| GPR84 | 0.02 | 7.38 | 12.79 | 11.70 | 0.01 | 2.36 | 1.74 | 1.22 | Sensitive | TRUE |
| IL1RL1 | 0.02 | 4.36 | 4.88 | 4.10 | 0.00 | -0.24 | -0.09 | 0.00 | Sensitive | TRUE |
| NLGN4Y | 0.02 | 6.00 | 8.72 | 7.76 | 0.00 | 0.76 | 0.35 | 0.10 | Sensitive | TRUE |
| KCNK5 | 0.02 | 2.77 | 2.26 | 1.68 | 0.00 | -1.16 | -0.61 | -0.27 | Sensitive | TRUE |
| TFPI | 0.02 | 5.12 | 6.51 | 5.65 | 0.01 | 0.67 | 0.30 | 0.06 | Sensitive | TRUE |
| ART3 | 0.02 | 6.00 | 8.72 | 7.76 | 0.00 | 0.69 | 0.31 | 0.07 | Sensitive | TRUE |
| SLC37A1 | 0.02 | 2.53 | 1.94 | 1.39 | 0.01 | 0.55 | 0.24 | 0.03 | Sensitive | TRUE |
| SLC43A3 | 0.02 | 6.13 | 9.05 | 8.08 | 0.00 | -2.79 | -2.28 | -1.70 | Sensitive | TRUE |
| MYOF | 0.02 | 4.78 | 5.75 | 4.93 | 0.00 | -0.26 | -0.10 | 0.00 | Sensitive | TRUE |
| MRC2 | 0.02 | 3.71 | 3.68 | 2.98 | 0.00 | -0.07 | -0.03 | 0.00 | Sensitive | TRUE |
| SSTR2 | 0.01 | 3.98 | 4.16 | 3.43 | 0.00 | 2.03 | 1.37 | 0.90 | Sensitive | TRUE |
| GALR2 | 0.01 | 5.12 | 6.50 | 5.64 | 0.00 | 1.32 | 0.73 | 0.36 | Sensitive | TRUE |
| HYAL2 | 0.01 | 3.94 | 4.09 | 3.37 | -0.01 | -3.42 | -3.20 | -2.54 | Sensitive | TRUE |
| ADAM28 | 0.01 | 3.89 | 3.99 | 3.27 | 0.00 | -0.57 | -0.24 | -0.03 | Sensitive | TRUE |
| SCARB1 | 0.01 | 3.11 | 2.73 | 2.11 | 0.00 | -0.86 | -0.41 | -0.15 | Sensitive | TRUE |

| | | | | | | | | | | |
|---|---|---|---|---|---|---|---|---|---|---|
| FLVCR2 | 0.01 | 3.90 | 4.02 | 3.30 | 0.00 | 1.70 | 1.05 | 0.63 | Sensitive | TRUE |
| ITGB8 | 0.01 | 4.18 | 4.53 | 3.77 | 0.00 | -0.04 | -0.01 | 0.00 | Sensitive | TRUE |
| ATP1A4 | 0.01 | 4.81 | 5.83 | 5.00 | 0.01 | 2.12 | 1.47 | 0.98 | Sensitive | TRUE |
| IL13RA1 | 0.01 | 4.16 | 4.50 | 3.75 | 0.00 | -1.17 | -0.61 | -0.28 | Sensitive | TRUE |
| SLC14A1 | 0.01 | 3.10 | 2.72 | 2.10 | 0.00 | 2.36 | 1.74 | 1.22 | Sensitive | TRUE |
| JAG1 | 0.01 | 3.93 | 4.07 | 3.34 | 0.00 | 0.27 | 0.11 | 0.00 | Sensitive | TRUE |
| TIE1 | 0.01 | 2.85 | 2.36 | 1.77 | 0.00 | 1.50 | 0.87 | 0.48 | Sensitive | TRUE |
| CD276 | 0.01 | 3.26 | 2.95 | 2.31 | 0.00 | -2.07 | -1.41 | -0.93 | Sensitive | TRUE |
| CD200 | 0.01 | 3.10 | 2.71 | 2.09 | 0.00 | 0.08 | 0.03 | 0.00 | Sensitive | TRUE |
| SLC46A1 | 0.01 | 3.54 | 3.40 | 2.72 | 0.00 | 0.15 | 0.06 | 0.00 | Sensitive | TRUE |
| FCGR1A | 0.01 | 3.08 | 2.68 | 2.06 | 0.00 | 0.48 | 0.20 | 0.03 | Sensitive | TRUE |
| SUCNR1 | 0.01 | 4.55 | 5.27 | 4.47 | 0.00 | 1.23 | 0.66 | 0.31 | Sensitive | TRUE |
| SLCO5A1 | 0.01 | 3.71 | 3.68 | 2.98 | 0.00 | 0.69 | 0.31 | 0.07 | Sensitive | TRUE |
| ITGA2B | 0.01 | 3.68 | 3.63 | 2.94 | 0.00 | 1.79 | 1.13 | 0.69 | Sensitive | TRUE |
| TENM3 | 0.01 | 3.91 | 4.03 | 3.31 | 0.00 | 1.43 | 0.81 | 0.44 | Sensitive | TRUE |
| P2RX1 | 0.01 | 3.24 | 2.92 | 2.28 | 0.00 | 2.05 | 1.39 | 0.92 | Sensitive | TRUE |
| SLC15A3 | 0.00 | 3.13 | 2.76 | 2.14 | 0.00 | 0.00 | 0.00 | 0.00 | Sensitive | TRUE |
| CD36 | 0.00 | 2.73 | 2.20 | 1.63 | 0.00 | -0.49 | -0.20 | -0.03 | Sensitive | TRUE |
| DLL3 | 0.00 | 3.83 | 3.89 | 3.18 | 0.00 | 0.20 | 0.08 | 0.00 | Sensitive | TRUE |
| SIRPA | 0.00 | 2.94 | 2.49 | 1.89 | 0.00 | 0.91 | 0.44 | 0.17 | Sensitive | TRUE |
| NOTCH4 | 0.00 | 3.02 | 2.60 | 1.99 | 0.00 | -0.79 | -0.37 | -0.11 | Sensitive | TRUE |
| CSF3R | 0.00 | 3.44 | 3.24 | 2.58 | 0.00 | 1.20 | 0.64 | 0.30 | Sensitive | TRUE |
| MERTK | 0.00 | 2.62 | 2.06 | 1.50 | 0.00 | -0.62 | -0.27 | -0.04 | Sensitive | TRUE |
| RXFP1 | 0.00 | 2.63 | 2.07 | 1.51 | 0.00 | -0.74 | -0.34 | -0.09 | Sensitive | TRUE |
| SSPN | 0.00 | 3.01 | 2.59 | 1.98 | 0.00 | 1.21 | 0.65 | 0.30 | Sensitive | TRUE |
| HCAR3 | 0.00 | 2.61 | 2.04 | 1.49 | 0.00 | 0.00 | 0.00 | 0.00 | Sensitive | TRUE |
| DLL4 | 0.00 | 3.18 | 2.84 | 2.21 | 0.00 | 0.53 | 0.22 | 0.03 | Sensitive | TRUE |
| SLC22A4 | 0.00 | 4.01 | 4.22 | 3.49 | 0.00 | 1.68 | 1.04 | 0.61 | Sensitive | TRUE |
| CADM1 | 0.00 | 2.88 | 2.40 | 1.80 | 0.00 | 0.21 | 0.08 | 0.00 | Sensitive | TRUE |
| SLC8A3 | 0.00 | 2.99 | 2.55 | 1.95 | 0.00 | 0.00 | 0.00 | 0.00 | Sensitive | TRUE |
| MUC4 | 0.00 | 2.80 | 2.29 | 1.71 | 0.00 | 1.48 | 0.86 | 0.47 | Sensitive | TRUE |
| TMPRSS6 | 0.00 | 3.13 | 2.76 | 2.13 | 0.00 | -0.48 | -0.20 | -0.03 | Sensitive | TRUE |
| LRRTM2 | 0.00 | 2.57 | 2.00 | 1.45 | 0.00 | 0.98 | 0.48 | 0.19 | Sensitive | TRUE |
| SLC5A3 | 0.06 | -0.25 | -0.09 | 0.00 | 0.28 | 10.82 | 26.56 | 25.15 | Resistant | TRUE |
| TNFRSF14 | 0.07 | 1.02 | 0.52 | 0.21 | 0.23 | 10.87 | 26.80 | 25.38 | Resistant | TRUE |
| ITGA1 | -0.16 | -11.12 | -27.99 | -26.56 | 0.21 | 6.72 | 10.75 | 9.71 | Resistant | TRUE |
| PIK3IP1 | -0.04 | -7.73 | -13.98 | -12.85 | 0.17 | 11.09 | 27.86 | 26.43 | Resistant | TRUE |
| SPN | -0.03 | -2.16 | -1.51 | -1.02 | 0.13 | 5.94 | 8.53 | 7.58 | Resistant | TRUE |
| HLA-C | -0.21 | -2.52 | -1.93 | -1.39 | 0.10 | 8.79 | 17.82 | 16.59 | Resistant | TRUE |
| CXCR4 | -0.07 | -4.89 | -6.00 | -5.16 | 0.10 | 3.52 | 3.36 | 2.68 | Resistant | TRUE |
| CD44 | 0.06 | 0.43 | 0.18 | 0.02 | 0.10 | 4.20 | 4.58 | 3.82 | Resistant | TRUE |
| ITGB7 | -0.12 | -10.48 | -24.96 | -23.58 | 0.09 | 3.97 | 4.14 | 3.41 | Resistant | TRUE |
| P2RY8 | -0.13 | -11.42 | -29.48 | -28.02 | 0.09 | 3.46 | 3.27 | 2.60 | Resistant | TRUE |
| EMP3 | -0.44 | -12.44 | -34.78 | -33.22 | 0.09 | 6.31 | 9.57 | 8.58 | Resistant | TRUE |
| CLN3 | -0.04 | -3.58 | -3.46 | -2.78 | 0.07 | 4.61 | 5.40 | 4.60 | Resistant | TRUE |
| TRPV2 | -0.05 | -7.06 | -11.79 | -10.72 | 0.07 | 2.64 | 2.08 | 1.52 | Resistant | TRUE |
| IL10RA | -0.17 | -10.37 | -24.45 | -23.08 | 0.07 | 2.76 | 2.23 | 1.66 | Resistant | TRUE |
| ITGAL | -0.01 | 0.62 | 0.27 | 0.04 | 0.04 | 2.67 | 2.12 | 1.55 | Resistant | TRUE |
| EMB | -0.49 | -21.73 | -104.02 | -101.79 | 0.04 | 2.98 | 2.54 | 1.94 | Resistant | TRUE |
| TMEM158 | -0.01 | -5.91 | -8.46 | -7.51 | 0.03 | 4.41 | 4.98 | 4.20 | Resistant | TRUE |
| RNF43 | 0.01 | 2.30 | 1.67 | 1.16 | 0.02 | 5.28 | 6.89 | 6.01 | Resistant | TRUE |
| HLA-E | -0.19 | -8.46 | -16.58 | -15.38 | 0.02 | 2.62 | 2.05 | 1.50 | Resistant | TRUE |
| SLC52A1 | 0.00 | 1.38 | 0.77 | 0.40 | 0.01 | 4.63 | 5.44 | 4.64 | Resistant | TRUE |
| CD34 | 0.00 | 0.68 | 0.30 | 0.07 | 0.01 | 2.68 | 2.14 | 1.57 | Resistant | TRUE |
| TLR3 | 0.00 | 1.18 | 0.62 | 0.28 | 0.01 | 2.48 | 1.88 | 1.34 | Resistant | TRUE |

| | | | | | | | | | | |
|---|---|---|---|---|---|---|---|---|---|---|
| RHBDL2 | 0.00 | 2.41 | 1.79 | 1.27 | 0.00 | 3.22 | 2.89 | 2.25 | Resistant | TRUE |
| DPEP3 | 0.00 | -1.98 | -1.33 | -0.86 | 0.00 | 2.88 | 2.40 | 1.81 | Resistant | TRUE |
| CD300LF | 0.00 | 0.00 | 0.00 | 0.00 | 0.00 | 2.86 | 2.38 | 1.79 | Resistant | TRUE |
| EDA | 0.00 | 0.37 | 0.15 | 0.00 | 0.00 | 2.67 | 2.12 | 1.56 | Resistant | TRUE |
